# Self-Organizing Neural Networks in Organoids Reveal Principles of Forebrain Circuit Assembly

**DOI:** 10.1101/2025.05.01.651773

**Authors:** Sebastian Hernandez, Hunter E. Schweiger, Isabel Cline, Gregory A. Kaurala, Ash Robbins, Daniel Solis, Jinghui Geng, Tjitse van der Molen, Francisco Reyes, Chinweike Norman Asogwa, Kateryna Voitiuk, Mattia Chini, Marco Rolandi, Sofie R. Salama, Bradley M. Colquitt, Tal Sharf, David Haussler, Mircea Teodorescu, Mohammed A. Mostajo-Radji

**Affiliations:** Genomics Institute, University of California Santa Cruz, Santa Cruz, CA, 95064, United States; Department of Electrical and Computer Engineering, University of California Santa Cruz, Santa Cruz, CA, 95064, United States; Department of Molecular, Cellular and Developmental Biology, University of California Santa Cruz, Santa Cruz, CA, 95064, United States; Department of Chemistry and Biochemistry, University of California Santa Cruz, Santa Cruz, CA 95064, United States; Department of Biomolecular Engineering, University of California Santa Cruz, Santa Cruz, CA, 95064, United States; Neuroscience Research Institute, University of California Santa Barbara, Santa Barbara, CA, 93106, United States; Department of Molecular, Cellular and Developmental Biology, University of California Santa Barbara, Santa Barbara, CA, 93106, United States; Biotechnology Program, Berkeley City College, Berkeley, CA, 94704, United States; Institute of Developmental Neurophysiology, Center for Molecular Neurobiology, University Medical Center Hamburg-Eppendorf, Hamburg, Germany

## Abstract

The mouse cortex is a canonical model for studying how functional neural networks emerge, yet it remains unclear which topological features arise from intrinsic cellular organization versus external regional cues. Mouse forebrain organoids provide a powerful system to investigate these intrinsic mechanisms. We generated dorsal (DF) and ventral (VF) forebrain organoids from mouse pluripotent stem cells and tracked their development using longitudinal electrophysiology. DF organoids showed progressively stronger network-wide correlations, while VF organoids developed more refined activity patterns, enhanced small-world topology, and increased modular organization. These differences emerged without extrinsic inputs and may be driven by the increased generation of Pvalb^+^ interneurons in VF organoids. Our findings demonstrate how variations in cellular composition influence the self-organization of neural circuits, establishing mouse forebrain organoids as a tractable platform to study how neuronal populations shape cortical network architecture.

## 1 Introduction

The assembly of neural circuits during brain development requires precise coordination of molecular cues and activity-dependent refinement^1,2^. Pluripotent stem cell (PSC)-derived forebrain organoids have emerged as invaluable tools for studying neuronal development, maturation, disease mechanisms, and evolution^3–6^. Over the past decade, advancements in tissue engineering and stem cell biology have significantly improved the reproducibility of forebrain organoid generation and their long-term maintenance, particularly in human and nonhuman primate models^5–10^.

Spontaneous electrical activity arises in forebrain organoids and strengthens as they mature^11–15^. However, the extent to which this activity mirrors normal developmental processes remains a topic of debate^16,17^. A key limitation is the scarcity of primary fetal tissue for comparative studies, compounded by challenges in maintaining its viability for longitudinal functional analyses^3,9,18^. These issues impede rigorous validation of organoid fidelity to native tissue.

The emergence of electrical networks in mouse brain development is well-documented^19^. During cortical development, neurons exhibit highly synchronized patterns of spontaneous activity, dominated by correlated bursts of action potential firing that shape early network dynamics^20^. As the excitation/inhibition (E-I) ratio shifts toward inhibition, this synchronized activity transitions to sparser and less correlated firing among cortical neurons^19–22^. During postnatal maturation, the network develops two defining characteristics: (1) a small fraction of hub neurons make disproportionately many connections and strongly influence overall network activity^23^, and (2) a “small-world architecture”, characterized by dense local connectivity between neighboring neurons with sparse long range connectivity^24,25^. Current evidence suggests that these properties may emerge from intrinsic developmental programs rather than sensory experience^23^, making them ideal targets for organoid-based investigation.

PSC-derived mouse forebrain organoids were first described by the Sasai group in 2005 and subsequently refined^26,27^. While most organoid research has focused on human models^8,28–30^, mouse forebrain organoids have typically followed the GMEM-based Sasai protocol^31–33^ or used reaggregated primary neuronal progenitors^34,35^. Alternative approaches have generated unguided organoids with forebrain properties^36,37^ or limited cortical induction^38^. Recent advances using N2B27 medium enabled generation of cortical projection neurons lasting 40 days^39,40^, but protocols for electrically mature mouse forebrain organoids suitable for network-level comparisons remain needed.

Here, we established an optimized system for generating dorsal (DF) and ventral forebrain (VF) organoids from mouse PSCs. We demonstrate that these models develop distinct network architectures. DF organoids exhibit progressive synchronization, whereas VF organoids, which are enriched with Pvalb^+^ interneurons, display refined hub dynamics and stabilized connectivity. Both types form small-world networks but show different topological organization, revealing how cellular composition shapes intrinsic self-organization. This work establishes mouse forebrain organoids as a valuable model for studying the developmental principles of cortical circuit assembly and their dysregulation in disease.

## 2 RESULTS

### 2.1 A Standardized Protocol for Dorsal Forebrain Organoid Generation

Previous work from our group and others has demonstrated that GMEM-based dorsal forebrain (DF) organoids can generate neurons capable of electrophysiological maturation^31–33^. However, these neurons are often sparse and insufficient for modeling circuit-level neuronal dynamics^31–33^. To address this limitation, we optimized a robust protocol for generating DF organoids using mouse embryonic stem cells (mESCs) (Figure 1A).

**Figure 1.**
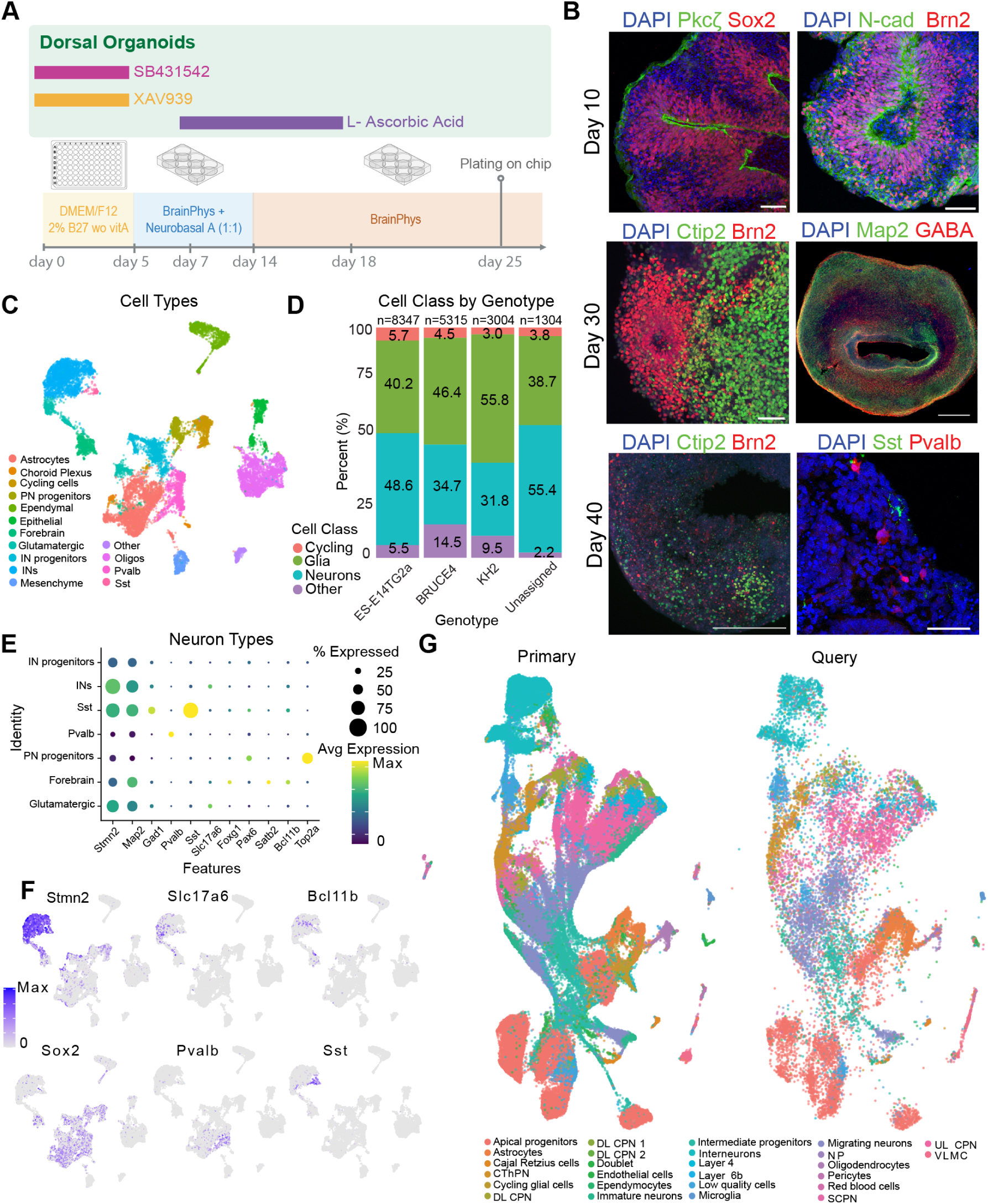
An optimized protocol for dorsal forebrain organoid development. (A) Schematic of the protocol for DF organoid development. (B) IHC of DF organoids at different time points. (Top) Day 10 DF organoids stained for Pkcζ (green, marks apical polarity in neuroepithelium) and Sox2 (red, neural progenitor marker); N-cadherin (green, marks apical adherens junctions) and Brn2 (red, upper-layer neural progenitor marker and callosal projection neuron marker). (Middle) Day 30 DF organoids stained for Ctip2 (green, marker for deep-layer corticofugal projection neurons) and Brn2 (red); Map2 (green, neuronal marker) and GABA (red, inhibitory interneuron marker). (Bottom) Day 40 DF organoids stained for Ctip2 (green, marker for deep-layer corticofugal projection neurons) and Brn2 (red); Sst (green, somatostatin-expressing interneuron marker) and Pvalb (red, parvalbumin-expressing interneuron marker). DAPI nuclear counterstain shown in blue. Scale bars: 50 or 100 µm. (C) UMAP visualization of cell types in DF organoids. INs = interneurons, PN progenitors = projection neuron progenitors. (D) Cell class distribution across three different cell lines: ES-E14TG2a, BRUCE4, and KH2. Cells that could not be confidently identified by genotype were labeled as “unassigned”. (E) Dot plot showing marker expression patterns across neuronal cell populations. (F) FeaturePlot of canonical neuronal markers: Stmn2, Slc17a6, Bcl11b, Sox2, Pvalb, and Sst. (G) Anchor-based label transfer mapping between primary tissue (developing mouse cerebral cortex) and organoid (DF organoids) datasets. DL CPN = deep layer callosal projection neuron, UL CPN = upper layer callosal projection neuron, SCPN = subcerebral projection neuron, CThPN = corticothalamic projection neuron, VLMC = vascular and leptomeningeal cells.

To establish DF organoids, we aggregated 3,000 mESCs per well in lipidure-coated V-bottom 96-well plates. After 24 hours, the resulting embryoid bodies were transitioned to forebrain differentiation medium (DMEM/F12 supplemented with N-2 and B-27 minus Vitamin A). Forebrain identity was induced by inhibiting WNT and TGF-β signaling using 5 µM XAV939 and 5 µM SB431542, with daily media changes being essential. On Day 5, organoids were transferred to ultra-low adhesion plates under continuous orbital shaking. From Days 6–14, neuronal differentiation was promoted using Neurobasal-A and BrainPhys media (1:1 ratio) supplemented with B-27 (minus Vitamin A), N-2, and 200 µM ascorbic acid to support progenitor expansion, with media refreshed every other day. By Day 15, organoids were maintained in BrainPhys medium enriched with B-27 Plus, chemically defined lipids, and heparin, while ascorbic acid was phased out by Day 25. To maintain consistency, organoid density was strictly controlled at 16 per well to ensure uniform nutrient availability (Figure 1A).

Our updated protocol led to a marked increase in Pax6 expression in DF organoids relative to our earlier GMEM-based method^33^, consistent with enhanced forebrain progenitor specification (Figure S1A-B). In addition, the new protocol reduced the proportion of off-target cell types and improved overall neuronal yield compared to the GMEM-based organoids (Figure S1C-F).

We evaluated marker expression in DF organoids using immunohistochemistry (IHC) at key developmental stages. By Day 10, DF organoids expressed progenitor markers (Sox2), exhibited axial polarity Pkcζ, and displayed extracellular matrix components of the neuroepithelium (N-cadherin) (Figure A-B). Organoids expressed the intermediate progenitor marker Tbr2, the neuronal marker Tubb3, and the dorsal forebrain markers Tbr1 and Brn2 (Figures 1B, B-D). This corresponds to mid corticogenesis, where deep-layer (Tbr1^+^) neurons have been born, and upper-layer progenitors (Brn2^+^) occupy the ventricular and subventricular zones^41–43^. Small populations of GABA^+^ interneurons were also detected (Figure E)^44,45^.

By Days 30–40, forebrain maturation was evident through the expression of the corticofugal projection neuron marker Bcl11b (also known as Ctip2) and continued Brn2 expression in postmitotic callosal projection neurons (Figure 1B)^43,46–48^. We observed the presence of Gfap^+^ astrocytes, along with GABA^+^ interneurons (Figures 1B, F-H)^49–51^. Notably, a small population of Pvalb^+^ interneurons was consistently observed, aligning with previous findings that a threedimensional environment supports their development^35,52^. Additionally, Sst^+^ interneurons were present (Figure 1B).

To systematically assess the robustness of our protocol, we performed single-cell RNA sequencing (scRNA-seq) on DF organoids derived from three genetically distinct mESC lines (Figure 1C-G): BRUCE4 (C57BL/6 background)^53^, ES-E14TG2a (129/Ola background)^54^, and KH2 (C57BL/6 × 129/Sv hybrid)^55^. Organoids were collected at Days 16, 30, and 60 to capture transcriptional dynamics across differentiation.

To minimize batch effects, cells from all three lines were pooled before sequencing and subsequently de-multiplexed by genotype. In total, we obtained single-cell transcriptomes for 17,970 cells (Day 16 = 5,696; Day 30 = 7,215; Day 60 = 5,059). Uniform manifold approximation projection (UMAP) visualization and subsequent analysis identified clusters corresponding to major cell classes (Figure S2A-B), categorized as Stmn2^+^/Map2^+^ neuronal cells, Top2a^+^ cycling progenitors, Gfap^+^/Vim^+^ glial cells, and ‘other’ if unclassified (Figure S2B-C). Further subdivision of clusters identified Slc17a6^+^ glutamatergic neurons, Ctip2^+^ and Satb2^+^ forebrain neurons, Pvalb^+^ and Sst^+^ interneurons, Pax6^+^ radial glia, and Top2a^+^ cycling cells (Figure 1C-F). Non-neuronal populations included Pdgfra^+^ oligodendrocytes, Folr1^+^ choroid plexus cells, Dcn^+^ mesenchymal cells, Krt8^+^ epithelial cells, and Krt15^+^ ependymal cells (Figure S2C).

As differentiation progressed, cellular diversity increased, yet proportional representation remained approximately consistent across all three mESC lines (Figures 1D-E and S2D-E). To further validate cellular identities, we performed anchor-based label transfer, mapping organoid transcriptomes onto a primary tissue reference UMAP^56,57^. As a reference, we used an atlas of the developing mouse cerebral cortex spanning E10.5 to postnatal day (P) 4 (Figure 1G)^3^. The organoid-derived cells successfully mapped onto the full spectrum of forebrain cell types, including neuronal progenitors, projection neurons, interneurons, and non-neuronal populations. Together, these findings indicate that our protocol reliably recapitulates forebrain specification while maintaining robustness across multiple genetic backgrounds.

### 2.2 Progressive Network Maturation in Dorsal Forebrain Organoids

To characterize the development of network activity in DF organoids, we performed longitudinal extracellular recordings using high-density multi-electrode arrays (HD-MEAs; MaxONE, Maxwell Biosystems). These arrays, equipped with 26,400 recording sites and simultaneous readout from 1,024 channels, enable network-level analysis at single-cell resolution^11–13,32,58^. Neural activity was analyzed across three developmental stages: early (days 23–33; 15 recordings with 3,678 aggregated putative neurons), intermediate (days 34–45; 55 recordings with 16,281 aggregated putative neurons), and late (days 46–64; 49 recordings with 10,037 aggregated putative neurons). We quantified network function using two key measures: firing rates, which capture individual neuronal activity (Figure 2B), and the spike-time tiling coefficient (STTC) with a window of 10ms, which reflects pairwise temporal correlations independent of firing rate^23,59^ (Figure 2C).

**Figure 2.**
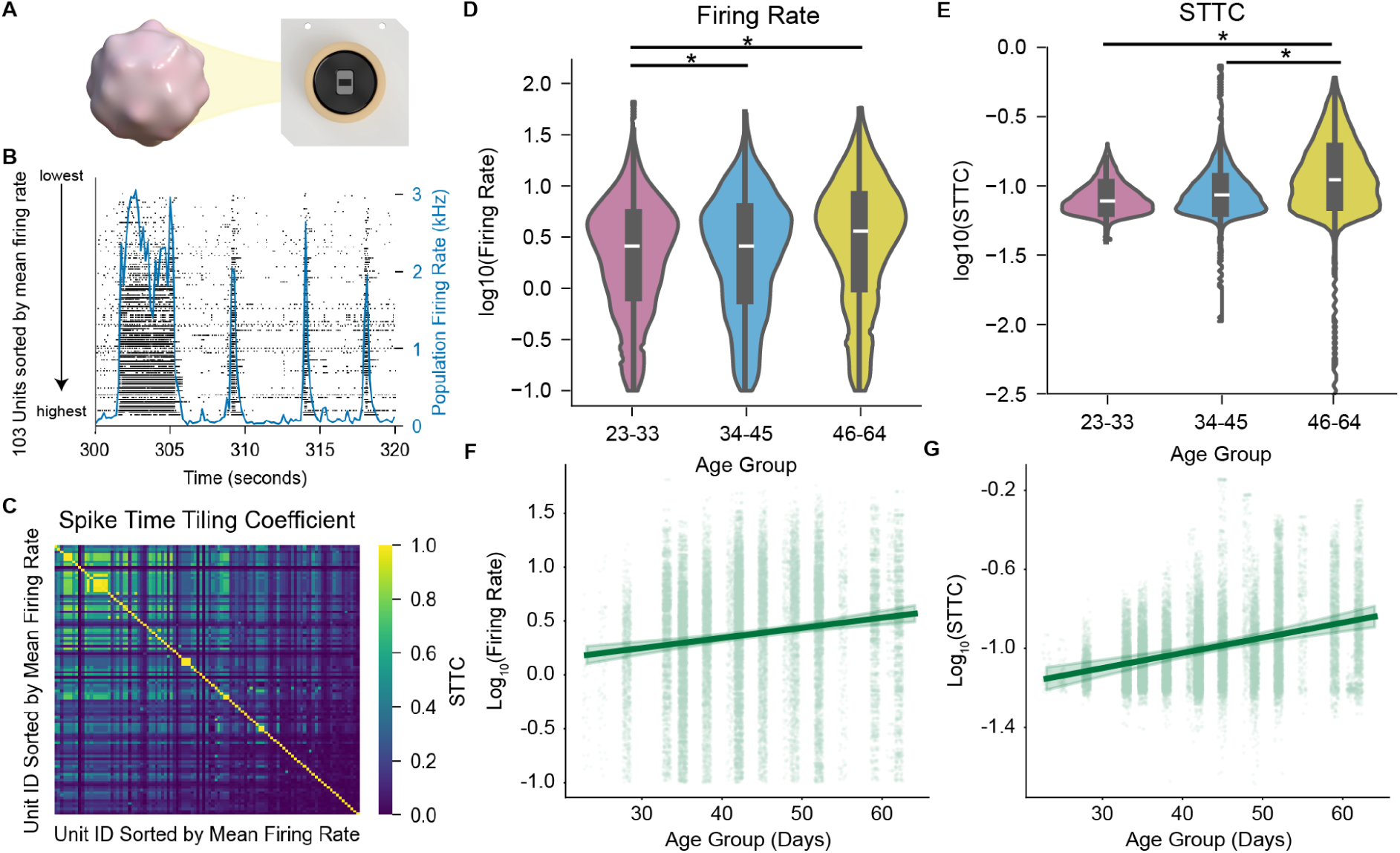
Electrophysiological characterization of dorsal forebrain organoid development. (A) Schematic of the recording setup using an HD-MEA chip. (B) Representative raster plot showing neuronal activity, with the population firing rate over time (blue). Units sorted by mean firing rate. (C) Spike time tiling coefficient (STTC) matrix showing correlation between unit spike trains, sorted by mean firing rate. (D-E) Violin plots showing log transformed mean firing rates (Hz) (D) and log transformed mean STTC (E) over early (23-33 days), mid (34-45 days), and late (46-64 days). (n = 16 organoids, 28,809 units) (F-G) Linear mixed-effects model predicted line plot of the log transformed mean firing rate distribution (F) and log transformed STTC (G). ns = not significant, * Significant after Bonferroni correction p < 0.017, Kolmogorov–Smirnov test (D-E), Mixed-effects model (F-G). Data shown as mean ± CI.

Both measures exhibited significant developmental increases. Log-transformed mean firing rates progressively rose across stages (early = 0.179 ± 0.04 Hz; intermediate = 0.38 ± 0.02 Hz; late = 0.45 ± 0.03 Hz; p < 0.001) (Table S1), consistent with prior *in vivo* observations^21,23^ (Figure 2B). Log-transformed mean STTC values also increased with age (early = -1.11 ± 0.04; intermediate = -1.02 ± 0.016; late = -0.92 ± 0.02; p < 0.001), indicating stronger spike-time correlations and progressive network synchronization (Figure 2C). Notably, this trend differs from the sparsification typically observed in the developing mouse brain and may reflect the absence of external inputs or interneuron-mediated refinement in DF organoids^21,23,60^. Despite these differences, the distributions of both measures followed log-normal distributions, consistent with fundamental electrophysiological features of neural systems^61^ (Figure S3A–D). These results underscore the utility of DF organoids as a minimalistic platform for studying principles of neural circuit maturation.

We next asked whether our differentiation protocol yields consistent electrophysiological profiles across distinct genetic backgrounds. To this end, we analyzed organoids derived from three cell lines (BRUCE4, ES-E14TG2A, KH2), as shown in Figure 1. When comparing log-transformed mean firing rates, no significant differences were detected during the early stage (BRUCE4: 0.18 ± 0.18 Hz; ES-E14TG2A: 0.12 ± 0.13 Hz; KH2: 0.30 ± 0.17 Hz; Bonferroni-corrected p > 0.17) (Figure S4A) (Table S2). In the intermediate stage, both BRUCE4 and KH2 exhibited slightly but significantly higher rates than ES-E14TG2A (BRUCE4: 0.38 ± 0.06 Hz; ES-E14TG2A: 0.24 ± 0.06 Hz; KH2: 0.43 ± 0.05 Hz; p < 0.016), while by the late stage, only BRUCE4 remained significantly different from ES-E14TG2A (0.45 ± 0.06 Hz vs. 0.33 ± 0.06 Hz; p = 0.025). However, linear mixed-effects modeling revealed no significant differences in firing rate trajectories across lines (BRUCE4: slope = 0.011, intercept = -0.07; ES-E14TG2A: slope = 0.01, intercept = -0.21; KH2: slope = 0.011, intercept = -0.03; p > 0.017) (Figure S4B). A similar pattern was observed for STTC values. In the early stage, differences between cell lines were not significant (BRUCE4: -1.11 ± 0.03; ES-E14TG2A: -1.124 ± 0.02; KH2: -1.16 ± 0.03; p > 0.17) (Figure S4C) (Table S2). In the intermediate stage, BRUCE4 exhibited higher STTC values than KH2 (-1.02 ± 0.04 vs. -1.14 ± 0.04; p = 0.002), and in the late stage, ES-E14TG2A surpassed KH2 (-0.87 ± 0.06 vs. -1.02 ± 0.06; p = 0.010). Yet, as with firing rates, developmental trajectories were similar (BRUCE4: slope = 0.006, intercept = -1.23; ES-E14TG2A: slope = 0.009, intercept = -1.35; KH2: slope = 0.005, intercept = -1.26; p > 0.05 for all comparisons) (Figure S4D).

In summary, while subtle differences in firing rate and STTC were evident at specific stages, overall developmental patterns were conserved across cell lines. These findings suggest that our protocol generates comparable electrophysiological networks regardless of genetic background.

### 2.3 Excitatory-Inhibitory Interplay Modulates Neural Dynamics in Dorsal Forebrain Organoids

The observed continual increase in DF organoids’ STTC may stem from the relatively low number of inhibitory interneurons^62^. This decrease in correlation is thought to be due to the integration and maturation of interneurons into the circuit shifting the E-I ratio towards inhibition^20,23^. To investigate how the E-I balance affects network dynamics, we pharmacologically manipulated synaptic activity.

As a control, we tested dimethyl sulfoxide (DMSO), the vehicle for drug treatments which had an insignificant effect on firing rate and STTC values (FR: baseline = 22.03 ± 1.19; DMSO = 19.63 ± 1.23; p = 0.2134) (STTC: baseline = 0.168 ± 0.014; DMSO = 0.116 ± 0.010; p = 0.5245) (Tables S3,S4) (Figure S5A). Blocking NMDA receptors with APV (2-amino-5-phosphonovaleric acid) produced no significant changes in connectivity relative to the vehicle control (STTC: baseline = 0.126 ± 0.010; APV = 0.132 ± 0.011; p = 0.4584) (Tables S3,S4) (Figure S5B). In contrast, inhibiting AMPA/Kainate receptors with NBQX (2,3-dihydroxy-6-nitro-7-sulfamoylbenzo(F)quinoxaline) significantly disrupted bursting activity and reduced network connectivity (STTC: baseline = 0.063 ± 0.010; NBQX = 0.022 ± 0.004; p = 0.0033) (Tables S3,S4) (Figure S5C). This result aligns with the established 3:1 AMPA:NMDA receptor ratio in cortical projection neurons, which accounts for the differential effects observed where AMPA/Kainate receptor inhibition substantially disrupted network connectivity while NMDA receptor blockade produced minimal impact^63^.

To examine the role of inhibition, we blocked GABA_A_ receptors with Gabazine, which artificially elevates the E-I ratio (Figures 3A-B, S5D)^12,49,64^. This treatment showed prolonged burst duration and inter-burst intervals (Figure S6). Gabazine also had a pronounced effect on network synchrony by increasing STTC values (baseline = 0.107 ± 0.011; Gabazine = 0.188 ± 0.014; p = 2.52 × 10^-5^) (Tables S3,S4), whereas firing rates remained largely unchanged (baseline = 20.81 ± 1.24; Gabazine = 23.06 ± 1.46; p = 0.789) (Tables S3,S4) (Figures 3C-D, S5D). The artificial reduction of inhibitory control underscores the key role of interneurons in structuring network activity, supporting the notion that they fine-tune connectivity patterns even in the absence of sensory input.

**Figure 3.**
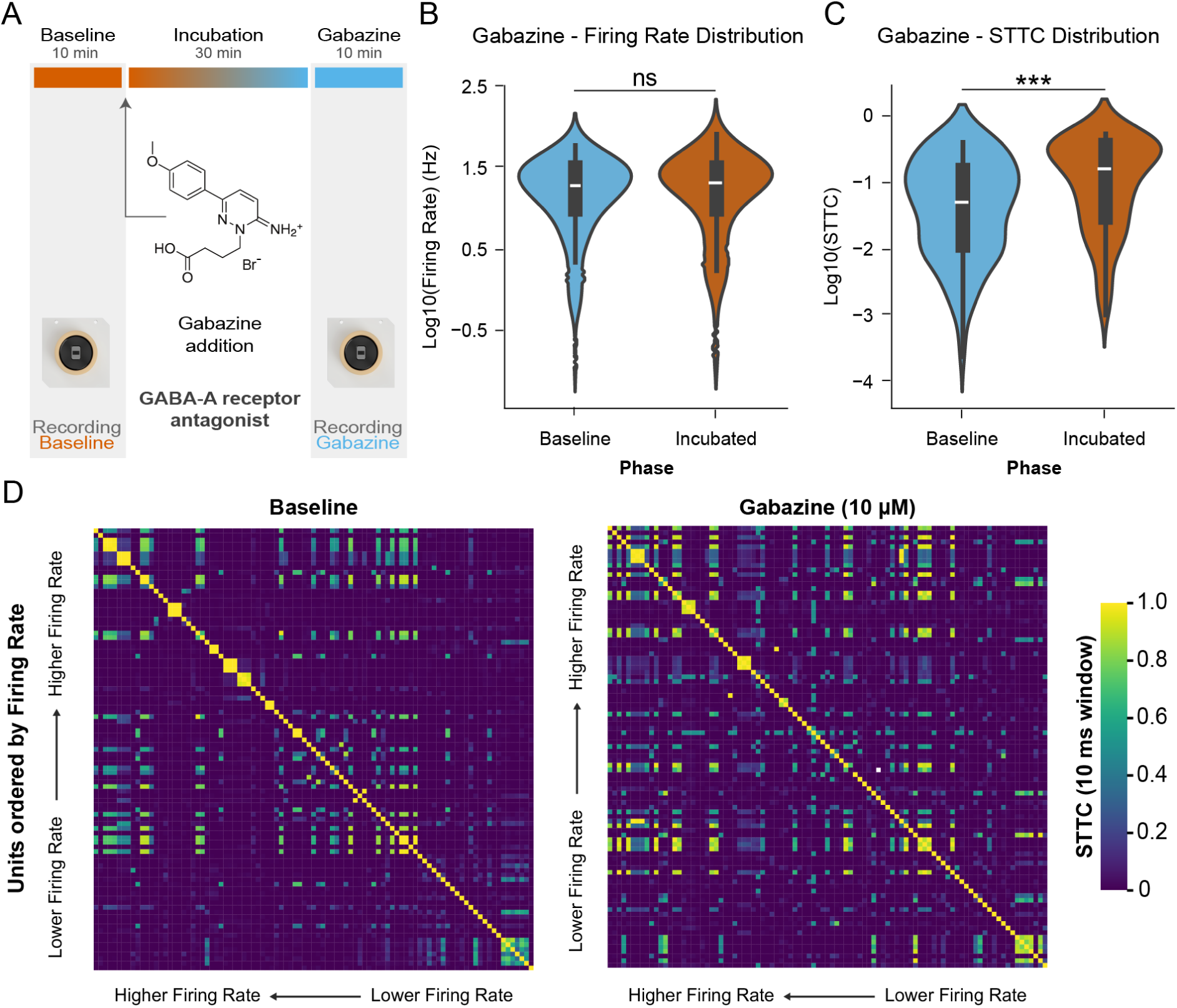
E-I balance regulates temporal coordination in dorsal forebrain organoid networks. (A) Experimental schematic of the recording protocol: 10-minute baseline recording, followed by a 30-minute drug incubation period, and a 10-minute post-incubation recording. (B-C) Violin plots showing (B) firing rates and (C) STTC distributions during baseline (blue) and after Gabazine incubation (orange). (n = 3 organoids, 133 total units). (D) STTC matrices sorted by firing rate (high to low). (Left) STTC matrix for baseline conditions. (Right) STTC matrix after Gabazine incubation. Color scale indicates STTC values from 0 to 1. ns = not significant, *p < 0.05, **p < 0.001, ***p < 0.0001, Mixed-effect models.

### 2.4 Generation and Characterization of Ventral Forebrain-Enriched Organoids

To investigate the role of inhibitory interneurons in network formation, we developed a ventral forebrain-enriched (VF) organoid model by temporally activating the Sonic Hedgehog (SHH) pathway^65–68^. Specifically, forebrain progenitors were treated with the smoothened agonist (SAG), a potent SHH activator^69^, during the first 14 days of differentiation (Figure 4A-B). This treatment led to the upregulation of the medial ganglionic eminence (MGE) progenitor marker Nkx2.1^70^ and downregulation of the dorsal forebrain progenitor marker Pax6^71^ by day 10 (Figures 4C, S7A-B and S8A-B). IHC quantification confirmed a significant shift in regional specification: Pax6 expression was enriched in DF organoids compared to VF organoids (DF = 57.78 ± 38.20%; VF = 16.64 ± 17.81%; p = 5.89×10^-9^), whereas Nkx2.1 expression was significantly higher in VF organoids (DF = 1.82 ± 2.12%; VF = 34.64 ± 20.06%; p = 3.77 × 10^-15^) (Figures 4D, S7A-B and S8A-B). VF organoids also expressed neuronal progenitor marker Sox2, the axial polarity marker Pkcζ, and the neuronal marker Tubb3 (Figure S7C-D). Furthermore, to bias the differentiation of interneurons toward a Pvalb^+^ identity, we treated the organoids with the MEK/ERK pathway inhibitor PD0325901 in conjunction with SAG^72,73^.

**Figure 4.**
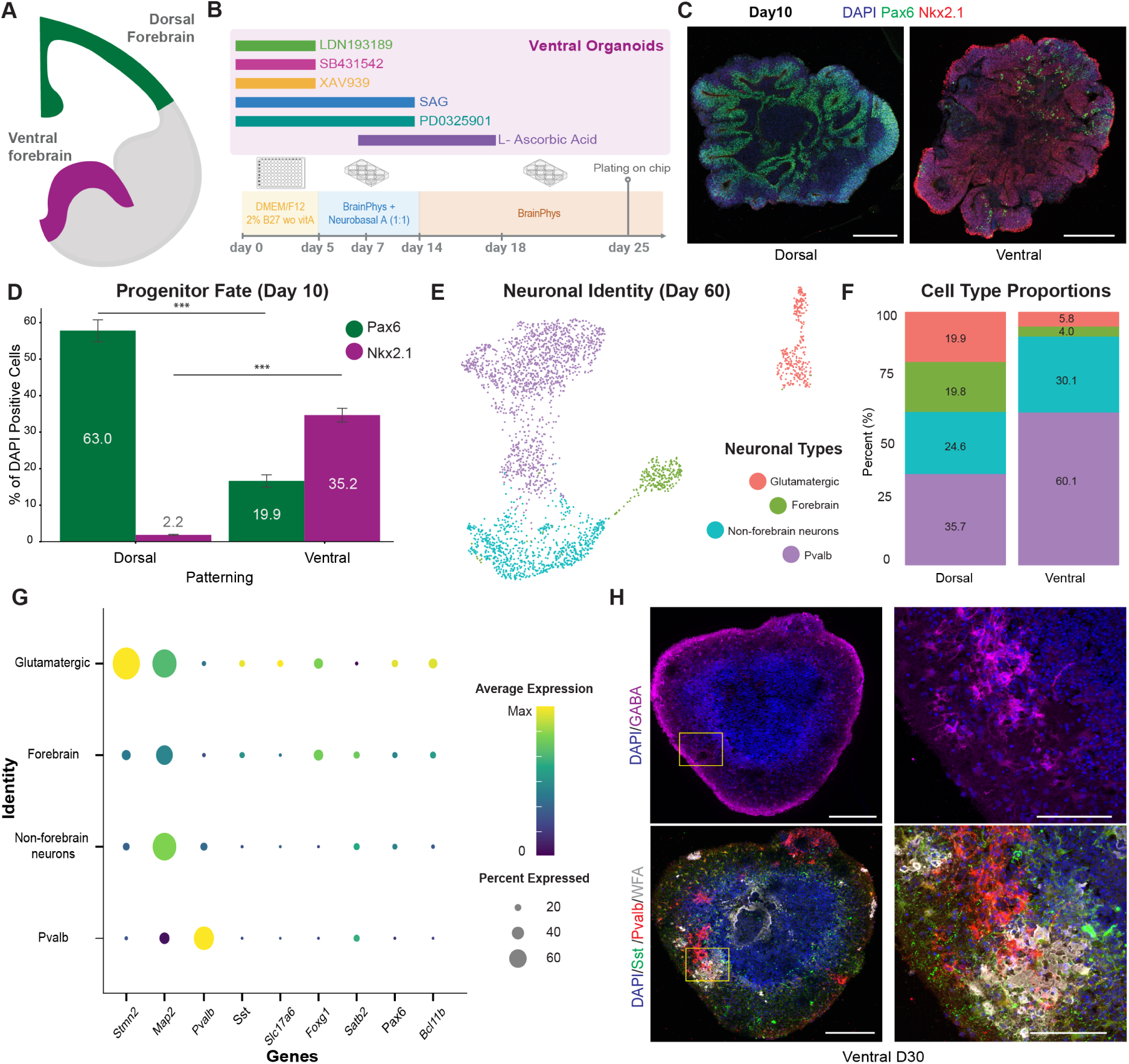
Characterization of the Braingeneers protocol for VF organoid development. (A) Schematic representation of DF (green) and VF (purple) regions. (B) Schematic of the Braingeneers protocol for VF organoid development. (C) IHC of Day 10 organoids showing DF marker Pax6 (green) and VF marker Nkx2.1 (red). DAPI nuclear counterstain shown in blue. Scale bars: 100 µm. (n = 20 organoids from 4 different batches for DF and VF each) (D) Quantification of Pax6^+^ and Nkx2.1^+^ cells across DF and VF patterned organoids. (E) UMAP visualization of neural populations identified in Day 60 single-cell RNA sequencing (scRNA-seq). (F) Cell type proportion distribution comparing DF and VF patterning. (G) Dot plot showing marker expression patterns across neuronal populations. (H) IHC of Day 30 VF organoids showing GABA (magenta), Sst (green), Pvalb (red), and WFA (gray). DAPI nuclear counterstain shown in blue. Scale bars: 100 µm and 50 µm (inset). *** p < 0.0001; Mann-Whitney U test. Data shown as mean ± SEM.

To further characterize VF organoids, we performed scRNAseq at differentiation day 60, integrating 5,059 DF and 6,111 VF cells into a unified UMAP space (Figure S8C). Cell classes were annotated based on marker genes, as shown in Figure 1, and their distributions remained consistent across the three cell lines analyzed (Figure S8D-F). Sub-setting the neuronal population and re-clustering in new UMAP space, cell classes were labeled as Glutamatergic (Slc17a6^+^), Forebrain (Satb2/Ctip2^+^), Non-forebrain (Satb2/Ctip2^-^ and Map2^+^), and Pvalb+. This revealed a distinct interneuron-enriched cluster in VF organoids, particularly within the Pvalb^+^ population (Figure 4E-G).

To validate interneuron identity, we performed IHC on serial 20 *`µ`*m cryosections, confirming robust GABA expression in the same regions as Pvalb^+^ and Sst^+^ cells (Figure 4H). We also examined perineuronal nets (PNNs), which serve as functional markers of mature Pvalb^+^ interneurons. In the mature brain, PNNs are induced by surrounding projection and inhibitory neurons, but not by Pvalb^+^ interneurons themselves^74^. Using Wisteria floribunda agglutinin (WFA) labeling^74^, we observed extensive PNN formation in Pvalb^+^ regions of VF organoids (Figure 4H). This finding is consistent with our previous work, where mouse interneuron progenitors grafted onto forebrain organoids upregulated Pvalb expression and formed PNNs^35^, further supporting the functional maturation of interneurons in VF organoids.

### 2.5 Dorsal and Ventral Forebrain Organoids Exhibit Distinct Network Dynamics

To understand the contribution of interneurons to circuit formation in organoids, we compared the electrophysiological development of VF organoids to DF organoids. First, we performed longitudinal HD-MEA recordings of the VF organoids at the same timepoints as those for the DF organoid recordings Figure 2. In VF organoids, log-transformed mean firing rates significantly increased from early to mid (p = 0.001) and early to late stages (p = 0.002), but not between mid and late development (p = 0.76). Specifically, firing rates increased from 0.10 ± 0.08 Hz (23–33 days) to 0.29 ± 0.09 Hz (34–45 days), and then plateaued at 0.27 ± 0.09 Hz (46–64 days) (Figure 5B, Table S5).

**Figure 5.**
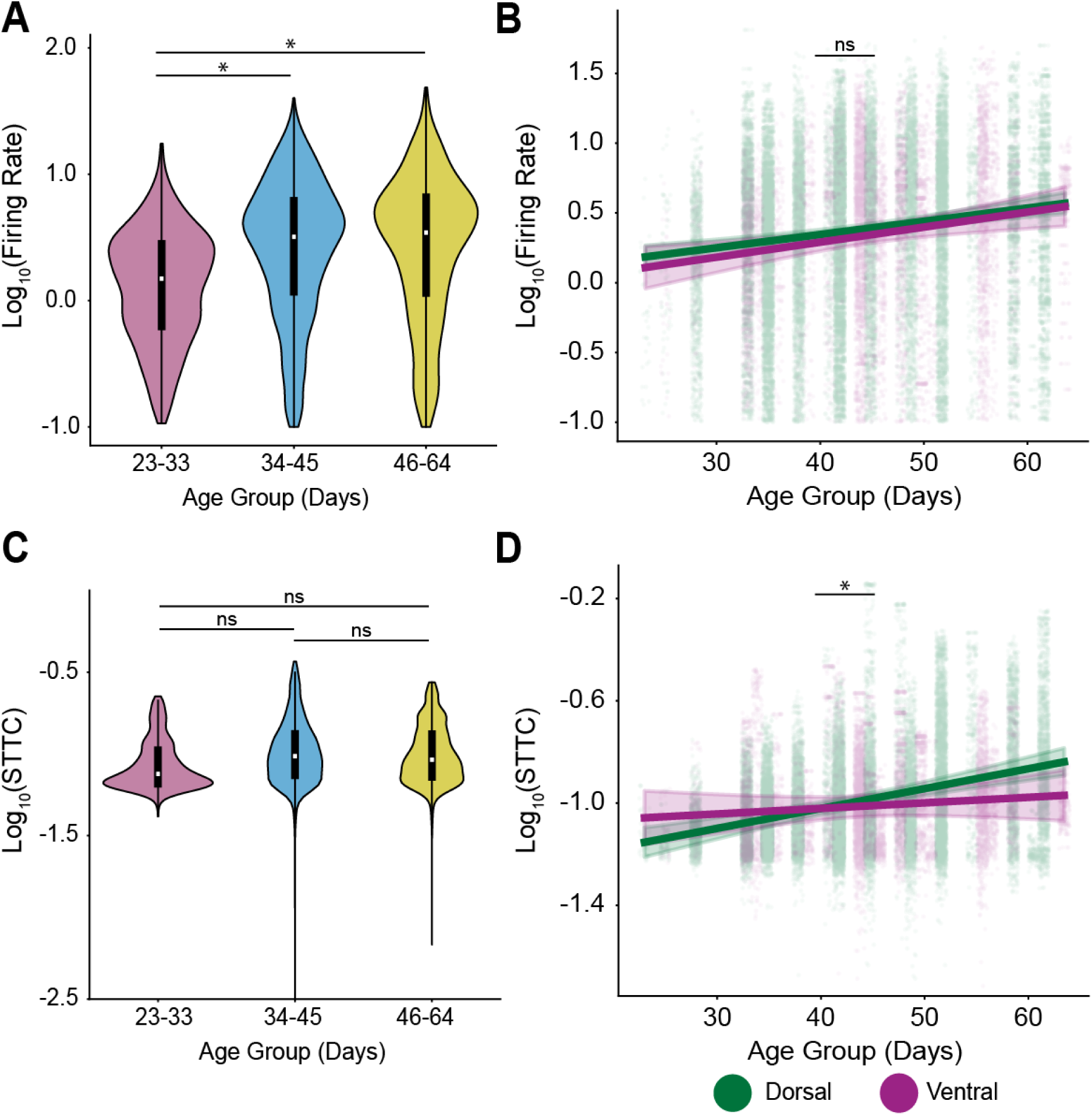
Dorsal and ventral forebrain organoids exhibit distinct developmental trajectories in neural dynamics. (A) Violin plots showing the distribution of log-transformed firing rates across three developmental stages in organoids: pink (23-33 days), blue (34-45 days), and yellow (46-64 days) (n = 18 organoids, 7,489 units). Asterisks indicate significant differences between age groups. (B) Scatter plot with regression lines (LME) showing the relationship between log-transformed firing rate (y-axis) and age in days (x-axis) for Dorsal (green) and Ventral (purple) organoids. Individual data points represent recorded units. “ns” indicates non-significant difference between the slopes of the two organoid types. (C) Violin plots displaying the distribution of log-transformed spike time tiling coefficients (STTC) across the same three developmental stages. Colors correspond to developmental stages: pink (23–33 days), blue (34–45 days), and yellow (46–64 days). “ns” indicate non-significant differences between age groups. (D) Scatter plot with regression lines illustrating the relationship between log-transformed STTC (y-axis) and age in days (x-axis) for Dorsal (green) and Ventral (purple) organoids. Statistical comparison was performed on slope. Asterisk indicates significant difference between the slopes of patterning types. *p < 0.05, **p < 0.001, ***p < 0.0001, ns = not significant, Mixed-effects model. Data shown as mean ± CI.

When comparing firing rates between VF and DF organoids at matched time points, we found no significant differences at early or mid stages. However, DF organoids displayed modest but statistically significant higher firing rates at late stages (DF: 0.45 ± 0.03 Hz; VF: 0.37 ± 0.05 Hz; p = 0.019) (Table S6). Mixed-effects modeling of age-related changes in firing rates showed no significant difference in developmental slopes (DF: 0.01 ± 0.002; VF: 0.0107 ± 0.004; p = 0.74) or intercepts (DF: -0.031 ± 0.08; VF: -0.14 ± 0.17; p = 0.54), indicating overall similar temporal dynamics between the two types of organoids (Figure 5B).

In contrast, when examining network synchrony, as measured by STTC, we found divergent developmental trajectories. STTC values in VF organoids remained relatively stable across development (Figure 5C, Table S6), whereas DF organoids exhibited a steady increase. Mixedeffects analysis confirmed a significant difference in the rate of change (slope) between DF and VF STTC values (DF: 0.008 ± 0.001; VF: 0.002 ± 0.003; p = 0.04), while intercepts were not significantly different (DF: -1.33 ± 0.06; VF: -1.10 ± 0.12; p = 0.06) (Figure 5D). These results suggest that although firing rates in VF and DF organoids follow similar patterns, their developmental progression in network synchrony diverges. Specifically, the absence of increasing STTC values in VF organoids suggests that the presence of interneurons alters how network synchrony evolves over time, leading to a different pattern of circuit refinement compared to DF organoids. However, unlike the progressive decorrelation seen *in vivo*^19,20,23^, neither organoid model displayed a continual reduction in synchrony, pointing to the likely importance of sensory input or other external factors for driving full maturation.

### 2.6 Ventral Organoids Develop Stronger Small-World Topology Through Enhanced Local Clustering

Understanding how interneurons shape hierarchical activity provides a foundation for exploring the broader network topology that emerges during organoid development. Beyond individual neuronal firing rates and correlations, network topology encompasses the overall organizational patterns that define information flow and processing efficiency^75–80^. Here, we leverage graph-theoretical approaches to examine how DF and VF organoids develop distinct network architectures and assess whether interneuron integration drives topological differences in these models.

Neural networks exhibit a spectrum of topological organization that directly impacts their information processing capabilities^80^ (Figure 6A). At one end, regular networks feature high clustering coefficients (C) and path lengths (L), creating tight-knit local connections but inefficient long-distance communication as signals must navigate through multiple intermediate nodes. At the opposite end, random networks with low values for both metrics offer shortcuts that reduce path length at the expense of coordinated local processing. Small-world networks represent a network architecture that balances local processing power with global efficiency. By maintaining high clustering coefficients while achieving short path lengths through strategic connections, these networks enable both specialized local computation and rapid information integration across distant regions. The small-world index (S) quantifies the extent to which a network exhibits these properties, calculated as the ratio of the normalized clustering coefficient to the normalized path length (S = C_norm_/L_norm_). Values significantly greater than 1 indicate a network structure that preserves local processing efficiency while ensuring rapid communication across distant regions^75,76^.

**Figure 6.**
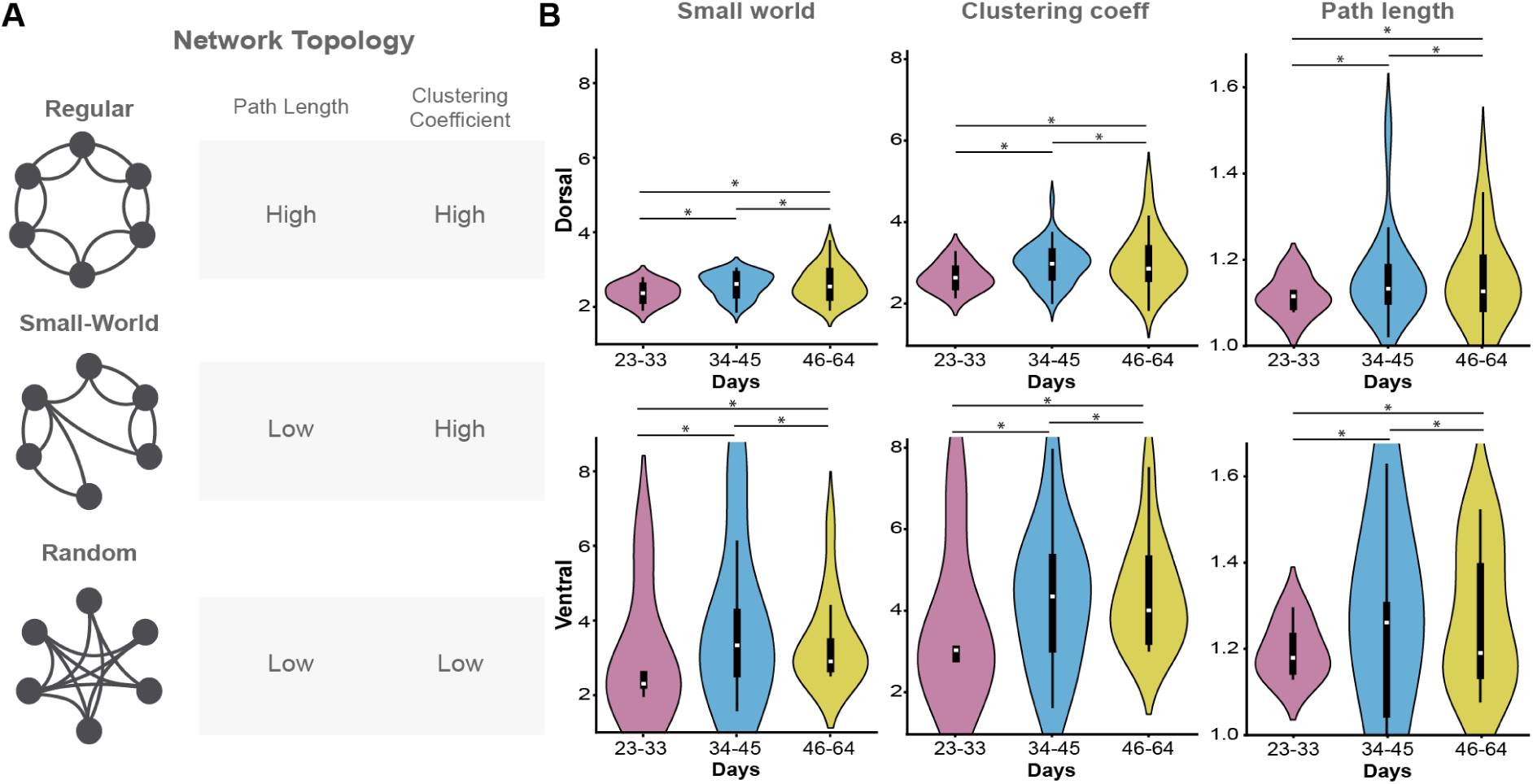
Distinct network topologies highlight organizational differences between dorsal and ventral forebrain organoids. (A) Schematic representations of different network topologies: Regular (Top), Small-World (Middle), and Random (Bottom). (B) Violin plots showing the distribution of small-world index (S) (Left) for DF (Top) and VF (Bottom), clustering coefficient (C) (Center), and path length (L) (Right), each normalized against random surrogate networks. *p < 0.0167, **p < 0.0033, ***p < 0.00033 (Bonferroni corrected), Mixed-effects model.

To evaluate network properties in our organoids and quantify their position along the topological spectrum from regular to small-world to random organization, we implemented an analytical framework based on surrogate data comparisons. For each organoid recording, we constructed a network representation by generating 1,000 surrogate datasets in which neuron IDs were shuffled while preserving mean firing rates and population activity. This approach maintained overall activity levels while disrupting temporal relationships between neurons^81,82^. STTC values exceeding the 90th percentile of the surrogate distributions were considered significant and included in the binary adjacency matrix for further analysis. These surrogates were used for all subsequent network topology characteristics^23^.

We compared S across developmental stages in both DF and VF organoids, revealing a progressive increase in small-world organization over development (Table S7, S8). During the early developmental stage (23–33 days), S values were significantly lower in DF organoids compared to VF organoids (DF mean = 2.46 ± 0.3; VF mean = 3.14 ± 1.7; p < 0.0033) (Figure 6B). This difference remained significant through the intermediate stage (34–45 days) (DF mean = 2.63 ± 0.3; VF mean = 3.30 ± 2.2; p < 0.0033) and persisted into the late developmental stage (46–64 days) (DF mean = 2.65 ± 0.4; VF mean = 3.37 pm1.2; p < 0.0033) (Figure 6B). These findings indicate that while the magnitude of regional differences remains consistent across age groups, VF organoids develop a more pronounced small-world topology over time. To further investigate the drivers of these topological differences, we analyzed L_norm_ and C_norm_ across conditions and developmental stages. Both metrics showed significant differences between DF and VF organoids at all time points (all p < 0.0033) (Table S9). However, C_norm_ emerged as the primary determinant of small-world organization, displaying a strong positive correlation with S (p = 3.57 × 10^-32^). Notably, VF organoids exhibited significantly higher C_norm_ than DF organoids, particularly during late maturation (46–64 days) (DF median = 3.37 ± 0.7; VF mean = 4.29 ± 1.3; p < 0.0033) (Table S9). This suggests that the more pronounced small-world topology observed in VF organoids is largely driven by increased local clustering, potentially reflecting enhanced interneuron-mediated connectivity.

### 2.7 Divergent Network Specialization in Dorsal and Ventral Organoids

Given the differences in small-world organization between DF and VF organoids, we next analyzed network specialization to further characterize their functional architecture. We applied *k*-core decomposition to assess hierarchical organization within the networks^83^. This iterative method identifies densely connected core regions by systematically removing nodes with fewer than k connections, beginning at k = 1. After each step, node degrees are recalculated, and the process continues until no more nodes can be pruned. The remaining subgraph at the highest k value represents the most interconnected “core” of the network, while the removed nodes constitute the “periphery”^84,85^ (Figure 7A). This approach allows us to probe the balance between centralized hubs and distributed connectivity across development.

**Figure 7.**
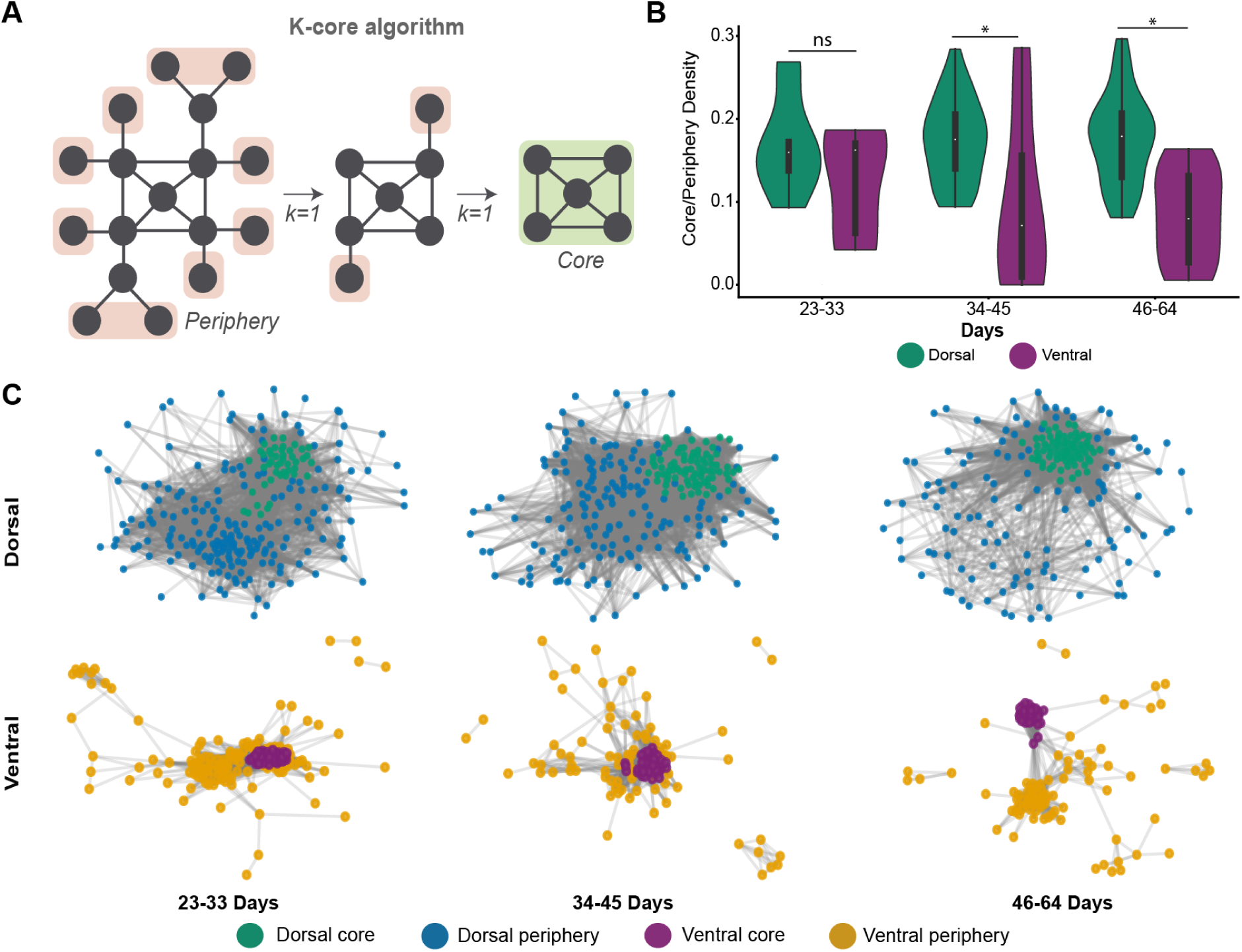
Divergent Core-Periphery Organization Reveals Distinct Network Specialization in Dorsal and Ventral Forebrain Organoids. (A) Schematic representation of the k-core algorithm used to identify core and peripheral regions within neural networks. (B) Violin plots showing core/periphery density measures across developmental stages (23–33, 34–45, and 46–64 days) for DF (green) and VF (purple) organoids. (C) Representative force-directed graph visualizations of core/periphery labeled nodes showing age group 46-64 DF (Top) core (dark green), DF periphery (blue), VF (Bottom) core (purple), and VF periphery (yellow) regions. *p < 0.05, **p < 0.001, ***p < 0.0001, Mixed-effects model

Core-periphery comparisons revealed no significant differences in functional connectivity between DF and VF organoids at early stages (days 23–33) (DF = 0.17 ± 0.02; VF = 0.13± 0.03; p = 0.54). However, by the intermediate stage, DF organoids exhibited significantly higher core-periphery interaction than VF organoids (DF = 0.17 ± 0.01; VF = 0.10 ± 0.03; p = 2.11 × 10^-4^), a difference that became more pronounced in later stages (DF = 0.18 ± 0.01; VF = 0.08 ± 0.02; p = 6.14 × 10^-5^) (Figure 7B-C). These results show a divergence in network organization: DF organoids sustain a highly integrated architecture with strong core-periphery connectivity, whereas VF organoids progressively adopt a more segregated and modular structure. This contrast suggests that dorsal networks prioritize globally integrated processing, while ventral networks increasingly rely on functionally distinct communities. Together, these findings reveal distinct organizational principles governing DF and VF networks, reflecting their divergent developmental trajectories and potential functional specializations.

### 2.8 Dorsal and Ventral Forebrain Organoids Develop Distinct Hub-Based Organization

We next examined network hubness, a key property of complex systems that highlights neurons with disproportionately high connectivity and influence over network dynamics (Figure 8A). Hub neurons have been identified *in vivo* and *in vitro* across multiple brain regions and species^86–92^. Given the distinct small-world and modular topologies of DF and VF organoids, we investigated whether these differences extend to the development and organization of hub neurons.

**Figure 8.**
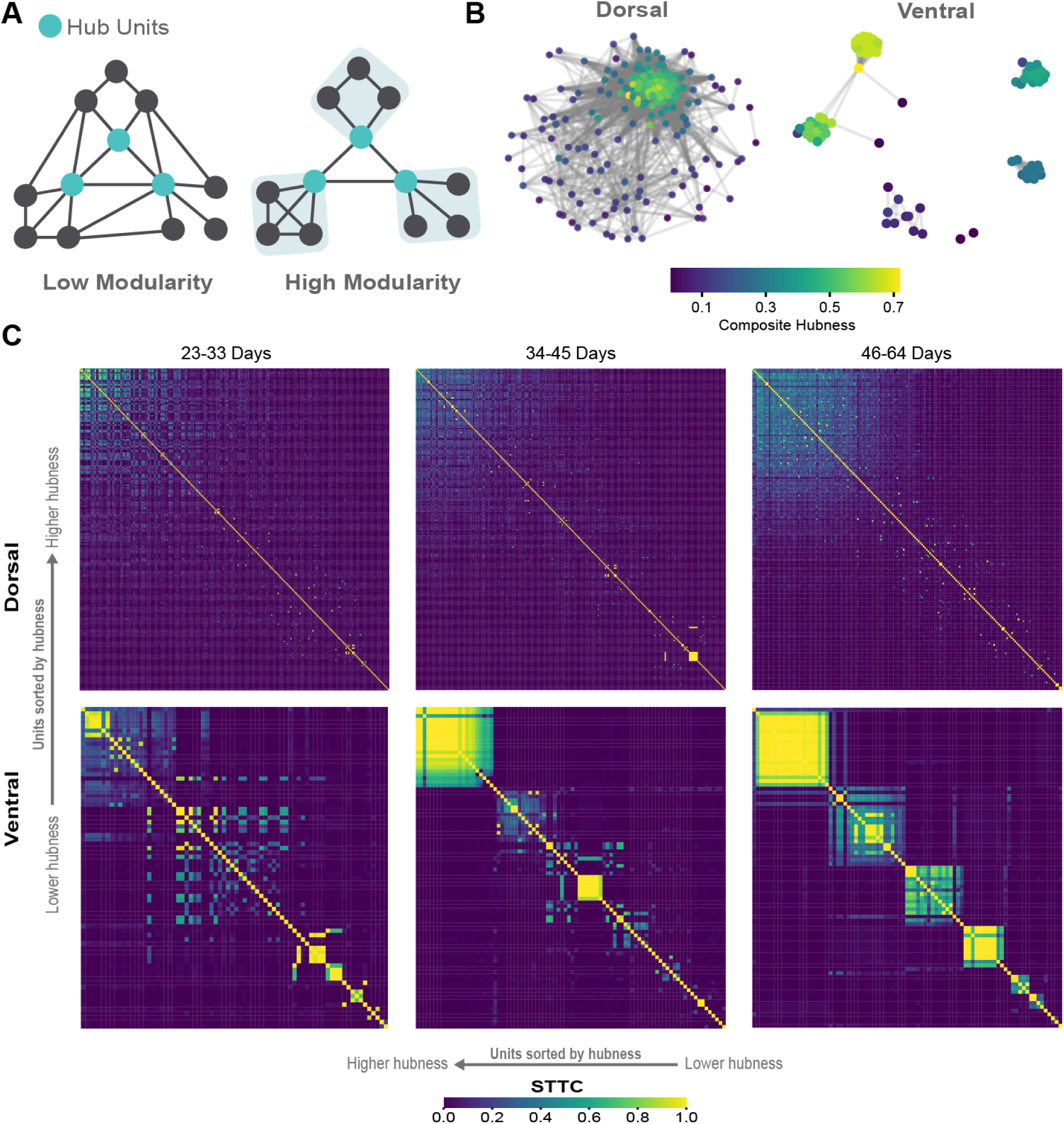
Network modularity dynamics distinguish dorsal and ventral forebrain organoid development. (A) Schematics illustrating network modularity, comparing low and high modularity states and highlighting the role of high-hub units. (B) Comparison of examples between DF and VF forebrain organoids at mature stage (46-64 days). (C) STTC matrix of units sorted by hubness score.

We calculated the composite hubness score that incorporated the node degree, node strength, betweenness, and closeness centrality^23,79^. This approach allowed us to identify neurons that not only had many connections but also occupied strategically important positions bridging network communities or enabling efficient signal propagation across the entire network. Our analysis revealed differences in hub organization between DF and VF organoids (Figure 8B, S9 S10). DF organoids formed densely interconnected networks with hub neurons distributed throughout the network core. In contrast, VF organoids developed more segregated clusters with localized hubs, exhibiting a more modular organization.

To better understand how hub units shape network topology, we sorted STTC matrices by hubness scores (Figure 8C, S9 S10). In DF organoids, highly synchronized activity was broadly distributed, consistent with an integrated network structure. In contrast, VF organoids exhibited spatially cohesive clusters of high-hubness nodes. These clusters emerged early, expanded during mid-stages, and became more spatially refined by late development, coinciding with increased modularity (23-33 days, DF = 0.226 ± 0.09, VF = 0.281 ± 0.1, p = 0.298; 34-45 days, DF = 0.273 ± 0.09, VF = 0.484 ± 0.2, p = 0.007; 46-64 days, DF = 0.262 ± 0.1, VF = 0.435 ± 0.2, p = 0.002) (Fig S11) (Table S10) and reduced core-periphery integration (Figure 7B-C). These observations suggest that hubs not only drive synchronization but also contribute to the structural compartmentalization of VF networks.

### 2.9 Distinct Core-Periphery Dynamics Underpin Developmental Specialization in Dorsal and Ventral Forebrain Organoids

To understand the functionality of the network, we examined the rigidity of bursting dynamics between DF and VF organoids. Specifically, we focused on *backbone* units, defined as neurons that spike at least twice in 90% of network bursts^12,93,94^. These backbone units are thought to form the stable core of sequential activity patterns, serving as a temporal scaffold for coordinated ensemble dynamics. Previous studies suggest that interneurons play a critical role in modulating these protosequences^12^. Our analysis revealed a significant difference in the proportion of rigid units between DF and VF organoids that was age-dependent. While no significant differences were observed in early (23-33 days: DF = 0.033 ± 0.078, VF = 0.068 ± 0.155, p = 0.8601) or intermediate stages (34-45 days: DF = 0.076 ± 0.124, VF = 0.136 ± 0.270, p = 0.1572), the late stage showed significantly higher proportion of rigid units in DF compared to VF organoids (46-64 days: DF = 0.112 ± 0.137, VF = 0.022 ± 0.035, p = 0.0003) (Figure S12A) (Table S11). This increase in rigidity suggests that DF organoids exhibit greater and more consistment unit recruitment in bursting activity.

To further investigate the organization of bursting dynamics, we applied Louvain community detection to identify functionally clustered modules^95^(Figure 9A-B). Burst events were detected within modules containing more than 10 units, a threshold chosen to reduce the likelihood of artifacts from coincidental firing among small groups of neurons. This analysis revealed that DF organoids exhibited higher burst-to-burst correlation across modules (DF = 0.239 ± 0.01; VF = 0.19 ± 0.01), indicating more stable and recurrent activation of specific neuronal ensembles (Figure 9A-B). In contrast, VF organoids showed more distributed and variable burst-to-burst correlation patterns (p = 0.001) (Figure 9C). Additionally, the temporal structure of bursting in DF organoids was more regular, as reflected by a narrower distribution in the standard deviation of burst-to-burst lag times (DF = 95.2 ± 0.9 ms; VF = 94.0 ± 1.4 ms; p = 0.019), whereas VF organoids exhibited a heterogeneous distribution, consistent with higher variance and reduced temporal precision in module recruitment (Figure 9D).

**Figure 9.**
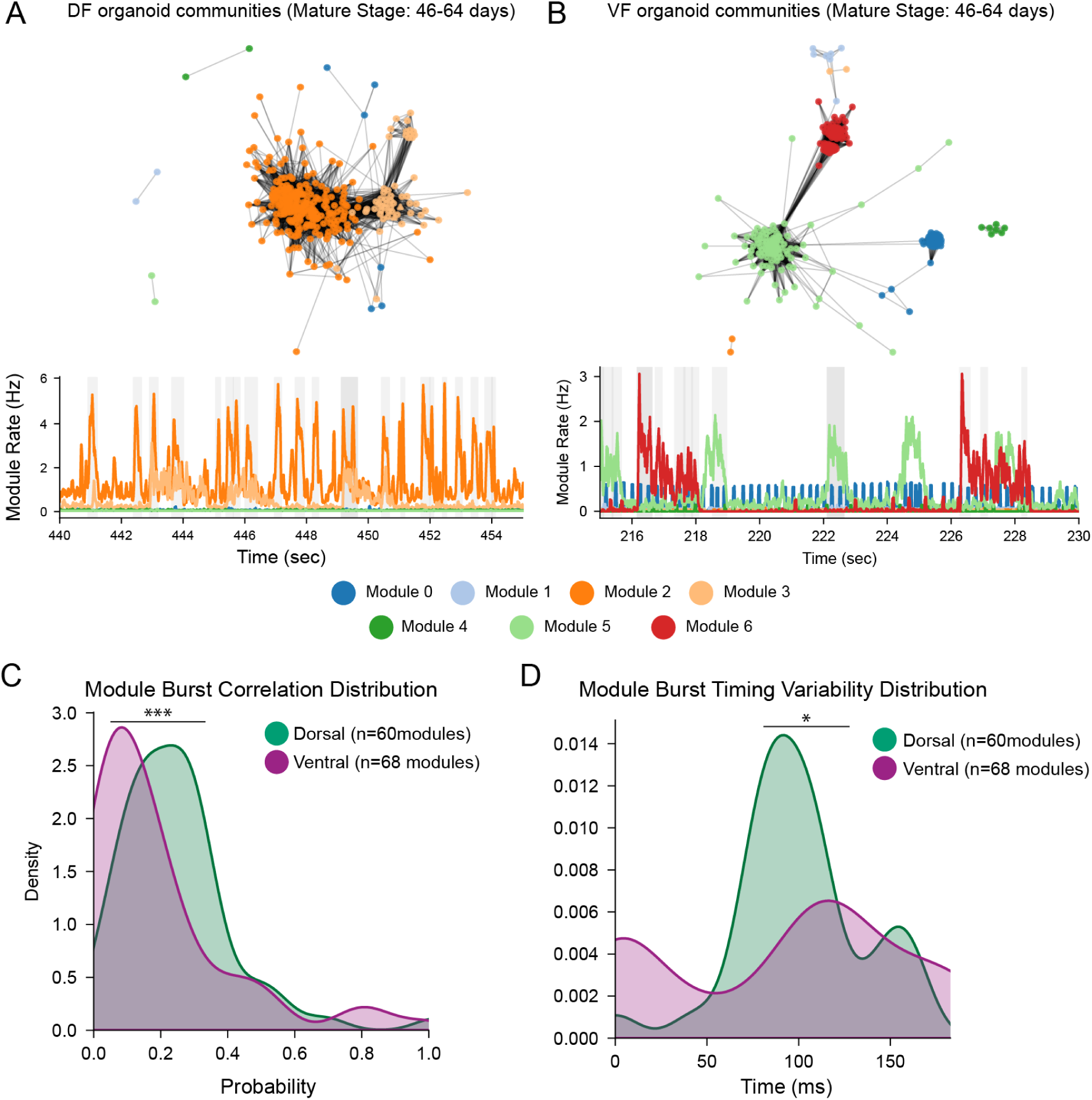
Functional community structure reveals functional differences between dorsal and ventral forebrain networks. (A) Network community structure of DF organoids at age group 46-64 showing a densely integrated organization with extensive interconnections between modules. (Top) Force-directed graph representation of STTC-derived network structure with node colors representing different modules. (Bottom) Representative time-series showing concurrent activity across modules, with Module 4 (green) and Module 6 (red) displaying highly correlated burst patterns. (B) VF organoids at the same developmental stage exhibit a more segregated community structure. (Top) Network visualization demonstrating reduced inter-module connectivity compared to DF organoids. (Bottom) Module activity patterns show distinct temporal signatures with less correlation between different functional communities. (C) Module burst correlation distribution reveals fundamental architectural differences between DF (green) and VF (purple) organoids. DF modules display higher probability of correlated bursting. (D) Module burst timing variability distribution demonstrates that VF modules (purple) exhibit broader temporal spread compared to DF modules (green), which show a narrower, more synchronized timing profile. *p < 0.05, ***p < 0.0001, Kolmogorov–Smirnov test.

These findings align with the divergent developmental trajectories identified in our coreperiphery analysis and likely reflect the complementary computational roles of excitatory and inhibitory circuits in neural processing. DF networks establish stable hierarchical structures, whereas VF networks develop flexible sub-circuits that enable context-dependent control and functional specialization.

## 3 DISCUSSION

Our study shows that mouse forebrain organoids can self-organize into physiologically relevant circuits that capture key principles of cortical network development. By optimizing protocols to generate DF and VF identities from mESCs, we systematically evaluated how cellular composition influences network dynamics. The emergence of small-world topology in both DF and VF organoids supports the idea that intrinsic developmental programs are sufficient to assemble complex network architectures, even in the absence of sensory input^23–25^. These findings establish forebrain organoids as a robust model to study how cortical circuits emerge from self-organizing developmental rules^15^.

Our results reveal that regional identity plays a central role in shaping both the dynamics and architecture of developing neural networks. DF organoids, composed primarily of excitatory projection neurons, exhibit progressive increases in firing rates and synchronization, culminating in more centralized network structures. In contrast, VF organoids, enriched in Pvalb^+^ interneurons, develop refined temporal coordination and stronger modular spatial organization without substantial changes in spike-time correlations over time. These differences highlight how projection neurons and inhibitory interneurons contribute in distinct ways to circuit refinement^19,22,23,96^. The emergence of hubs and spatial clustering in VF organoids reflects known organizational principles of Pvalb^+^ interneuron networks^97,98^, and the developmental timing and spatial features of hub formation align with the maturation trajectory of Pvalb^+^ cells^96^. Although this study did not resolve interneuron subtypes, future experiments using optogenetic, chemogenetic, or juxtacellular tagging approaches could enable selective manipulation of interneuron subclasses to define their contributions to network topology and reconfiguration^99–103^.

Despite their capacity for spontaneous self-organization, organoids do not fully recapitulate *in vivo* developmental trajectories, particularly the gradual activity decorrelation observed in the developing cortex^19,20,23^. These findings suggest that while intrinsic programs are sufficient to initiate network formation, additional external inputs, such as patterned sensory activity or longrange connections, may be required for full maturation^19,104^. Previous studies have shown that early postnatal sensory input is not essential for the emergence of several network features, including activity decorrelation, at least in the barrel cortex^20^. However, embryonic thalamic input has been demonstrated to be critical for functional specialization of the cerebral cortex^105–108^, suggesting that prenatal activity patterns may drive the formation of network topologies. Given their developmental stage, forebrain organoids could offer a platform to dissect how early activity inputs contribute to circuit assembly.

By establishing protocols for both DF and VF organoids, we provide a flexible platform for dissecting intrinsic mechanisms of cortical circuit assembly. Mouse organoids serve as a powerful complement to human models, particularly for applications that benefit from genetic precision and lineage control. While initiatives such as the MorPhic and SSPsyGene consortia are generating genome-edited human iPSC lines at scale^109–111^, the mouse research community already has access to thousands of well-characterized mESC lines. Resources such as the Mutant Mouse Resource and Research Center (MMRRC)^112^, the Texas A&M Institute for Genomic Medicine (TIGM)^113^, and the European Mouse Mutant Cell Repository (EuMMCR)^114^ offer genetically consistent lines, often derived from C57BL/6 backgrounds. This consistency enables controlled comparisons both within and across experiments, as well as between *in vitro* and *in vivo* systems.

Moreover, mouse organoids provide unique access to early stages of circuit formation. Chronic recordings in neonatal mice remain challenging due to factors such as skull fragility and maternal behavior, even with advanced platforms designed for *in vivo* use, such as Neuropixels probes^115,116^. Importantly, several neurodevelopmental disorders, including Autism spectrum disorders, schizophrenia, and Rett syndrome, are thought to arise from critical alterations in neural activity during embryonic and neonatal periods, particularly during Pvalb^+^ interneuron maturation^19,22,117–119^. Forebrain organoids offer a scalable and accessible platform to investigate how different neuronal subtypes contribute to circuit assembly and maturation during these sensitive windows. Advances in recording technologies further enhance this potential: coupling organoids with HD-MEA recordings enables high-throughput, longitudinal analysis of network activity. Notably, HD-MEAs can often be cleaned and reused across experiments, offering logistical and cost advantages over traditional *in vivo* electrophysiology platforms^120–122^. Altogether, mouse forebrain organoids represent a scalable, genetically tractable system for linking molecular perturbations to emergent circuit phenotypes, providing a valuable intermediate between genetic manipulation and behavioral outcomes.

## 4 LIMITATIONS OF THE STUDY

Several limitations should be considered when interpreting our findings. First, while organoids recapitulate core network properties, they lack key *in vivo* features including vascularization^123^ and complete cellular diversity^35,124,125^, such as Vip^+^ interneurons that can modulate network activity^126^, and microglia that have a role in synaptic prunning^127^. Structural differences may further limit their physiological relevance. Second, planar MEAs primarily sample surface neurons, potentially biasing our network analyses and hub characterizations^128^. While high-density configurations improve resolution, they cannot fully capture three-dimensional circuit organization^128^. Third, our model simplifies the complex synaptic landscape of developing circuits. We demonstrate global E-I balance effects but do not resolve subtype-specific synaptic mechanisms or short-term plasticity dynamics that shape network refinement^129^. The self-contained nature of organoids also precludes studying how sensory inputs or long-range connections influence development, despite their known importance *in vivo*^130^. These limitations define clear paths for future work: (1) incorporating additional cell types like vasculature, microglia and additional interneurons subtypes, (2) implementing 3D recording technologies to sample deeper networks, and (3) developing stimulation paradigms to study input-dependent maturation.

## 5 RESOURCE AVAILABILITY

### 5.1 Lead Contact

Further information and requests for resources and reagents should be directed to and will be fulfilled by the Lead Contact, Mohammed A. Mostajo-Radji

### 5.2 Materials availability

This study did not generate new unique reagents.

### 5.3 Data and code availability

- All scRNAseq data has been deposited in GEO under accession number GSE290330.
- All HD-MEA data has been deposited in DANDI under accession number 001374.
- All code used for plotting and analysis has been deposited at Github: https://github.com/braingeneers/Sakura_final
- Any additional information required to reanalyze the data reported in this article is available from the lead contact upon request.

## 6 ACKNOWLEDGMENTS

We thank the Colquitt lab for assistance with scRNA-seq library preparation, Kristof Tigyi for providing mESC lines, and Tomasz Nowakowski for critical feedback on this manuscript. This work was supported by Schmidt Futures (SF857), the National Human Genome Research Institute (1RM1HG011543), and the National Science Foundation (NSF2134955) to S.R.S., M.T., and D.H.; by the NSF (NSF2034037) to M.T.; by the National Institute of Mental Health (1U24MH132628) to D.H. and M.A.M.-R.; by the California Institute for Regenerative Medicine (CIRM) (DISC4-16285) to S.R.S., M.T. and M.A.M.-R., and (DISC4-16337) to M.A.M.-R.; and by the University of California Office of the President (M25PR9045) to S.R.S., M.T. and M.A.M.-R.

H.E.S. was partially supported by the NSF Graduate Research Fellowship Program (GRFP). S.H. received support from the UC Doctoral Diversity Initiative (DDI-UCSC-IBSC), and F.R. was supported by the CIRM Bridges to Stem Cell Research program.

We acknowledge the National Research Platform (NRP), supported by the National Science Foundation under award numbers CNS-1730158, ACI-1540112, ACI-1541349, and OAC-1826967, as well as by the University of California Office of the President and the UC San Diego California Institute for Telecommunications and Information Technology/Qualcomm Institute.

Sequencing was performed by the DNA Technologies and Expression Analysis Core at the UC Davis Genome Center (RRID:SCR_012480), supported by NIH Shared Instrumentation Grant 1S10OD010786-01. We also thank the UC Santa Cruz Life Sciences Microscopy Center Core Facility (RRID:SCR_021135).

## 7 AUTHOR CONTRIBUTIONS

S.H., H.E.S., M.T., and M.A.M.-R. conceptualized the project. S.H., H.E.S., I.C., G.A.K., A.R., D.S., J.G., T.v.d.M., F.R., C.A., and K.V. conducted the experiments. M.C., M.R., S.R.S., B.C., T.S., D.H., M.T., and M.A.M.-R. provided supervision and secured funding. S.H., H.E.S., and M.A.M.-R. wrote the manuscript with input from all authors.

## 9 DECLARATION OF GENERATIVE AI AND AI-ASSISTED TECHNOLOGIES

During the preparation of this work, the authors utilized ChatGPT and Claude to enhance language clarity and readability. All content was subsequently reviewed and edited as needed, and the authors take full responsibility for the final publication.

## 10 STAR METHODS

## 11 EXPERIMENTAL MODEL AND STUDY PARTICIPANT DETAILS

### 11.1 Mouse embryonic stem cell lines

We used three established mESC lines: BRUCE4 (C57BL/6 background)^53^ (RRID:CVCL_K037, Millipore Sigma # SF-CMTI-2); ES-E14TG2a (129/Ola background)^54^ (RRID:CVCL_Y481, ATCC # CRL-1821), and KH2 (C57BL/6 × 129/Sv hybrid)^55^

(RRID:CVCL_C317, Gift from Rudolf Jaenisch’s lab). All mESC lines are male. Mycoplasma testing by MycoAlert (Lonza #LT07-318) confirmed lack of contamination.

## 12 METHOD DETAILS

### 12.1 mESC Maintenance

mESCs were maintained on plates coated with 0.5 µg/mL recombinant human vitronectin (Thermo Fisher Scientific # A14700) in 1× PBS (pH 7.4; Thermo Fisher Scientific # 70011044) for 15 min at room temperature. Cells were cultured in mESC maintenance medium consisting of Glasgow Minimum Essential Medium (GMEM; Thermo Fisher Scientific # 11710035) supplemented with 10% embryonic stem cell-qualified fetal bovine serum (Thermo Fisher Scientific # 10439001), 0.1 mM MEM Non-Essential Amino Acids (Thermo Fisher Scientific # 11140050), 1 mM sodium pyruvate (Millipore Sigma # S8636), 2 mM GlutaMAX supplement (Thermo Fisher Scientific # 35050061), 0.1 mM 2-mercaptoethanol (Millipore Sigma # M3148), 0.05 mg/mL Primocin (InvivoGen # ant-pm-05), and 1000 U/mL recombinant mouse leukemia inhibitory factor (Millipore Sigma # ESG1107), with daily medium changes. Cells were passaged using ReLeSR (Stem Cell Technologies # 05872) according to manufacturer instructions and cryopreserved in mFreSR medium (Stem Cell Technologies # 05855).

### 12.2 GMEM-Based Dorsal Forebrain Protocol

Mouse forebrain organoids were generated following a modified version of a previously described protocol^27,32^. mESCs were dissociated into single cells using TrypLE Express Enzyme (Thermo Fisher Scientific # 12604021) for 5 minutes at 37°C. The cells were re-aggregated in Lipidure-coated 96-well V-bottom plates at a density of 3,000 cells per well in 100 µL of differentiation medium containing Glasgow Minimum Essential Medium (GMEM; Thermo Fisher Scientific # 11710035) supplemented with 10% Knockout Serum Replacement (Thermo Fisher Scientific # 10828028), 0.1 mM MEM Non-Essential Amino Acids (Thermo Fisher Scientific # 11140050), 1 mM Sodium Pyruvate (Millipore Sigma # S8636), 2 mM GlutaMAX supplement (Thermo Fisher Scientific # 35050061), 0.1 mM 2-Mercaptoethanol (Millipore Sigma # M3148), and 0.05 mg/mL Primocin (InvivoGen # ant-pm-05). The medium was further supplemented with 20 µM Rho kinase inhibitor Y-27632 (Tocris Bioscience # 1254), 3 µM WNT inhibitor IWR1-*`ɛ`*(Cayman Chemical # 13659), and 5 µM TGF-*`β`* inhibitor SB431542 ( Tocris Bioscience # 1614). Medium was changed daily from days 0 to 7.

On day 7, organoids were transferred to ultra-low adhesion plates (Millipore Sigma # CLS3471) containing N2 medium composed of DMEM/F12 with GlutaMAX (Thermo Fisher Scientific # 10565018), 1X N-2 Supplement (Thermo Fisher Scientific # 17502048), and 0.05 mg/mL Primocin (InvivoGen # ant-pm-05). Organoids were maintained on an orbital shaker at 60 rpm under 5% CO_2_, with medium changes every 2-3 days.

From day 14 onward, organoids were cultured in neuronal maturation medium consisting of BrainPhys Neuronal Medium (Stem Cell Technologies # 05790) supplemented with 1X N-2 Supplement (Thermo Fisher Scientific # 17502048), 1X Chemically Defined Lipid Concentrate (Thermo Fisher Scientific # 11905031), 1X B-27 Supplement (Thermo Fisher Scientific # 17504044), 0.05 mg/mL Primocin (InvivoGen # ant-pm-05), and 0.5% (v/v) Matrigel GFR Basement Membrane Matrix (LDEV-free) (Corning # 354230).

### 12.3 DMEM-based Dorsal Forebrain Protocol

mESCs were dissociated into single cells using TrypLE Express Enzyme (Thermo Fisher Scientific # 12604021) for 5 minutes at 37°C. After dissociation, the cells were re-aggregated in Lipidure-coated 96-well V-bottom plates at a density of 3,000 cells per well in 150 µL of mESC maintenance medium, supplemented with 10 µM Rho Kinase Inhibitor Y-27632 (Tocris Bioscience # 1254) and 1,000 units/mL Recombinant Mouse Leukemia Inhibitory Factor (Millipore Sigma # ESG1107). Following 24 hours of re-aggregation, the medium was replaced with forebrain patterning medium composed of DMEM/F12 with GlutaMAX (Thermo Fisher Scientific # 10565018), 10% Knockout Serum Replacement (Thermo Fisher Scientific # 10828028), 0.1 mM MEM Non-Essential Amino Acids (Thermo Fisher Scientific # 11140050), 1 mM Sodium Pyruvate (Millipore Sigma # S8636), 1X N-2 Supplement (Thermo Fisher Scientific # 17502048), 2X B-27 minus Vitamin A (Thermo Fisher Scientific # 12587010), 0.1 mM 2-Mercaptoethanol (Millipore Sigma # M3148), and 0.05 mg/mL Primocin (InvivoGen # ant-pm-05).

For dorsal forebrain patterning, the medium was further supplemented with 10 µM Rho Kinase Inhibitor Y-27632 (Tocris Bioscience # 1254), 5 µM WNT inhibitor XAV939 (StemCell Technologies # 100-1052), and 5 µM TGF-*`β`* inhibitor SB431542 (Tocris Bioscience # 1614). Medium was changed daily, with N-2 and B-27 supplements added post-filtration to preserve hydrophobic components. On day 5, organoids were transferred to ultra-low adhesion plates (Millipore Sigma # CLS3471) containing fresh neuronal differentiation medium and maintained on an orbital shaker at 68 rpm.

From days 6 to 12, progenitor expansion medium consisted of Neurobasal-A (Thermo Fisher Scientific # 10565018), BrainPhys Neuronal Medium (Stem Cell Technologies # 05790), 1X B-27 minus Vitamin A, 1X N-2 Supplement, 0.1 mM MEM Non-Essential Amino Acids, 0.05 mg/mL Primocin (InvivoGen # ant-pm-05), and 200 µM Ascorbic Acid (Sigma Aldrich # 49752). Organoids were cultured under 5% CO_2_ with medium changes every 2-3 days.

From day 15 onward, neural maturation medium contained BrainPhys Neuronal Medium supplemented with 1X B-27 Plus Supplement (Thermo Fisher Scientific, # A3582801), 1X N-2 Supplement, 1X Chemically Defined Lipid Concentrate (Thermo Fisher Scientific # 11905031), 5 µg/mL Heparin (Sigma Aldrich # H3149), and 0.05 mg/mL Primocin (InvivoGen # ant-pm-05). The medium also included 200 µM Ascorbic Acid until day 25. Medium was changed every 2-3 days with organoids maintained at 60 rpm (16 organoids per well) to minimize fusion.

### 12.4 Ventral Forebrain Protocol

Ventral forebrain organoids were generated similarly to dorsal forebrain organoids with the following modifications. The medium was supplemented with 250 nM BMP inhibitor LDN193189 (StemCell Technologies # 72147) from days 0 to 5. Additionally, from days 0 to 14, the medium contained 100 nM MEK/ERK inhibitor PD0325901 (StemCell Technologies # 72184) and 100 nM Smoothened agonist (SAG, Millipore Sigma # SIAL-SML1314).

### 12.5 Single-Cell Dissociation and Library Preparation

Mouse forebrain organoids (8-10 per genotype) were enzymatically dissociated using the Worthington Papain Dissociation System (Worthington # LK003150) following manufacturer protocols. The dissociation solution contained 20 U/mL papain, 1 mM L-cysteine, and 0.5 mM EDTA in Earle’s Balanced Salt Solution (EBSS), activated by 30 min incubation at 37°C with 200 U/mL DNase I added post-activation. Tissue samples were incubated in this solution for 30 min at 37°C with gentle agitation every 10 min, followed by mechanical dissociation using flame-polished glass Pasteur pipettes (Fisher Scientific # 13-678-6B). After centrifugation (300 RCF, 3 min), cells were resuspended in 1X PBS with 0.1% Bovine Serum Albumin (Millipore Sigma # A3311), filtered through a 40 µm cell strainer (Corning # 431750), and counted manually. For each genotype, 3,333 cells were pooled (total 10,000 cells) and processed using the PIPseq T2 Single Cell RNA v4.0PLUS platform (Fluent BioSciences # FBS-SCR-T2-8-V4.05) according to manufacturer specifications^131^.

### 12.6 Cryosection Immunohistochemistry

Organoids were fixed in 4% paraformaldehyde (Thermo Fisher Scientific # 28908), cryoprotected in 30% sucrose (Millipore Sigma # S8501), and embedded in 1:1 Tissue-Tek O.C.T. Compound (Sakura # 4583):30% sucrose. Cryosectioning (20 µm; Leica CM3050) was performed directly onto slides. After PBS washes, sections were blocked (5% donkey serum, 0.1% Triton X-100) for 1 h, incubated with primary antibodies overnight at 4°C, washed, and incubated with secondary antibodies (90 min, RT). Following final washes, sections were mounted with Fluoromount-G (Thermo Fisher Scientific # 00-4958-02).

### 12.7 Vibratome Section Immunohistochemistry

For whole-mount analysis, organoids were fixed in 4% PFA (4°C, overnight), embedded in 4% low-melt agarose (Invitrogen #16520-050), and sectioned (50 µm; Leica VT1000s vibratome). Sections underwent sequential blocking:

- Initial block: 5% donkey serum, 1% BSA, 0.5% Triton X-100 (4°C, 1 h)
- Antibody block: 2% donkey serum, 0.1% Triton X-100 (primary antibodies, overnight)

After PBS washes, sections were incubated with secondary antibodies (30 min, RT), counterstained with Hoechst 33342, and mounted with Fluoromount-G (Fisher Scientific # OB100-01).

### 12.8 Antibody Panel and Imaging

The following primary antibodies were used for immunohistochemistry, listed alphabetically by target antigen:

- Anti-Brn2 (rabbit; Thermo Fisher Scientific # PA530124, RRID:AB_2547598; 1:400)
- Anti-Ctip2 (rat; Abcam # ab18465, RRID:AB_2064130; 1:250)
- Anti-GABA (rabbit; Thermo Fisher Scientific # PA5-32241, RRID:AB_2549714; 1:375)
- Anti-GFAP (mouse; Thermo Fisher Scientific # G6171, RRID:AB_1840893; 1:100)
- Anti-Map2 (rabbit; Proteintech # 17490-1-AP, RRID:AB_2137880; 1:2000)
- Anti-N-cadherin (mouse; Abcam # ab98952, RRID:AB_10696943; 1:250)
- Anti-Nkx2.1 (rabbit; Abcam # ab76013, RRID:AB_1310784; 1:400)
- Anti-Parvalbumin (rabbit; Swant # PV27, RRID:AB_2631173; 1:375)
- Anti-Pax6 (mouse; BD Biosciences # 561462, RRID:AB_10715442; 1:100)
- Anti-PKCζ (mouse; Santa Cruz Biotechnology # sc17781, RRID:AB_628148; 1:500)
- Anti-Sox2 (mouse; Santa Cruz Biotechnology # sc365823, RRID:AB_10842165; 1:500)
- Anti-SST (mouse; Santa Cruz Biotechnology # sc-55565, RRID:AB_831726; 1:100)

Secondary detection used Alexa Fluor-conjugated antibodies (1:750) and biotinylated WFA (Vector Laboratories # B-1355-2, RRID:AB_2336874; 1:200) with Alexa 488-streptavidin (Thermo Fisher # S11223; 1:500). Nuclear counterstaining employed 300 nM DAPI (Thermo Fisher # D1306). Imaging was performed using either: Zeiss 880 Confocal Microscope with Airyscan Fast or Zeiss AxioImager Z2 Widefield Microscope, with acquisition via Zen Blue software and analysis in Zen Black/ImageJ.

### 12.9 Electrophysiological Preparation

For electrophysiological recordings, day 25 organoids were plated on MaxOne high-density multielectrode arrays (HD-MEAs; Maxwell Biosystems, # PSM). MEAs were first coated with 0.01% polyethylenimine (PEI; Millipore Sigma, # 408727) in 1× PBS for 1 h at 37°C, followed by three washes with deionized water and air-drying for 10 min. Subsequently, MEAs were coated with 20 µg/mL mouse laminin (Fisher Scientific, # CB40232) and 5 µg/mL human fibronectin (Fisher Scientific, # CB40008) in 1× PBS for 1 h at 37°C. Organoids were placed on coated MEAs, excess medium was removed, and samples were incubated at 37°C for 5–8 min to promote adhesion before adding pre-warmed neuronal differentiation medium.

### 12.10 Electrophysiological Data Processing

Electrophysiological activity was monitored every 2–3 days using Maxwell Biosystems acquisition software, sampling signals from 1024 of the ∼26,000 electrodes in a sweeping checkerboard pattern (30 s per configuration). The 1020 most active electrodes with minimum 50 µm spacing were selected for recording to ensure single-unit resolution. All recordings were performed in a humidified incubator (5% CO_2_, 37°C) at 20 kHz sampling rate and saved in HDF5 format. Raw extracellular recordings were band-pass filtered between 300–6000 Hz and spikesorted using Kilosort2^132,133^ through a custom Python pipeline. Quality control excluded units with interspike interval violation rates exceeding 0.5, mean firing rates below 0.1 Hz, or signal-to-noise ratios (SNR) below 3.

### 12.11 Pharmacological Modulation of Neuronal Activity

Dorsal forebrain (DF) organoids aged 60–65 days were scanned for spontaneous activity, with electrodes selected based on the highest activity levels following the criteria described in the Electrophysiology Data Processing section. Drug concentrations were selected based on established effective doses from previous studies^12,134^.

Following a 10-minute baseline recording, we applied the following pharmacological agents:

- Gabazine (SR95531; Abcam # ab120042) at 1 µM
- NBQX (Abcam # ab120045) at 20 µM
- APV at 100 µM

Stock solutions were prepared to enable 1:1000 dilution (1 µL per 1 mL medium), with Gabazine and NBQX dissolved in DMSO and APV in water. After drug administration, organoids were incubated for 30 minutes before acquiring 10-minute recordings of drug-modulated activity.

All recordings were processed through the following analysis pipeline:

- Concatenation using SpikeInterface^133^
- Spike sorting as described in the Electrophysiology Data Processing section
- Manual curation using Phy visualization software^135^

## 13 QUANTIFICATION AND STATISTICAL ANALYSIS

Statistical analysis was performed in Python. The statistical test, sample size, and p-value for each experiment are described in the figure legends results. Statistical significance was defined as a p-value less than 0.05 after correction for multiple comparisons when warranted.

### 13.1 Analysis of Immunohistochemistry

Organoid imaging was performed using a Zeiss AxioImager Z2 microscope with 10x magnification and Zen Blue software. For each organoid, we acquired Z-stacks at 1.53 µm spacing from three non-adjacent cryosections, with tile scanning implemented for organoids exceeding a single field of view. The analysis included 4-5 organoid replicates per cell line and protocol condition (dorsal/ventral) across two independent cell lines (ES-E14TG2a and KH2).

Raw .czi files were converted to .ims format using the Imaris file converter and subsequently deconvolved using AutoQuant X3 3.1. Processed images were analyzed in Imaris (v 10.2) beginning with nuclear segmentation on the DAPI channel. Spot detection parameters included an XY diameter of 4.5 (determined by measuring average cell diameters in Slice mode), model PSF elongation of 15 µm, background subtraction, quality filter threshold > 1747, and average distance to 3 nearest neighbors between 4.83 and 12.0 µm.

For marker quantification, Pax6 and Nkx2.1 positive cells were identified using identical spot detection parameters with additional colocalization constraints requiring maximum DAPI distances of 14 µm from the center of spot to spot. The entire pipeline was automated through Imaris Arena with parameter consistency across each patterning condition.

Exported quantitative metrics included absolute counts of DAPI+ nuclei, Pax6+/DAPI+ doublepositive cells, and Nkx2.1+/DAPI+ double-positive cells. Statistical analysis of 162 dorsal and 113 ventral images per condition employed Mann-Whitney U test to compare the proportion of cells labeled Pax6 for dorsal versus ventral and Nkx2.1 dorsal versus ventral. Statistical significance was set at p < 0.05.

Quality control measures included blinded analysis (experimenter masked to conditions) (data not shown).

### 13.2 Single-Cell RNA Sequencing and Computational Analysis

Sequencing was performed on an AVITI PE75 Flowcell at the UC Davis Technologies Core, generating 900M reads. Data processing utilized the PIPseeker pipeline (v3.3) with mouse genome GRCm39 (GENCODE vM29 2022.04, Ensembl 106) as reference. FASTQ files were processed with default parameters for alignment, transcript quantification, and cell calling.

Downstream analysis used Seurat (v5.1.0)^136^ with sensitivity 5 matrices. Quality control included:

- Genotype demultiplexing using Souporcell^137^
- Doublet detection with DoubletFinder v2.0.4^138^
- Dataset integration via Harmony^139^

Cells were filtered based on mitochondrial content (>20%), unique gene counts (<5th percentile), and total RNA (>50,000 counts). SCTransform normalized the data while regressing out mitochondrial genes, identifying the top 3,000 variable genes^140,141^. Dimensionality reduction used 40 principal components (selected via elbow plot) for Leiden clustering at resolutions 0.5–2. Cluster visualization employed UMAP^142^, with resolution selection guided by marker gene expression. Cell type annotation referenced the Allen Brain Atlas^143^, UCSC Cell Browser^144^, and Arlotta developmental atlas^3^.

Reference mapping followed Seurat’s integration workflow^136^, combining dorsal forebrain samples, normalizing (log-normalize, scale factor 10,000), identifying variable genes, scaling data, and performing PCA (30 components). Integration used Harmony before transferring annotations via CCA-based anchor identification.

### 13.3 STTC Analysis

We quantified pairwise neuronal synchronization using the STTC with a Δt = 10 ms timescale^21,23,59^. The STTC is defined as:

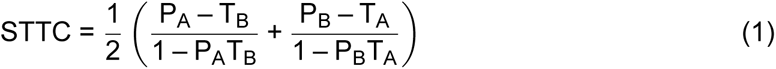

where:

- P_A_ = Proportion of spikes in train A occurring within ±Δt of any spike in train B
- T_A_ = Proportion of the recording duration “tiled” by ±Δt windows around spikes in train A
- P_B_ and T_B_ = Analogous measures for spike train B

This symmetric measure ranges from -1 (perfect anti-correlation) to +1 (perfect synchrony), with 0 indicating independence.

### 13.4 Functional Network Analysis

#### 13.4.1 Network Construction

Functional connectivity matrices were derived from thresholded, binarized STTC values. To establish significance thresholds while preserving population rate dynamics, we:

1. Generated 1000 surrogate datasets by spike identity shuffling
2. Computed STTC distributions from shuffled data
3. Set thresholds at the 90^th^ percentile of null distributions
4. Binarized matrices using these subject-specific thresholds

#### 13.4.2 Global Network Metrics

Using NetworkX (^145^) and custom Numba-accelerated functions, we computed:

- **Clustering coefficient:** Local density of connections using a Numba-accelerated parallel implementation (`compute_clustering_coeff_parallel`)
- **Characteristic path length:** Mean shortest path distance using NetworkX’s `average_shortest_path_length` on the largest connected component

All metrics were normalized by dividing by corresponding values from 100 synthetic random networks (generated via `generate_random_graph`) with identical node and edge counts. Small-worldness was calculated as:

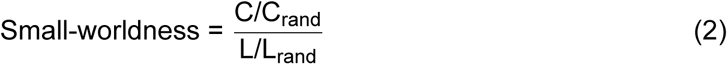

where C and L are clustering coefficient and path length, respectively. Binary functional networks were created using spike time tiling coefficients (STTC) thresholded at the 90th percentile of surrogate values obtained by shuffling neuron identities across 1000 randomized networks while preserving firing rate distributions.

#### 13.4.3 Hub Identification

We computed a composite hubness score integrating four nodal metrics:

- **Degree**: Number of connections (`degrees_und`)
- **Strength**: Sum of connection weights (`strengths_und`, using weighted matrices)
- **Betweenness centrality**: Fraction of shortest paths passing through node (`betweenness_bin`)
- **Closeness centrality**: Inverse average shortest path length (`distance_bin` derived)

Each metric was z-scored across nodes before summation to create the composite score. Analysis computed:

- Firing rate distributions (mean ± SEM across replicates)
- Coefficient of variation (CV) of interspike intervals
- Population synchrony (pairwise spike train correlations)
- E/I balance ratios (excitatory vs inhibitory input currents)
- Weight distribution evolution (Kolmogorov-Smirnov tests)

## References

1. Cadwell, C. R., Bhaduri, A., Mostajo-Radji, M. A., Keefe, M. G., and Nowakowski, T. J. (2019). Development and arealization of the cerebral cortex. Neuron, 103 (6), 980–1004. DOI: 10.1016/j.neuron.2019.07.009.

2. Nowakowski, T. J., Bhaduri, A., Pollen, A. A., Alvarado, B., Mostajo-Radji, M. A., Di Lullo, E., Haeussler, M., Sandoval-Espinosa, C., Liu, S. J., Velmeshev, D., et al. (2017). Spatiotemporal gene expression trajectories reveal developmental hierarchies of the human cortex. Science, 358 (6368), 1318–1323. DOI: 10.1126/science.aap8809.

3. Di Bella, D. J., Habibi, E., Stickels, R. R., Scalia, G., Brown, J., Yadollahpour, P., Yang, S. M., Abbate, C., Biancalani, T., Macosko, E. Z., et al. (2021). Molecular Logic of Cellular Diversification in the Mouse Cerebral Cortex. Nature, 595 (7868), 554–559. DOI: 10.1038/s41586-021-03670-5.

4. Velasco, S., Paulsen, B., and Arlotta, P. (2020). 3D Brain Organoids: Studying Brain Development and Disease Outside the Embryo. Annu. Rev. Neurosci. 43, 375–389. DOI: 10.1146/annurev-neuro-070918-050154.

5. Mostajo-Radji, M. A., Schmitz, M. T., Montoya, S. T., and Pollen, A. A. (2020). Reverse engineering human brain evolution using organoid models. Brain Res. 1729, 146582. DOI: 10.1016/j.brainres.2019.146582.

6. Kyrousi, C. and Cappello, S. (2020). Using brain organoids to study human neurodevelopment, evolution and disease. *WIREs Dev*. Biol. 9 (1), e347. DOI: 10.1002/wdev.347.

7. Velasco, S., Kedaigle, A. J., Simmons, S. K., Nash, A., Rocha, M., Quadrato, G., Paulsen, B., Nguyen, L., Adiconis, X., Regev, A., et al. (2019). Individual Brain Organoids Reproducibly Form Cell Diversity of the Human Cerebral Cortex. Nature, 570 (7762), 523–527. DOI: 10.1038/s41586-019-1289-x.

8. Quadrato, G., Nguyen, T., Macosko, E. Z., Sherwood, J. L., Min Yang, S., Berger, D. R., Maria, N., Scholvin, J., Goldman, M., Kinney, J. P., et al. (2017). Cell Diversity and Network Dynamics in Photosensitive Human Brain Organoids. Nature, 545 (7652), 48–53. DOI: 10.1038/nature22047.

9. Pollen, A. A., Bhaduri, A., Andrews, M. G., Nowakowski, T. J., Meyerson, O. S., MostajoRadji, M. A., Di Lullo, E., Alvarado, B., Bedolli, M., Dougherty, M. L., et al. (2019). Establishing Cerebral Organoids as Models of Human-Specific Brain Evolution. Cell, 176 (4), 743–756.e17. DOI: 10.1016/j.cell.2019.01.017.

10. Nolbrant, S., Wallace, J. L., Ding, J., Zhu, T., Sevetson, J. L., Kajtez, J., Baldacci, I. A., Corrigan, E. K., Hoglin, K., McMullen, R., et al. (2024). Interspecies Organoids Reveal Human-Specific Molecular Features of Dopaminergic Neuron Development and Vulnerability. Preprint at BioRxiv, DOI: 10.1101/2024.11.14.623592.

11. Sharf, T., van der Molen, T., Glasauer, S. M. K., Guzman, E., Buccino, A. P., Luna, G., Cheng, Z., Audouard, M., Ranasinghe, K. G., Kudo, K., et al. (2022). Functional Neuronal Circuitry and Oscillatory Dynamics in Human Brain Organoids. Nat. Commun. 13 (1), 4403. DOI: 10.1038/s41467-022-32115-4.

12. Van der Molen, T., Spaeth, A., Chini, M., Hernandez, S., Kaurala, G. A., Schweiger, H. E., Duncan, C., McKenna, S., Geng, J., Lim, M., et al. (2025). Protosequences in brain organoids model intrinsic brain states. Preprint at BioRxiv, DOI: 10.1101/2023.12.29.573646.

13. Trujillo, C. A., Gao, R., Negraes, P. D., Gu, J., Buchanan, J., Preissl, S., Wang, A., Wu, W., Haddad, G. G., Chaim, I. A., et al. (2019). Complex Oscillatory Waves Emerging from Cortical Organoids Model Early Human Brain Network Development. Cell Stem Cell, 25 (4), 558–569.e7. DOI: 10.1016/j.stem.2019.08.002.

14. Zafeiriou, M.-P., Bao, G., Hudson, J., Halder, R., Blenkle, A., Schreiber, M.-K., Fischer, A., Schild, D., and Zimmermann, W.-H. (2020). Developmental GABA polarity switch and neuronal plasticity in Bioengineered Neuronal Organoids. Nat. Commun. 11 (1), 3791. DOI: 10.1038/s41467-020-17521-w.

15. Samarasinghe, R. A., Miranda, O. A., Buth, J. E., Mitchell, S., Ferando, I., Watanabe, M., Allison, T. F., Kurdian, A., Fotion, N. N., Gandal, M. J., et al. (2021). Identification of neural oscillations and epileptiform changes in human brain organoids. Nat. Neurosci. 24 (10), 1488–1500. DOI: 10.1038/s41593-021-00906-5.

16. Kang, R., Park, S., Shin, S., Bak, G., and Park, J.-C. (2024). Electrophysiological Insights with Brain Organoid Models: A Brief Review. BMB Rep. 57 (7), 311–317. DOI: 10.5483/BMBRep.2024-0077.

17. Gu, L., Cai, H., Chen, L., Gu, M., Tchieu, J., and Guo, F. (2024). Functional Neural Networks in Human Brain Organoids. BME Front. 5, 0065. DOI: 10.34133/bmef.0065.

18. Humpel, C. (2015). Organotypic brain slice cultures: A review. Neuroscience, 305, 86–98. DOI: 10.1016/j.neuroscience.2015.07.086.

19. Wu, M. W., Kourdougli, N., and Portera-Cailliau, C. (2024). Network state transitions during cortical development. Nat. Rev. Neurosci. 25 (8), 535–552. DOI: 10.1038/s41583-024-00824-y.

20. Golshani, P., Gonçalves, J. T., Khoshkhoo, S., Mostany, R., Smirnakis, S., and Portera-Cailliau, C. (2009). Internally Mediated Developmental Desynchronization of Neocortical Network Activity. J. Neurosci. 29 (35), 10890–10899. DOI: 10.1523/JNEUROSCI.2012-09.2009.

21. Chini, M., Pfeffer, T., and Hanganu-Opatz, I. (2022). An Increase of Inhibition Drives the Developmental Decorrelation of Neural Activity. eLife, 11, e78811. DOI: 10.7554/eLife.78811.

22. Contractor, A., Ethell, I. M., and Portera-Cailliau, C. (2021). Cortical interneurons in autism. Nat. Neurosci. 24 (12), 1648–1659. DOI: 10.1038/s41593-021-00967-6.

23. Chini, M., Hnida, M., Kostka, J. K., Chen, Y.-N., and Hanganu-Opatz, I. L. (2024). Pre-configured Architecture of the Developing Mouse Brain. Cell Rep. 43 (6), 114267. DOI: 10.1016/j.celrep.2024.114267.

24. Hilgetag, C. C. and Kaiser, M. (2004). Clustered Organization of Cortical Connectivity. Neuroinformatics, 2 (3), 353–360. DOI: 10.1385/NI:2:3:353.

25. Sporns, O. and Zwi, J. D. (2004). The small world of the cerebral cortex. Neuroinformatics, 2 (2), 145–162. DOI: 10.1385/NI:2:2:145.

26. Watanabe, K., Kamiya, D., Nishiyama, A., Katayama, T., Nozaki, S., Kawasaki, H., Watanabe, Y., Mizuseki, K., and Sasai, Y. (2005). Directed Differentiation of Telencephalic Precursors from Embryonic Stem Cells. Nat. Neurosci. 8 (3), 288–296. DOI: 10.1038/nn1402.

27. Eiraku, M., Watanabe, K., Matsuo-Takasaki, M., Kawada, M., Yonemura, S., Matsumura, M., Wataya, T., Nishiyama, A., Muguruma, K., and Sasai, Y. (2008). Self-Organized Formation of Polarized Cortical Tissues from ESCs and Its Active Manipulation by Extrinsic Signals. Cell Stem Cell, 3 (5), 519–532. DOI: 10.1016/j.stem.2008.09.002.

28. Kadoshima, T., Sakaguchi, H., Nakano, T., Soen, M., Ando, S., Eiraku, M., and Sasai, Y. (2013). Self-Organization of Axial Polarity, Inside-Out Layer Pattern, and Species-Specific Progenitor Dynamics in Human ES Cell-Derived Neocortex. Proc. Natl. Acad. Sci. USA, 110 (50), 20284–20289. DOI: 10.1073/pnas.1315710110.

29. Lancaster, M. A. and Knoblich, J. A. (2014). Generation of Cerebral Organoids from Human Pluripotent Stem Cells. Nat. Protoc. 9 (10), 2329–2340. DOI: 10.1038/nprot.2014.158.

30. Paşca, A. M., Sloan, S. A., Clarke, L. E., Tian, Y., Makinson, C. D., Huber, N., Kim, C. H., Park, J.-Y., O’Rourke, N. A., Nguyen, K. D., et al. (2015). Functional Cortical Neurons and Astrocytes from Human Pluripotent Stem Cells in 3D Culture. Nat. Methods, 12 (7), 671– 678. DOI: 10.1038/nmeth.3415.

31. Voitiuk, K., Seiler, S. T., Pessoa de Melo, M., Geng, J., van der Molen, T., Hernandez, S., Schweiger, H. E., Sevetson, J. L., Parks, D. F., Robbins, A., et al. (2024). A Feedback-Driven Brain Organoid Platform Enables Automated Maintenance and High-Resolution Neural Activity Monitoring. Preprint at BioRxiv, DOI: 10.1101/2024.03.15.585237.

32. Elliott, M. A., Schweiger, H. E., Robbins, A., Vera-Choqqueccota, S., Ehrlich, D., Hernandez, S., Voitiuk, K., Geng, J., Sevetson, J. L., Rosen, Y. M., et al. (2023). Internet-Connected Cortical Organoids for Project-Based Stem Cell and Neuroscience Education. eNeuro, 10 (12), ENEURO.0308–23.2023. DOI: 10.1101/2023.07.13.546418.

33. Park, Y., Hernandez, S., Hernandez, C. O., Schweiger, H. E., Li, H., Voitiuk, K., Dechiraju, H., Hawthorne, N., Muzzy, E. M., Selberg, J. A., et al. (2024). Modulation of neuronal activity in cortical organoids with bioelectronic delivery of ions and neurotransmitters. Cell Reports Methods, 4 (1), 100686. DOI: 10.1016/j.crmeth.2023.100686.

34. Ciarpella, F., Zamfir, R. G., Campanelli, A., Ren, E., Pedrotti, G., Bottani, E., Borioli, A., Caron, D., Chio, M. D., Dolci, S., et al. (2021). Murine cerebral organoids develop network of functional neurons and hippocampal brain region identity. iScience, 24 (12), 103438. DOI: 10.1016/j.isci.2021.103438.

35. Mostajo-Radji, M. A., Leon, W. R. M., Breevoort, A., Gonzalez-Ferrer, J., Schweiger, H. E., Lehrer, J., Zhou, L., Schmitz, M. T., Perez, Y., Mukhtar, T., et al. (2025). Fate Plasticity of Interneuron Specification. iScience, 28 (4), 112295. DOI: 10.1016/j.isci.2025.112295.

36. Sánchez, D. J. L.-D., Lindhout, F. W., Anderson, A. J., Pellegrini, L., and Lancaster, M. A. (2024). Mouse Brain Organoids Model In Vivo Neurodevelopment and Function and Capture Differences to Human. Preprint at BioRxiv, DOI: 10.1101/2024.12.21.629881.

37. Lindhout, F. W., Szafranska, H. M., Imaz-Rosshandler, I., Guglielmi, L., Moarefian, M., Voitiuk, K., Zernicka-Glover, N. K., Boulanger, J., Schulze, U., Sánchez, D. J. L.-D., et al. (2025). Calcium Dynamics Tune Developmental Tempo to Generate Evolutionarily Divergent Axon Tract Lengths. Preprint at BioRxiv, DOI: 10.1101/2024.12.28.630576.

38. Medina-Cano, D., Corrigan, E. K., Glenn, R. A., Islam, M. T., Lin, Y., Kim, J., Cho, H., and Vierbuchen, T. (2022). Rapid and robust directed differentiation of mouse epiblast stem cells into definitive endoderm and forebrain organoids. Development, 149 (20), dev200561. DOI: 10.1242/dev.200561.

39. Medina-Cano, D., Islam, M. T., Petrova, V., Dixit, S., Balic, Z., Yang, M. G., Stadtfeld, M., Wong, E. S., and Vierbuchen, T. (2024). A Mouse Organoid Platform for Modeling Cerebral Cortex Development and Cis-Regulatory Evolution in Vitro. Preprint at BioRxiv, DOI: 10.1101/2024.09.30.615887.

40. Li, Y., Mao, X., Zhou, X., Su, Y., Zhou, X., Shi, K., and Zhao, S. (2020). An Optimized Method for Neuronal Differentiation of Embryonic Stem Cells in Vitro. J. Neurosci. Methods, 330, 108486. DOI: 10.1016/j.jneumeth.2019.108486.

41. Hevner, R. F., Daza, R. A., Rubenstein, J. L., Stunnenberg, H., Olavarria, J. F., and Englund, C. (2003). Beyond Laminar Fate: Toward a Molecular Classification of Cortical Projection/Pyramidal Neurons. Dev. Neurosci. 25 (2-4), 139–151. DOI: 10.1159/000072263.

42. Molyneaux, B. J., Arlotta, P., Menezes, J. R. L., and Macklis, J. D. (2007). Neuronal Subtype Specification in the Cerebral Cortex. Nat. Rev. Neurosci. 8 (6), 427–437. DOI: 10.1038/nrn2151.

43. Fame, R. M., MacDonald, J. L., and Macklis, J. D. (2011). Development, Specification, and Diversity of Callosal Projection Neurons. Trends Neurosci. 34 (1), 41–50. DOI: 10.1016/j.tins.2010.10.002.

44. Wang, D. D. and Kriegstein, A. R. (2009). Defining the Role of GABA in Cortical Development. J. Physiol. 587 (9), 1873–1879. DOI: 10.1113/jphysiol.2008.167635.

45. Del Rio, J. A., Soriano, E., and Ferrer, I. (1992). Development of GABA-Immunoreactivity in the Neocortex of the Mouse. J. Comp. Neurol. 326 (4), 501–526. DOI: 10.1002/cne.903260403.

46. Hagino-Yamagishi, K., Saijoh, Y., Ikeda, M., Ichikawa, M., Minamikawa-Tachino, R., and Hamada, H. (1997). Predominant Expression of Brn-2 in the Postmitotic Neurons of the Developing Mouse Neocortex. Brain Res. 752 (1-2), 261–268. DOI: 10.1016/s0006-8993(96)01472-2.

47. Dominguez, M. H., Ayoub, A. E., and Rakic, P. (2013). POU-III Transcription Factors (Brn1, Brn2, and Oct6) Influence Neurogenesis, Molecular Identity, and Migratory Destination of Upper-Layer Cells of the Cerebral Cortex. Cereb. Cortex, 23 (11), 2632–2643. DOI: 10.1093/cercor/bhs252.

48. Arlotta, P., Molyneaux, B. J., Chen, J., Inoue, J., Kominami, R., and Macklis, J. D. (2005). Neuronal Subtype-Specific Genes That Control Corticospinal Motor Neuron Development In Vivo. Neuron, 45 (2), 207–221. DOI: 10.1016/j.neuron.2004.12.036.

49. Ahtiainen, A., Genocchi, B., Subramaniyam, N. P., Tanskanen, J. M. A., Rantamäki, T., and Hyttinen, J. A. K. (2024). Astrocytes Facilitate Gabazine-Evoked Electrophysiological Hyperactivity and Distinct Biochemical Responses in Mature Neuronal Cultures. J. Neurochem. 168 (9), 3076–3094. DOI: 10.1111/jnc.16182.

50. Allen, N. J., Bennett, M. L., Foo, L. C., Wang, G. X., Chakraborty, C., Smith, S. J., and Barres, B. A. (2012). Astrocyte Glypicans 4 and 6 Promote Formation of Excitatory Synapses via GluA1 AMPA Receptors. Nature, 486 (7403), 410–414. DOI: 10.1038/nature11059.

51. Ullian, E. M., Sapperstein, S. K., Christopherson, K. S., and Barres, B. A. (2001). Control of Synapse Number by Glia. Science, 291 (5504), 657–661. DOI: 10.1126/science.291.5504.657.

52. Walsh, R. M., Crabtree, G. W., Kalpana, K., Jubierre, L., Koo, S. Y., Ciceri, G., Gogos, J. A., Kruglikov, I., and Studer, L. (2024). Cortical Assembloids Support the Development of Fast-Spiking Human PVALB+ Cortical Interneurons and Uncover Schizophrenia-Associated Defects. Preprint at BioRxiv, DOI: 10.1101/2024.11.26.624368.

53. Köntgen, F., Süss, G., Stewart, C., Steinmetz, M., and Bluethmann, H. (1993). Targeted Disruption of the MHC Class II Aa Gene in C57BL/6 Mice. Int. Immunol. 5 (8), 957–964. DOI: 10.1093/intimm/5.8.957.

54. Hooper, M., Hardy, K., Handyside, A., Hunter, S., and Monk, M. (1987). HPRT-Deficient (Lesch-Nyhan) Mouse Embryos Derived from Germline Colonization by Cultured Cells. Nature, 326 (6110), 292–295. DOI: 10.1038/326292a0.

55. Beard, C., Hochedlinger, K., Plath, K., Wutz, A., and Jaenisch, R. (2006). Efficient Method to Generate Single-Copy Transgenic Mice by Site-Specific Integration in Embryonic Stem Cells. Genesis, 44 (1), 23–28. DOI: 10.1002/gene.20180.

56. Shin, D., Kim, C. N., Ross, J., Hennick, K. M., Wu, S.-R., Paranjape, N., Leonard, R., Wang, J. C., Keefe, M. G., Pavlovic, B. J., et al. (2024). Thalamocortical Organoids Enable in Vitro Modeling of 22q11.2 Microdeletion Associated With Neuropsychiatric Disorders. Cell Stem Cell, 31 (3), 421–432.e8. DOI: 10.1016/j.stem.2024.01.010.

57. Amin, N. D., Kelley, K. W., Kaganovsky, K., Onesto, M., Hao, J., Miura, Y., McQueen, J. P., Reis, N., Narazaki, G., Li, T., et al. (2024). Generating Human Neural Diversity With a Multiplexed Morphogen Screen in Organoids. Cell Stem Cell, 31 (12), 1831–1846.e9. DOI: 10.1016/j.stem.2024.10.016.

58. Andrews, J. P., Geng, J., Voitiuk, K., Elliott, M. A. T., Shin, D., Robbins, A., Spaeth, A., Wang, A., Li, L., Solis, D., et al. (2024). Multimodal Evaluation of Network Activity and Optogenetic Interventions in Human Hippocampal Slices. Nat. Neurosci. 27 (12), 2487– 2499. DOI: 10.1038/s41593-024-01782-5.

59. Cutts, C. S. and Eglen, S. J. (2014). Detecting Pairwise Correlations in Spike Trains: An Objective Comparison of Methods and Application to the Study of Retinal Waves. J. Neurosci. 34 (43), 14288–14303. DOI: 10.1523/JNEUROSCI.2767-14.2014.

60. Rochefort, N. L., Garaschuk, O., Milos, R.-I., Narushima, M., Marandi, N., Pichler, B., Kovalchuk, Y., and Konnerth, A. (2009). Sparsification of Neuronal Activity in the Visual Cortex at Eye-Opening. Proc. Natl. Acad. Sci. USA, 106 (35), 15049–15054. DOI: 10.1073/pnas.0907660106.

61. Buzsáki, G. and Mizuseki, K. (2014). The Log-Dynamic Brain: How Skewed Distributions Affect Network Operations. Nat. Rev. Neurosci. 15 (4), 264–278. DOI: 10.1038/nrn3687.

62. Crocco, E., Iannello, L., Tonelli, F., Lagani, G., Pandolfini, L., Amato, G., Di Garbo, A., and Cremisi, F. (2025). A Proper Excitatory/Inhibitory Ratio Is Required to Develop Synchronized Network Activity in Mouse Cortical Cultures. Preprint at BioRxiv, DOI: 10.1101/2025.02.28.640720.

63. Myme, C. I. O., Sugino, K., Turrigiano, G. G., and Nelson, S. B. (2003). The NMDA-to-AMPA Ratio at Synapses Onto Layer 2/3 Pyramidal Neurons Is Conserved Across Prefrontal and Visual Cortices. J. Neurophysiol. 90 (2), 771–779. DOI: 10.1152/jn.00070.2003.

64. Barbero-Castillo, A., Mateos-Aparicio, P., Porta, L. D., Camassa, A., Perez-Mendez, L., and Sanchez-Vives, M. V. (2021). Impact of GABAA and GABAB Inhibition on Cortical Dynamics and Perturbational Complexity During Synchronous and Desynchronized States. J. Neurosci. 41 (23), 5029–5044. DOI: 10.1523/JNEUROSCI.1837-20.2021.

65. Birey, F., Andersen, J., Makinson, C. D., Islam, S., Wei, W., Huber, N., Fan, H. C., Metzler, K. R. C., Panagiotakos, G., Thom, N., et al. (2017). Assembly of Functionally Integrated Human Forebrain Spheroids. Nature, 545 (7652), 54–59. DOI: 10.1038/nature22330.

66. Nasu, M., Takata, N., Danjo, T., Sakaguchi, H., Kadoshima, T., Futaki, S., Sekiguchi, K., Eiraku, M., and Sasai, Y. (2012). Robust Formation and Maintenance of Continuous Stratified Cortical Neuroepithelium by Laminin-Containing Matrix in Mouse ES Cell Culture. PLoS ONE, 7 (12), e53024. DOI: 10.1371/journal.pone.0053024.

67. Xiang, Y., Yoshiaki, T., Patterson, B., Cakir, B., Kim, K.-Y., Cho, Y. S., and Park, I.-H. (2018). Generation and Fusion of Human Cortical and Medial Ganglionic Eminence Brain Organoids. Curr. Protoc. Stem Cell Biol. 47 (1), e61. DOI: 10.1002/cpsc.61.

68. Pavon, N., Diep, K., Yang, F., Sebastian, R., Martinez-Martin, B., Ranjan, R., Sun, Y., and Pak, C. (2024). Patterning Ganglionic Eminences in Developing Human Brain Organoids Using a Morphogen-Gradient-Inducing Device. *Cell Rep*. Methods, 4 (1), 100689. DOI: 10.1016/j.crmeth.2023.100689.

69. Stanton, B. Z. and Peng, L. F. (2010). Small-molecule modulators of the Sonic Hedgehog signaling pathway. Mol. BioSyst. 6 (1), 44–54. DOI: 10.1039/B910196A.

70. Flandin, P., Kimura, S., and Rubenstein, J. L. (2010). The progenitor zone of the ventral medial ganglionic eminence requires Nkx2-1 to generate most of the globus pallidus but few neocortical interneurons. J. Neurosci. 30 (8), 2812–2823. DOI: 10.1523/JNEUROSCI.4228-09.2010.

71. Warren, N., Caric, D., Pratt, T., Clausen, J. A., Asavaritikrai, P., Mason, J. O., Hill, R. E., and Price, D. J. (1999). The transcription factor, Pax6, is required for cell proliferation and differentiation in the developing cerebral cortex. Cereb. Cortex, 9 (6), 627–635. DOI: 10.1002/dneu.20895.

72. Holter, M. C., Hewitt, L. T., Nishimura, K. J., Knowles, S. J., Bjorklund, G. R., Shah, S., Fry, N. R., Rees, K. P., Gupta, T. A., Daniels, C. W., et al. (2021). Hyperactive MEK1 Signaling in Cortical GABAergic Neurons Promotes Embryonic Parvalbumin Neuron Loss and Defects in Behavioral Inhibition. Cereb. Cortex, 31 (6), 3064–3081. DOI: 10.1093/cercor/bhaa413.

73. Knowles, S. J., Holter, M. C., Li, G., Bjorklund, G. R., Rees, K. P., Martinez-Fuentes, J. S., Nishimura, K. J., Afshari, A. E., Fry, N., Stafford, A. M., et al. (2023). Multifunctional Requirements for ERK1/2 Signaling in the Development of Ganglionic Eminence Derived Glia and Cortical Inhibitory Neurons. eLife, 12 DOI: 10.7554/eLife.88313.1.

74. Wen, T. H., Binder, D. K., Ethell, I. M., and Razak, K. A. (2018). The Perineuronal ‘Safety’ Net? Perineuronal Net Abnormalities in Neurological Disorders. Front. Mol. Neurosci. 11, 270. DOI: 10.3389/fnmol.2018.00270.

75. Bassett, D. S. and Bullmore, E. T. (2017). Small-World Brain Networks Revisited. Neuroscientist, 23 (5), 499–516. DOI: 10.1177/1073858416667720.

76. Bassett, D. S. and Bullmore, E. (2006). Small-world brain networks. Neuroscientist, 12 (6), 512–523.

77. Gerhard, F., Pipa, G., Lima, B., Neuenschwander, S., and Gerstner, W. (2011). Extraction of Network Topology From Multi-Electrode Recordings: Is There a Small-World Effect? Front. Comput. Neurosci. 5, 4. DOI: 10.3389/fncom.2011.00004.

78. Akarca, D., Dunn, A. W., Hornauer, P. J., Ronchi, S., Fiscella, M., Wang, C., Terrigno, M., Jagasia, R., Vertes, P. E., Mierau, S. B., et al. (2022). Homophilic Wiring Principles Underpin Neuronal Network Topology In Vitro. Preprint at BioRxiv, DOI: 10.1101/2022.03.09.483605.

79. Antonello, P. C., Varley, T. F., Beggs, J., Porcionatto, M., Sporns, O., and Faber, J. (2022). Self-Organization of In Vitro Neuronal Assemblies Drives to Complex Network Topology. eLife, 11, e74921. DOI: 10.7554/eLife.74921.

80. Achard, S., Salvador, R., Whitcher, B., Suckling, J., and Bullmore, E. (2006). A resilient, low-frequency, small-world human brain functional network with highly connected association cortical hubs. J. Neurosci. 26 (1), 63–72. DOI: 10.1523/JNEUROSCI.3874-05.2006.

81. Okun, M., Steinmetz, N. A., Cossell, L., Iacaruso, M. F., Ko, H., Barthó, P., Moore, T., Hofer, S. B., Mrsic-Flogel, T. D., Carandini, M., et al. (2015). Diverse Coupling of Neurons to Populations in Sensory Cortex. Nature, 521 (7553), 511–515. DOI: 10.1038/nature14273.

82. Okun, M., Yger, P., Marguet, S. L., Gerard-Mercier, F., Benucci, A., Katzner, S., Busse, L., Carandini, M., and Harris, K. D. (2012). Population Rate Dynamics and Multineuron Firing Patterns in Sensory Cortex. J. Neurosci. 32 (48), 17108–17119. DOI: 10.1523/JNEUROSCI.1831-12.2012.

83. Lahav, N., Ksherim, B., Ben-Simon, E., Maron-Katz, A., Cohen, R., and Havlin, S. (2016). K-shell decomposition reveals hierarchical cortical organization of the human brain. New J. Phys. 18 (8), 083013. DOI: 10.1088/1367-2630/18/8/083013.

84. Bassett, D. S., Wymbs, N. F., Rombach, M. P., Porter, M. A., Mucha, P. J., and Grafton, S. T. (2013). Task-Based Core-Periphery Organization of Human Brain Dynamics. PLoS Comput. Biol. 9 (9), e1003171. DOI: 10.1371/journal.pcbi.1003171.

85. Gu, S., Xia, C. H., Ciric, R., Moore, T. M., Gur, R. C., Gur, R. E., Satterthwaite, T. D., and Bassett, D. S. (2020). Unifying the Notions of Modularity and Core-Periphery Structure in Functional Brain Networks During Youth. Cereb. Cortex, 30 (3), 1087–1102. DOI: 10.1093/cercor/bhz150.

86. Sit, T. P. H., Feord, R. C., Dunn, A. W. E., Chabros, J., Oluigbo, D., Smith, H. H., Burn, L., Chang, E., Boschi, A., Yuan, Y., et al. (2024). MEA-NAP: A Flexible Network Analysis Pipeline for Neuronal 2D and 3D Organoid Multielectrode Recordings. *Cell Rep*. Methods, 4 (11), 100901. DOI: 10.1016/j.crmeth.2024.100901.

87. Bollmann, Y., Modol, L., Tressard, T., Vorobyev, A., Dard, R., Brustlein, S., Sims, R., Bendifallah, I., Leprince, E., de Sars, V., et al. (2023). Prominent In Vivo Influence of Single Interneurons in the Developing Barrel Cortex. Nat. Neurosci. 26 (9), 1555–1565. DOI: 10.1038/s41593-023-01405-5.

88. Vega-Zuniga, T., Sumser, A., Symonova, O., Koppensteiner, P., Schmidt, F. H., and Joesch, M. (2025). A Thalamic Hub-and-Spoke Network Enables Visual Perception During Action by Coordinating Visuomotor Dynamics. Nat. Neurosci. 28 (3), 627–639. DOI: 10.1038/s41593-025-01874-w.

89. Jin, S.-H., Jeong, W., Seol, J., Kwon, J., and Chung, C. K. (2013). Functional Cortical Hubs in the Eyes-Closed Resting Human Brain from an Electrophysiological Perspective Using Magnetoencephalography. PLoS ONE, 8 (7), e68192. DOI: 10.1371/journal.pone.0068192.

90. Hanalioglu, S., Bahadir, S., Isikay, I., Celtikci, P., Celtikci, E., Yeh, F.-C., Oguz, K. K., and Khaniyev, T. (2021). Group-Level Ranking-Based Hubness Analysis of Human Brain Connectome Reveals Significant Interhemispheric Asymmetry and Intraparcel Heterogeneities. Front. Neurosci. 15, 782995. DOI: 10.3389/fnins.2021.782995.

91. Mòdol, L., Sousa, V. H., Malvache, A., Tressard, T., Baude, A., and Cossart, R. (2017). Spatial Embryonic Origin Delineates GABAergic Hub Neurons Driving Network Dynamics in the Developing Entorhinal Cortex. Cereb. Cortex, 27 (9), 4649–4661. DOI: 10.1093/cercor/bhx198.

92. Picardo, M., Guigue, P., Bonifazi, P., Batista-Brito, R., Allene, C., Ribas, A., Fishell, G., Baude, A., and Cossart, R. (2011). Pioneer GABA Cells Comprise a Subpopulation of Hub Neurons in the Developing Hippocampus. Neuron, 71 (4), 695–709. DOI: 10.1016/j.neuron.2011.06.018.

93. Grosmark, A. D. and Buzsáki, G. (2016). Diversity in neural firing dynamics supports both rigid and learned hippocampal sequences. Science, 351 (6280), 1440–1443. DOI: 10.1126/science.aad1935.

94. Vaz, A. P., Wittig Jr, J. H., Inati, S. K., and Zaghloul, K. A. (2023). Backbone spiking sequence as a basis for preplay, replay, and default states in human cortex. Nat. Commun. 14 (1), 4723. DOI: 10.1038/s41467-023-40440-5.

95. Schuurman, T. and Bruner, E. (2024). Modularity and community detection in human brain morphology. Anat. Rec. 307 (2), 345–355. DOI: 10.1002/ar.25308.

96. Hensch, T. K. (2005). Critical Period Plasticity in Local Cortical Circuits. Nat. Rev. Neurosci. 6 (11), 877–888. DOI: 10.1038/nrn1787.

97. Freund, T. F. and Katona, I. (2007). Perisomatic Inhibition. Neuron, 56 (1), 33–42. DOI: 10.1016/j.neuron.2007.09.012.

98. Ye, Z., Mostajo-Radji, M. A., Brown, J. R., Rouaux, C., Tomassy, G. S., Hensch, T. K., and Arlotta, P. (2015). Instructing Perisomatic Inhibition by Direct Lineage Reprogramming of Neocortical Projection Neurons. Neuron, 88 (3), 475–483. DOI: 10.1016/j.neuron.2015.10.006.

99. Ding, L., Balsamo, G., Chen, H., Blanco-Hernandez, E., Zouridis, I. S., Naumann, R., Preston-Ferrer, P., and Burgalossi, A. (2022). Juxtacellular opto-tagging of hippocampal CA1 neurons in freely moving mice. eLife, 11, e71720. DOI: 10.7554/eLife.71720.

100. Beau, M., Herzfeld, D. J., Naveros, F., Hemelt, M. E., D’Agostino, F., Oostland, M., Sánchez-López, A., Chung, Y. Y., Maibach, M., Kyranakis, S., et al. (2025). A deeplearning strategy to identify cell types across species from high-density extracellular recordings. Cell, 188 (8), 2218–2234.e22. DOI: 10.1016/j.cell.2025.01.041.

101. Zou, D., Chen, L., Deng, D., Jiang, D., Dong, F., McSweeney, C., Zhou, Y., Liu, L., Chen, G., Wu, Y., et al. (2016). DREADD in parvalbumin interneurons of the dentate gyrus modulates anxiety, social interaction and memory extinction. Curr. Mol. Med. 16 (1), 91–102. DOI: 10.2174/1566524016666151222150024.

102. De Vries, S. E., Siegle, J. H., and Koch, C. (2023). Sharing neurophysiology data from the Allen Brain Observatory. eLife, 12, e85550. DOI: 10.7554/eLife.85550.

103. Gonzalez-Ferrer, J., Lehrer, J., Schweiger, H. E., Geng, J., Hernandez, S., Reyes, F., Sevetson, J. L., Salama, S., Teodorescu, M., Haussler, D., et al. (2025). HIPPIE: A Multimodal Deep Learning Model for Electrophysiological Classification of Neurons. Preprint at BioRxiv, DOI: 10.1101/2025.03.14.642461.

104. Dorrn, A. L., Yuan, K., Barker, A. J., Schreiner, C. E., and Froemke, R. C. (2010). Developmental sensory experience balances cortical excitation and inhibition. Nature, 465 (7300), 932–936. DOI: 10.1038/nature09119.

105. Anton-Bolanos, N., Sempere-Ferrandez, A., Guillamon-Vivancos, T., Martini, F. J., Perez-Saiz, L., Gezelius, H., Filipchuk, A., Valdeolmillos, M., and Lopez-Bendito, G. (2019). Prenatal activity from thalamic neurons governs the emergence of functional cortical maps in mice. Science, 364 (6444), 987–990. DOI: 10.1126/science.aav7617.

106. Martini, F. J., Guillamon-Vivancos, T., Moreno-Juan, V., Valdeolmillos, M., and Lopez-Bendito, G. (2021). Spontaneous activity in developing thalamic and cortical sensory networks. Neuron, 109 (16), 2519–2534. DOI: 10.1016/j.neuron.2021.06.026.

107. Anibal-Martinez, M., Puche-Aroca, L., Perez-Montoyo, E., Pumo, G., Madrigal, M. P., Rodriguez-Malmierca, L. M., Martini, F. J., Rijli, F. M., and Lopez-Bendito, G. (2025). A prenatal window for enhancing spatial resolution of cortical barrel maps. Nat. Commun. 16 (1), 1955. DOI: 10.1038/s41467-025-57052-w.

108. Anton-Bolanos, N., Espinosa, A., and Lopez-Bendito, G. (2018). Developmental interactions between thalamus and cortex: a true love reciprocal story. Curr. Opin. Neurobiol. 52, 33–41. DOI: 10.1016/j.conb.2018.04.018.

109. Adli, M., Przybyla, L., Burdett, T., Burridge, P. W., Cacheiro, P., Chang, H. Y., Engreitz, J. M., Gilbert, L. A., Greenleaf, W. J., Hsu, L., et al. (2025). MorPhiC Consortium: towards functional characterization of all human genes. Nature, 638 (8050), 351–359. DOI: 10.1038/s41586-024-08243-w.

110. Gonzalez-Ferrer, J. and Mostajo-Radji, M. A. (2025). Towards Automated and Explainable High-Throughput Perturbation Analysis in Single Cells. Patterns, 6 (4), 101228. DOI: 10.1016/j.patter.2025.101228.

111. Zhang, H., McCarroll, A., Peyton, L., de León-Guerrerro, S. D., Zhang, S., Gowda, P., Sirkin, D., ElAchwah, M., Duhe, A., Wood, W. G., et al. (2024). Scaled and efficient derivation of loss-of-function alleles in risk genes for neurodevelopmental and psychiatric disorders in human iPSCs. Stem Cell Rep. 19 (10), 1489–1504. DOI: 10.1016/j.stemcr.2024.08.003.

112. Amos-Landgraf, J., Franklin, C., Godfrey, V., Grieder, F., Grimsrud, K., Korf, I., Lutz, C., Magnuson, T., Mirochnitchenko, O., Patel, S., et al. (2022). The Mutant Mouse Resource and Research Center (MMRRC): the NIH-supported national public repository and distribution archive of mutant mouse models in the USA. Mamm. Genome, 33, 203–212. DOI: 10.1007/s00335-021-09894-0.

113. Hansen, G. M., Markesich, D. C., Burnett, M. B., Zhu, Q., Dionne, K. M., Richter, L. J., Finnell, R. H., Sands, A. T., Zambrowicz, B. P., and Abuin, A. (2008). Large-scale gene trapping in C57BL/6N mouse embryonic stem cells. Genome Res. 18 (10), 1670–1679. DOI: 10.1101/gr.078352.108.

114. Wilkinson, P., Sengerova, J., Matteoni, R., Chen, C.-K., Soulat, G., Ureta-Vidal, A., Fessele, S., Hagn, M., Massimi, M., Pickford, K., et al. (2010). EMMA—mouse mutant resources for the international scientific community. Nucleic Acids Res. 38 (1), D570–D576. DOI: 10.1093/nar/gkp799.

115. Wang, D. C., Santos-Valencia, F., Song, J. H., Franks, K. M., and Luo, L. (2024). Embryonically active piriform cortex neurons promote intracortical recurrent connectivity during development. Neuron, 112 (17), 2938–2954. DOI: 10.1016/j.neuron.2024.06.007.

116. Leighton, A. H., Busch, M. V. F., Coppens, J. E., Heimel, J. A., and Lohmann, C. (2022). Lightweight, wireless LED implant for chronic manipulation in vivo of spontaneous activity in neonatal mice. J. Neurosci. Methods, 373, 109548. DOI: 10.1016/j.jneumeth.2022.109548.

117. Nahar, L., Delacroix, B. M., and Nam, H. W. (2021). The role of parvalbumin interneurons in neurotransmitter balance and neurological disease. Front. Psychiatry, 12, 679960. DOI: 10.3389/fpsyt.2021.679960.

118. Tomassy, G. S., Morello, N., Calcagno, E., and Giustetto, M. (2014). Developmental abnormalities of cortical interneurons precede symptoms onset in a mouse model of Rett syndrome. J. Neurochem. 131 (1), 115–127. DOI: 10.1111/jnc.12803.

119. Ito-Ishida, A., Ure, K., Chen, H., Swann, J. W., and Zoghbi, H. Y. (2015). Loss of MeCP2 in parvalbumin-and somatostatin-expressing neurons in mice leads to distinct Rett syndrome-like phenotypes. Neuron, 88 (4), 651–658. DOI: 10.1016/j.neuron.2015.10.029.

120. Juavinett, A. L., Bekheet, G., and Churchland, A. K. (2019). Chronically implanted Neuropixels probes enable high-yield recordings in freely moving mice. eLife, 8, e47188. DOI: 10.7554/eLife.47188.

121. Hales, C. M., Rolston, J. D., and Potter, S. M. (2010). How to culture, record and stimulate neuronal networks on micro-electrode arrays (MEAs). J. Vis. Exp.: JoVE, (39), 2056. DOI: 10.3791/2056.

122. Bimbard, C., Takacs, F., Catarino, J. A., Fabre, J. M., Gupta, S., Lenzi, S. C., Melin, M. D., O’Neill, N., Orsolic, I., Robacha, M., et al. (2025). An adaptable, reusable, and light implant for chronic Neuropixels probes. eLife, 13, RP98522. DOI: 10.7554/eLife.98522.3.

123. Matsui, T. K., Tsuru, Y., Hasegawa, K., and Kuwako, K.-i. (2021). Vascularization of human brain organoids. Stem Cells, 39 (8), 1017–1024. DOI: 10.1002/stem.3368.

124. Gonzalez-Ferrer, J., Lehrer, J., O’Farrell, A., Paten, B., Teodorescu, M., Haussler, D., Jonsson, V. D., and Mostajo-Radji, M. A. (2024). SIMS: A deep-learning label transfer tool for single-cell RNA sequencing analysis. Cell Genom. 4 (6), 100581. DOI: 10.1016/j.xgen.2024.100581.

125. Bhaduri, A., Andrews, M. G., Mancia Leon, W., Jung, D., Shin, D., Allen, D., Jung, D., Schmunk, G., Haeussler, M., Salma, J., et al. (2020). Cell stress in cortical organoids impairs molecular subtype specification. Nature, 578 (7793), 142–148. DOI: 10.1038/s41586-020-1962-0.

126. Jackson, J., Ayzenshtat, I., Karnani, M. M., and Yuste, R. (2016). VIP+ interneurons control neocortical activity across brain states. J. Neurophysiol. 115 (6), 3008–3017. DOI: 10.1152/jn.01124.2015.

127. Paolicelli, R. C., Bolasco, G., Pagani, F., Maggi, L., Scianni, M., Panzanelli, P., Giustetto, M., Ferreira, T. A., Guiducci, E., Dumas, L., et al. (2011). Synaptic pruning by microglia is necessary for normal brain development. Science, 333 (6048), 1456–1458. DOI: 10.1126/science.1202529.

128. Tanveer, M. S., Patel, D., Schweiger, H. E., Abu-Bonsrah, K. D., Watmuff, B., Azadi, A., Pryshchep, S., Narayanan, K., Puleo, C., Natarajan, K., et al. (2025). Starting a Synthetic Biological Intelligence Lab from Scratch. Patterns, 6 (5), 101232. DOI: 10.1016/j.patter.2025.101232.

129. Scott, D. N. and Frank, M. J. (2023). Adaptive control of synaptic plasticity integrates micro- and macroscopic network function. Neuropsychopharmacol. 48 (1), 121–144. DOI: 10.1038/s41386-022-01374-6.

130. Osaki, T., Duenki, T., Chow, S. Y. A., Ikegami, Y., Beaubois, R., Levi, T., Nakagawa-Tamagawa, N., Hirano, Y., and Ikeuchi, Y. (2024). Complex activity and short-term plasticity of human cerebral organoids reciprocally connected with axons. Nat. Commun. 15 (1), 2945. DOI: 10.1038/s41467-024-46787-7.

131. Clark, I. C., Fontanez, K. M., Meltzer, R. H., Xue, Y., Hayford, C., May-Zhang, A., D’Amato, C., Osman, A., Zhang, J. Q., Hettige, P., et al. (2023). Microfluidics-free single-cell genomics with templated emulsification. Nat. Biotechnol. 41 (11), 1557–1566. DOI: 10.1038/s41587-023-01685-z.

132. Pachitariu, M., Steinmetz, N. A., Kadir, S. N., Carandini, M., and Harris, K. D. (2016). Fast and accurate spike sorting of high-channel count probes with KiloSort. 30th Conference on Neural Information Processing Systems (NIPS 2016), Barcelona, Spain. 29.

133. Hill, D. N., Mehta, S. B., and Kleinfeld, D. (2011). Quality Metrics to Accompany Spike Sorting of Extracellular Signals. J. Neurosci. 31 (24), 8699–8705. DOI: 10.1523/JNEUROSCI.0971-11.2011.

134. Mayer, S., Chen, J., Velmeshev, D., Mayer, A., Eze, U. C., Bhaduri, A., Cunha, C. E., Jung, D., Arjun, A., Li, E., et al. (2019). Multimodal Single-Cell Analysis Reveals Physiological Maturation in the Developing Human Neocortex. Neuron, 102 (1), 143–158.e7. DOI: 10.1016/j.neuron.2019.01.027.

135. Rossant, C., Hunter, M., Steinmetz, N., Wallace, M., Spacek, M., Gestes, C., McKenzie, Z., Nolan, C., Buccino, A., Zapp, S., et al. phy: Interactive Visualization and Manual Spike Sorting of Large-Scale Ephys Data [Python]. The Cortical Processing Laboratory at UCL. (2023).

136. Hao, Y., Stuart, T., Kowalski, M. H., Choudhary, S., Hoffman, P., Hartman, A., Srivastava, A., Molla, G., Madad, S., Fernandez-Granda, C., et al. (2024). Dictionary Learning for Integrative, Multimodal and Scalable Single-Cell Analysis. Nat. Biotechnol. 42 (2), 293–304. DOI: 10.1038/s41587-023-01767-y.

137. Heaton, H., Talman, A. M., Knights, A., Imaz, M., Gaffney, D. J., Durbin, R., Hemberg, M., and Lawniczak, M. K. N. (2020). Souporcell: Robust Clustering of Single-Cell RNA-seq Data by Genotype Without Reference Genotypes. Nat. Methods, 17 (6), 615–620. DOI: 10.1038/s41592-020-0820-1.

138. McGinnis, C. S., Murrow, L. M., and Gartner, Z. J. (2019). DoubletFinder: Doublet Detection in Single-Cell RNA Sequencing Data Using Artificial Nearest Neighbors. Cell Syst. 8 (4), 329–337.e4. DOI: 10.1016/j.cels.2019.03.003.

139. Korsunsky, I., Millard, N., Fan, J., Slowikowski, K., Zhang, F., Wei, K., Baglaenko, Y., Brenner, M., Loh, P.-r., and Raychaudhuri, S. (2019). Fast, Sensitive and Accurate Integration of Single-Cell Data with Harmony. Nat. Methods, 16 (12), 1289–1296. DOI: 10.1038/s41592-019-0619-0.

140. Lause, J., Berens, P., and Kobak, D. (2021). Analytic Pearson Residuals for Normalization of Single-Cell RNA-seq UMI Data. Genome Biol. 22 (1), 258. DOI: 10.1186/s13059-021-02451-7.

141. Choudhary, S. and Satija, R. (2022). Comparison and Evaluation of Statistical Error Models for scRNA-seq. Genome Biol. 23 (1), 27. DOI: 10.1186/s13059-021-02584-9.

142. Becht, E., McInnes, L., Healy, J., Dutertre, C.-A., Kwok, I. W. H., Ng, L. G., Ginhoux, F., and Newell, E. W. (2019). Dimensionality Reduction for Visualizing Single-Cell Data Using UMAP. Nat. Biotechnol. 37 (1), 38–44. DOI: 10.1038/nbt.4314.

143. Yao, Z., Velthoven, C. T. J. v., Nguyen, T. N., Goldy, J., Sedeno-Cortes, A. E., Baftizadeh, F., Bertagnolli, D., Casper, T., Chiang, M., Crichton, K., et al. (2021). A Taxonomy of Transcriptomic Cell Types Across the Isocortex and Hippocampal Formation. Cell, 184 (12), 3222–3241.e26. DOI: 10.1016/j.cell.2021.04.021.

144. Speir, M. L., Bhaduri, A., Markov, N. S., Moreno, P., Nowakowski, T. J., Papatheodorou, I., Pollen, A. A., Raney, B. J., Seninge, L., Kent, W. J., et al. (2021). UCSC Cell Browser: visualize your single-cell data. Bioinformatics, 37 (23), 4578–4580. DOI: 10.1093/bioinformatics/btab503.

145. Hagberg, A. A., Schult, D. A., and Swart, P. J. (2008). Exploring network structure, dynamics, and function using NetworkX. Proceedings of the 7th Python in Science Conference (SciPy2008), 11–15.

