## Supplemental Figures and Tables for "Self-Organizing Neural Networks in Organoids Reveal Principles of Forebrain Circuit Assembly"

### Supplemental Information

### Supplemental Figures

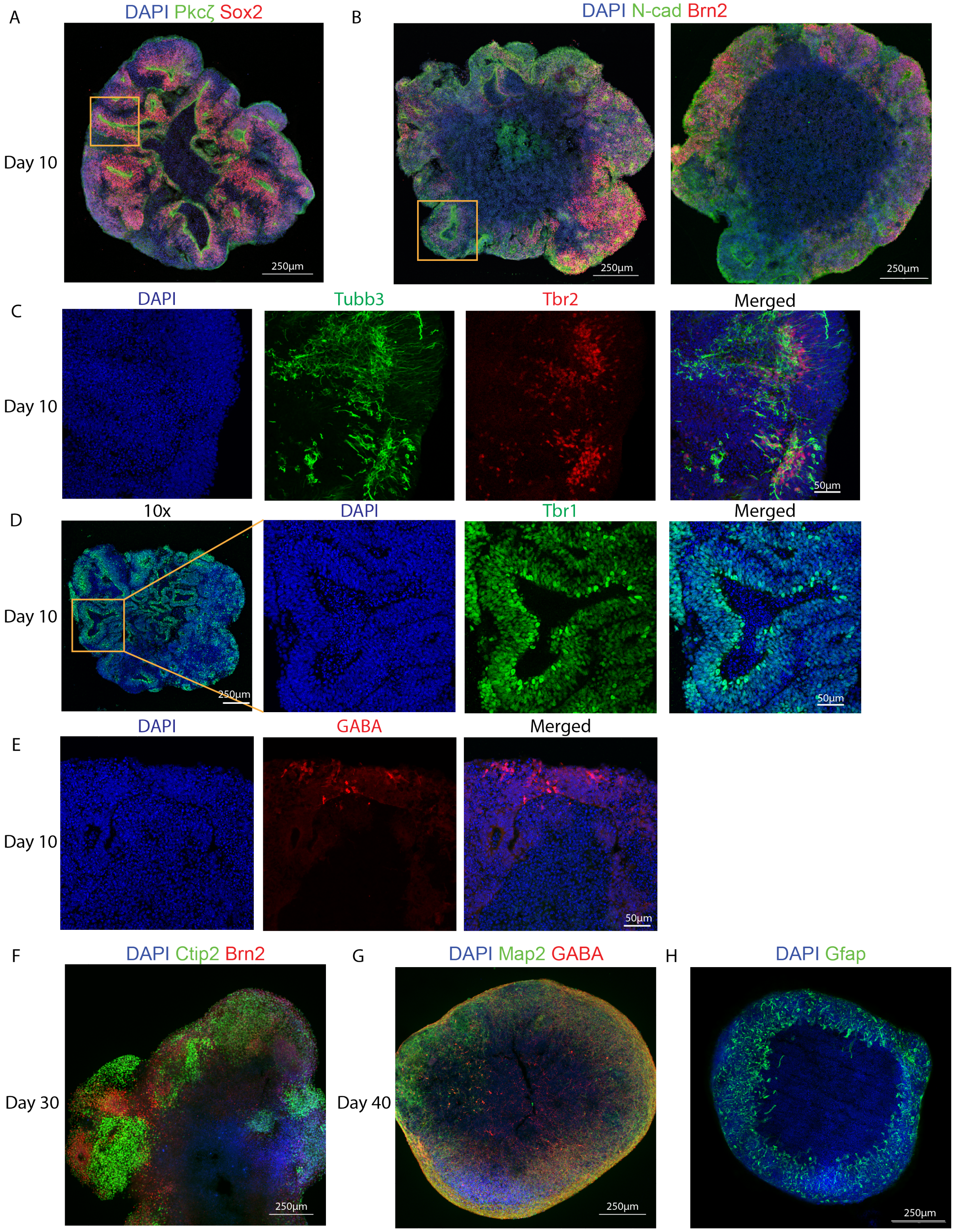

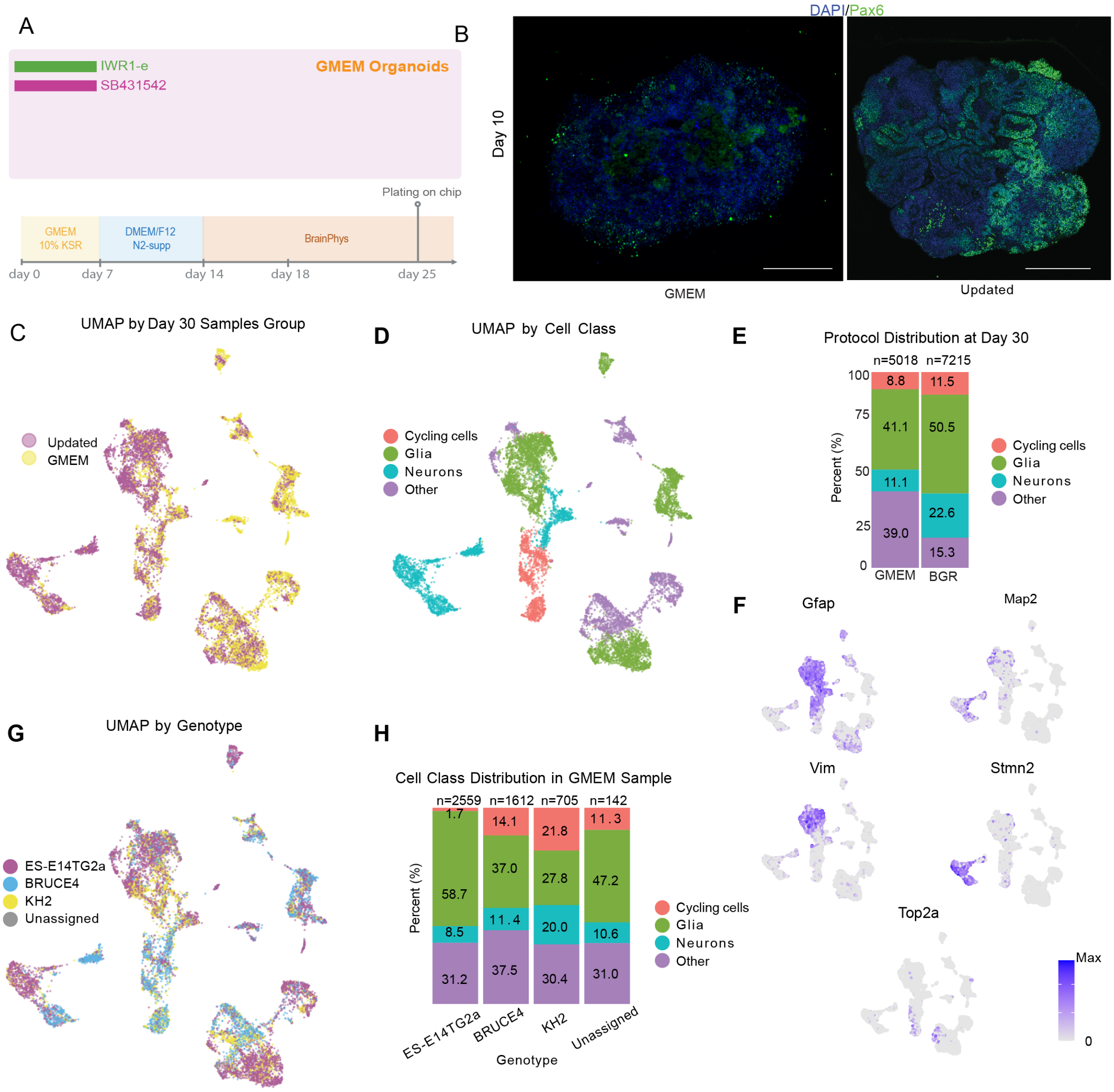


**Figure S1: Comparison of GMEM-based and Braingeneers (BGR) protocols in dorsal forebrain organoid development, related to Figure 1. (A) Schematic of the GMEM-based protocol.**
**(B) Representative IHC images of day 10 organoids generated using the GMEM-based (left) and BGR (right) protocols, stained for DAPI (blue) and Pax6 (green). Scale bars: 250 µm.**
**(C) UMAP visualization of single-cell RNA sequencing data from day 30 samples, colored by protocol (BGR and GMEM-based).**
**(D) UMAP visualization showing cell class distribution (Cycling cells, Glia, Neurons, and Other).**
**(E) Stacked bar plot comparing cell class distributions between GMEM-based (n = 5,018 cells) and BGR (n = 7,215 cells) protocols at day 30. The BGR protocol shows an increase in neuronal populations and a decrease in off-target ”Other” cells.**
**(F) UMAP plots displaying expression of key marker genes (Gfap, Map2, Vim, Stmn2, and Top2a).**
**(G) UMAP visualization colored by genotype (ES-E14TG2a, BRUCE4, KH2, and Unassigned).**
**(H) Detailed cell class distribution across different genotypes in GMEM-based samples.**

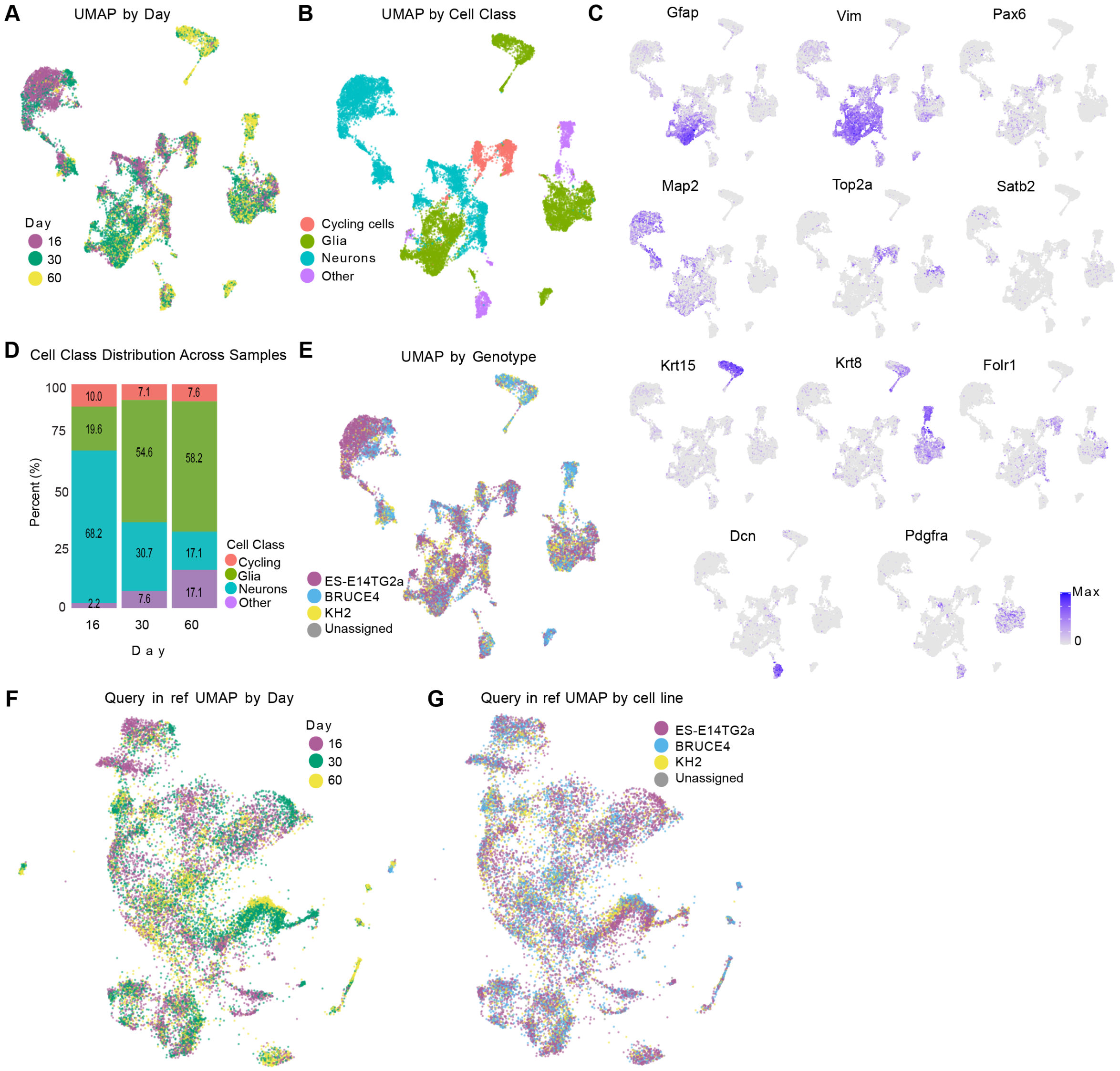


**Figure S2: Developmental progression and cellular composition of BGR protocol dorsal forebrain organoids, related to Figure 1.**
**(A) UMAP by time point visualization of dorsal forebrain cells (days 16, 30, and 60).**
**(B) UMAP representation highlighting major cell classes (Cycling cells, Glia, Neurons, and Other).**
**(C) Stacked bar plot showing the proportional distribution of cell classes at each time-point.**
**(D) UMAP visualization of cells grouped by genotype (ES-E14TG2a, BRUCE4, KH2, and Unassigned).**
**(E) Feature plot displaying expression levels of canonical marker genes used for cell type classification (Gfap, Vim, Pax6, Map2, Top2a, Satb2, Krt15, Krt8, Folr1, Dcn, and Pdgfra). Expression intensity is indicated by a color gradient from gray (low) to purple (high).**

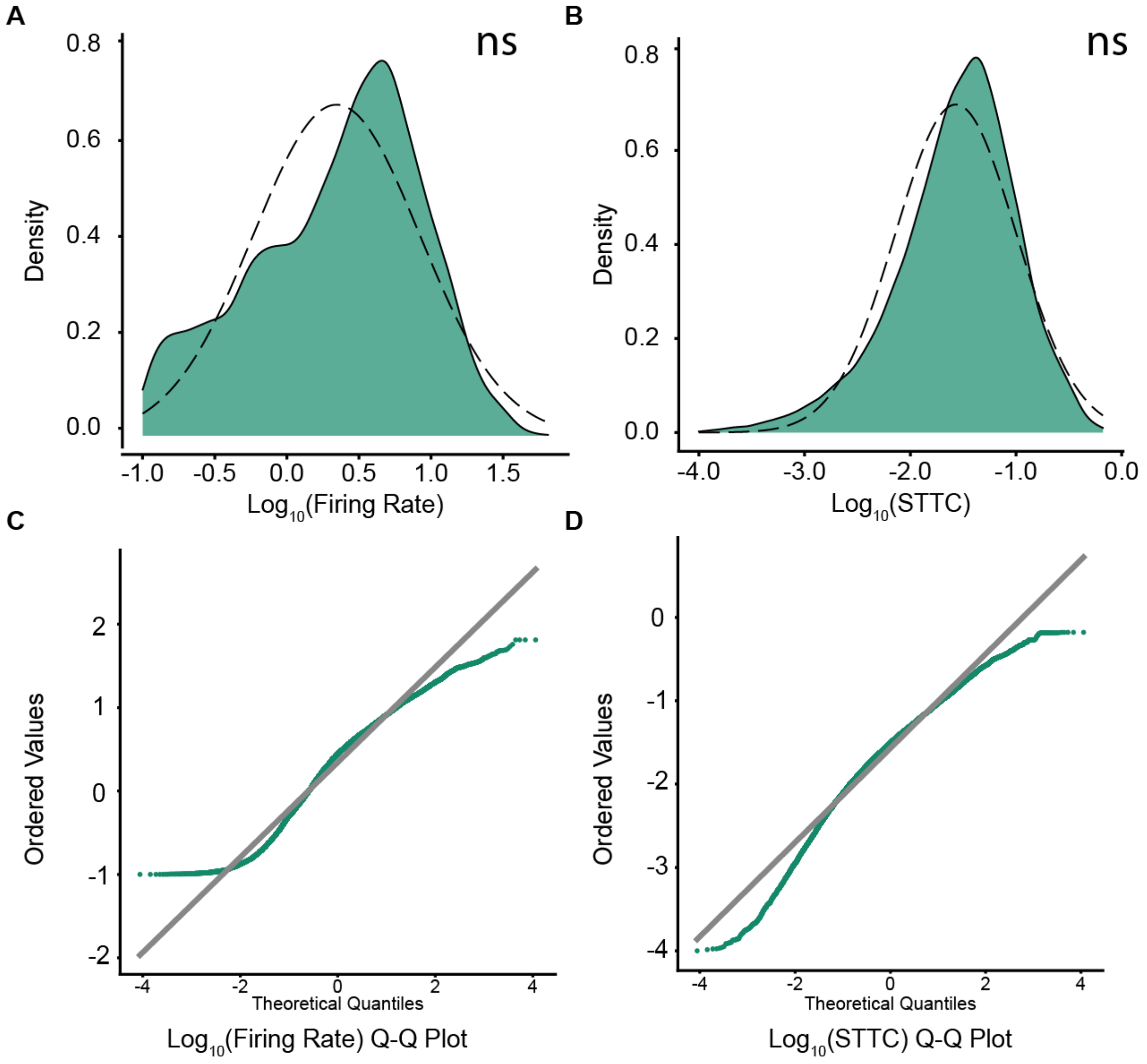


**Figure S3: Log normal, related to Figure 2.**
**(A-B) Log-normal distribution of log transformed mean firing rate distribution (A) and log transformed mean STTC (B) (green) with theoretical normal distribution (dashed line). (C-D) Q-Q Plots showing that both log transformed mean firing rate distribution (A) and log transformed mean STTC follow a log-normal distribution**
**ns = not significant. Kolmogorov–Smirnov test**

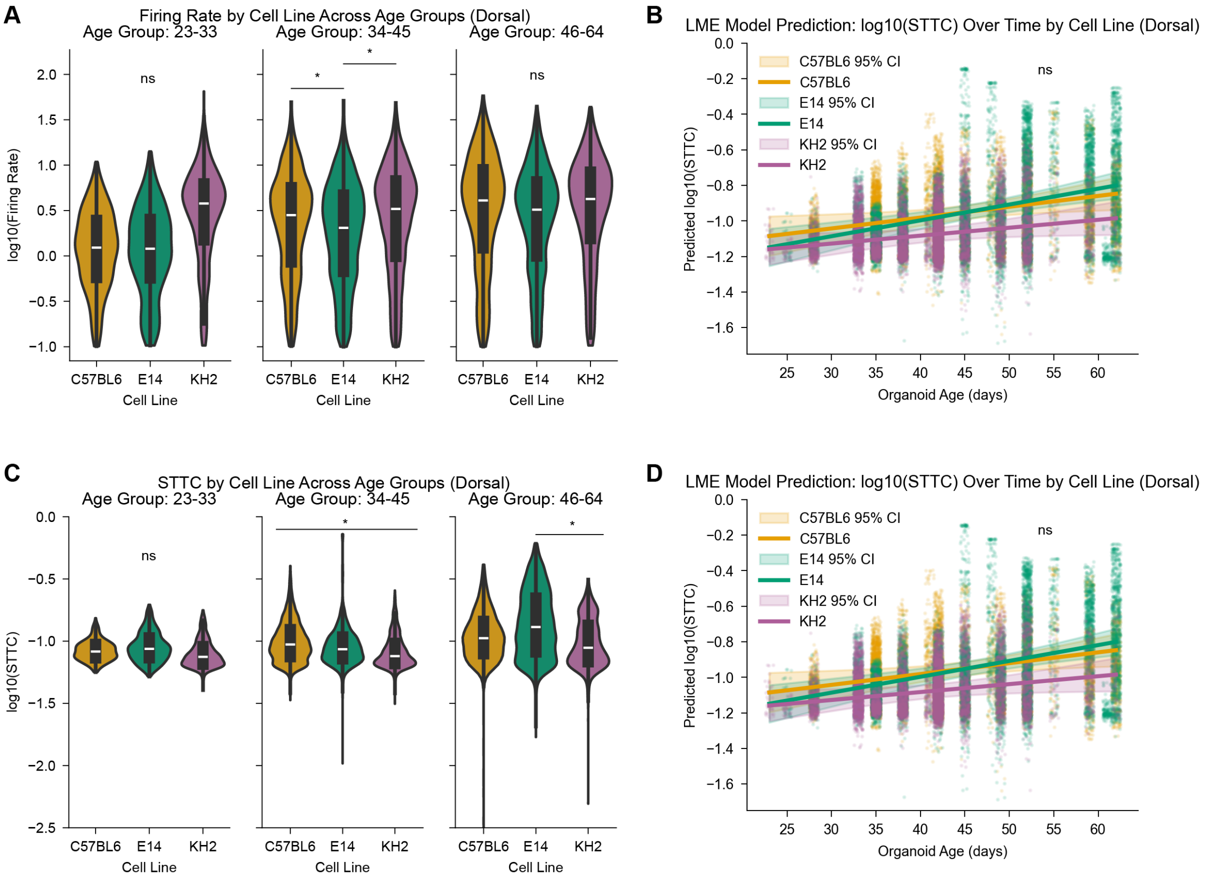


**Figure S4: Similar DF electrophysiological distributions across cell lines, related to Figure 2.**
**(A) Violin plots showing log10(Firing Rate) distributions by cell line (C57BL6, E14, and KH2) across three age groups (23-33, 34-45, and 46-64) in DF organoids. (B) Linear mixed effects (LME) model predictions of log10(STTC) over development by cell line for dorsal organoids. (C) Violin plots showing log10(STTC) distributions by cell line across the same three age groups in DF organoids. (D) LME model prediction of log10(STTC) over development by cell line for DF organoids, similar to panel B but with potentially different parameter settings.**
**ns = not significant, $p < 0.017$ (Bonferroni corrected). Mixed-effects model**

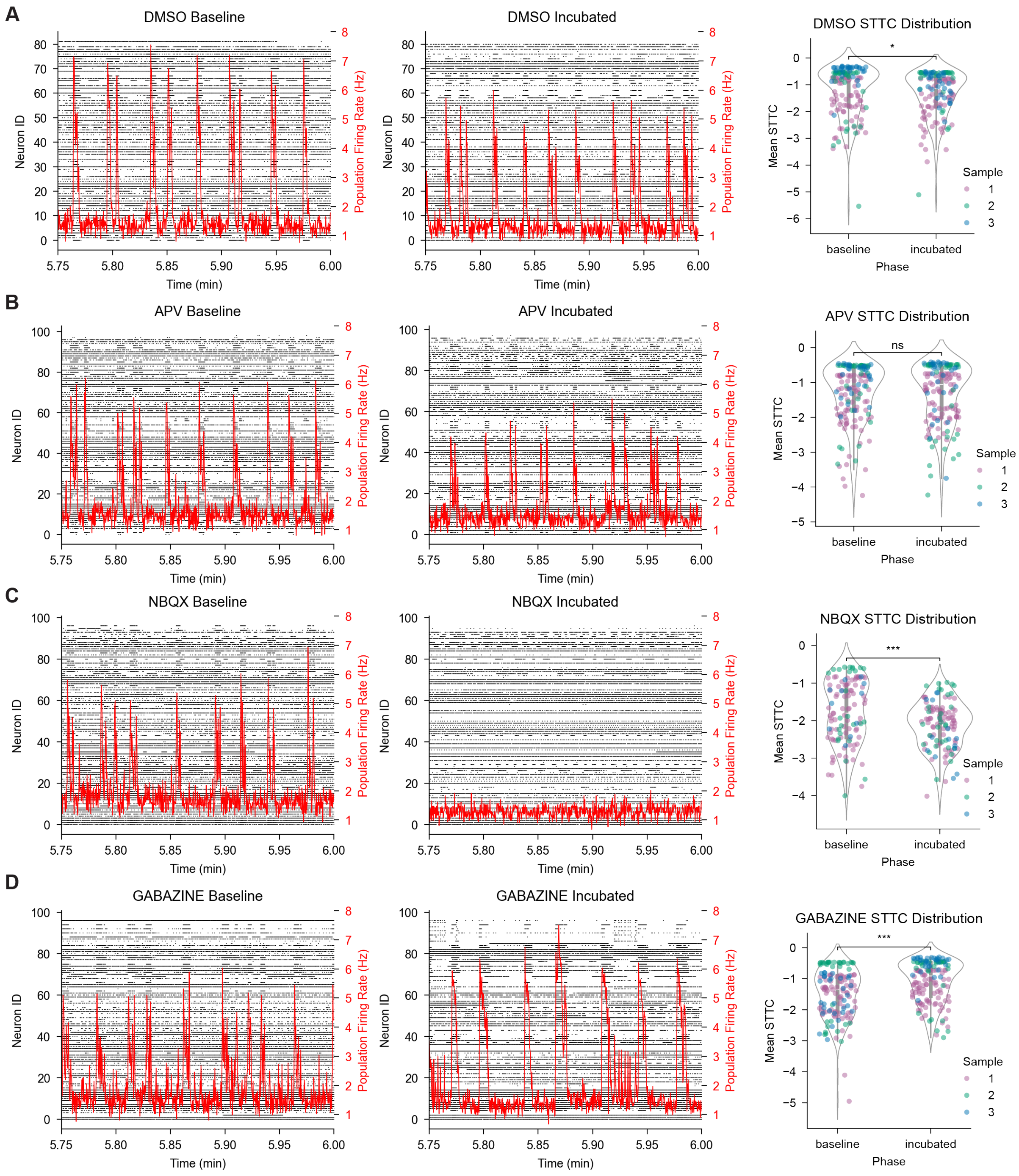


**Figure S5: Impact of pharmacological perturbation on overall neural connectivity, related to Figure 3.**
**(A) Raster plots of neural activity (gray) and population firing rate (red) during baseline (left) and post-drug incubation (middle) for a 15s window in DMSO vehicle control. Each row represents individual unit spike trains. (Right) STTC distribution showing the mean log STTC values across all 3 organoid samples. Organoid 1 is shown in purple, organoid 2 in green, and organoid 3 in yellow. Statistical significance indicated by asterisks: * $p < 0.05$, ** $p < 0.001$, *** $p < 0.0001$, ns: not significant.**
**(B) As in panel A but following treatment with NMDA antagonist APV.**
**(C) As in panel A but following treatment with AMPA/Kainate antagonist NBQX.**
**(D) As in panel A but following treatment with $GABA_A$ antagonist Gabazine.**

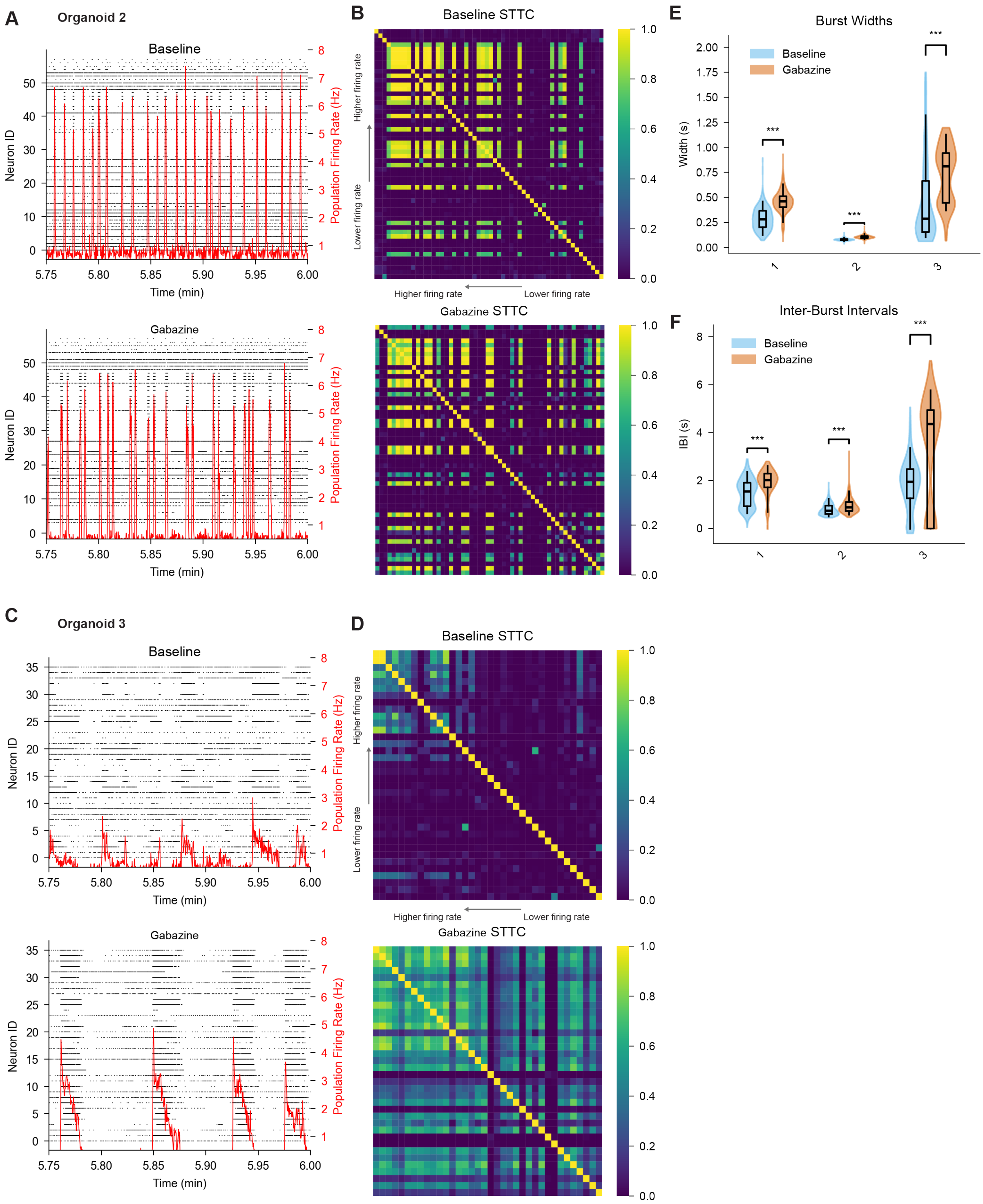


**Figure S6: Effects of GABA receptor antagonism on bursting activity in dorsal forebrain organoids, related to Figure 3.**
**(A) Representative raster plots from Sample 2 showing neural activity (gray) and population firing rate (red) during baseline (left) and post-Gabazine incubation (middle) over a 15s window. Each row represents a single-unit spike train.**
**(B) STTC matrices sorted by firing rate (ascending to descending). (Left) Baseline STTC matrix. Middle: STTC matrix post-Gabazine incubation. (right) Difference matrix showing STTC changes.**
**(C) Same as (A), but for Sample 3.**
**(D) Same as (B), but for Sample 3.**
**(E) Burst width across three organoids.**
**(F) Inter-burst interval across three organoids.**
**(E–F) Statistical comparison of baseline vs. Gabazine conditions. Statistical significance: * $p < 0.05$, ** $p < 0.001$, *** $p < 0.0001$; ns = not significant. Mann–Whitney U test.**

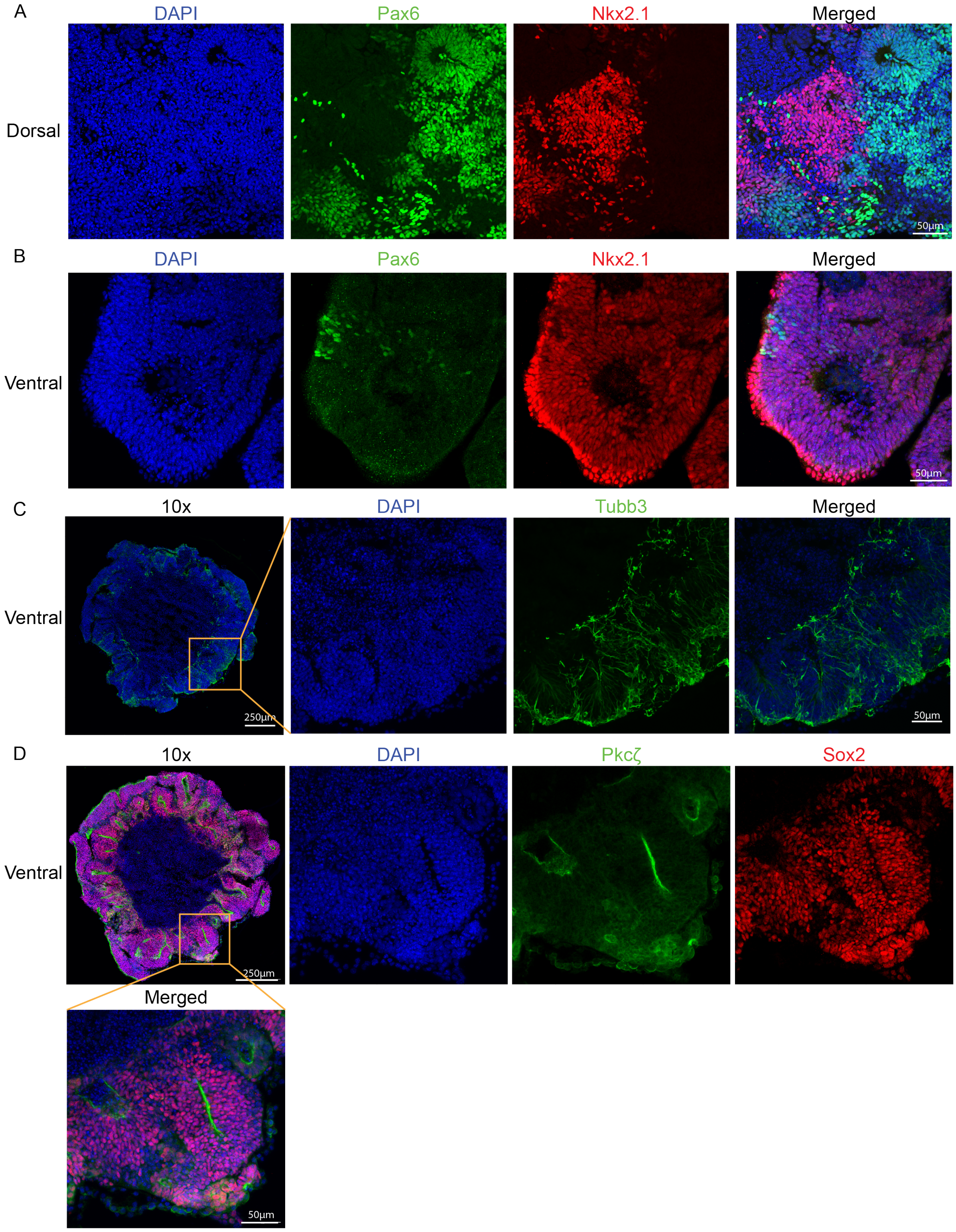


**Figure S7: Patterning marker expression in developing DF and VF organoids, related to Figure 4.**
**(A) High-magnification view of the DF from Figure 4C, showing Pax6 (green) and Nkx2.1 (red) staining with the merged image.**
**(B) High-magnification view of the VF from Figure 4C, showing Pax6 (green) and Nkx2.1 (red) staining with the merged image.**
**(C) Day 10 VF organoid. (Left) Low-magnification overview. (right) High-magnification view of Tubb3 (green) staining with the merged image.**
**(D) Day 10 VF organoid. (Left) Low-magnification overview. (right) High-magnification view of Pkcζ (green) and Sox2 (red).**
**All panels include DAPI nuclear counterstain (blue), with scale bars as indicated (50 or 250 μm).**

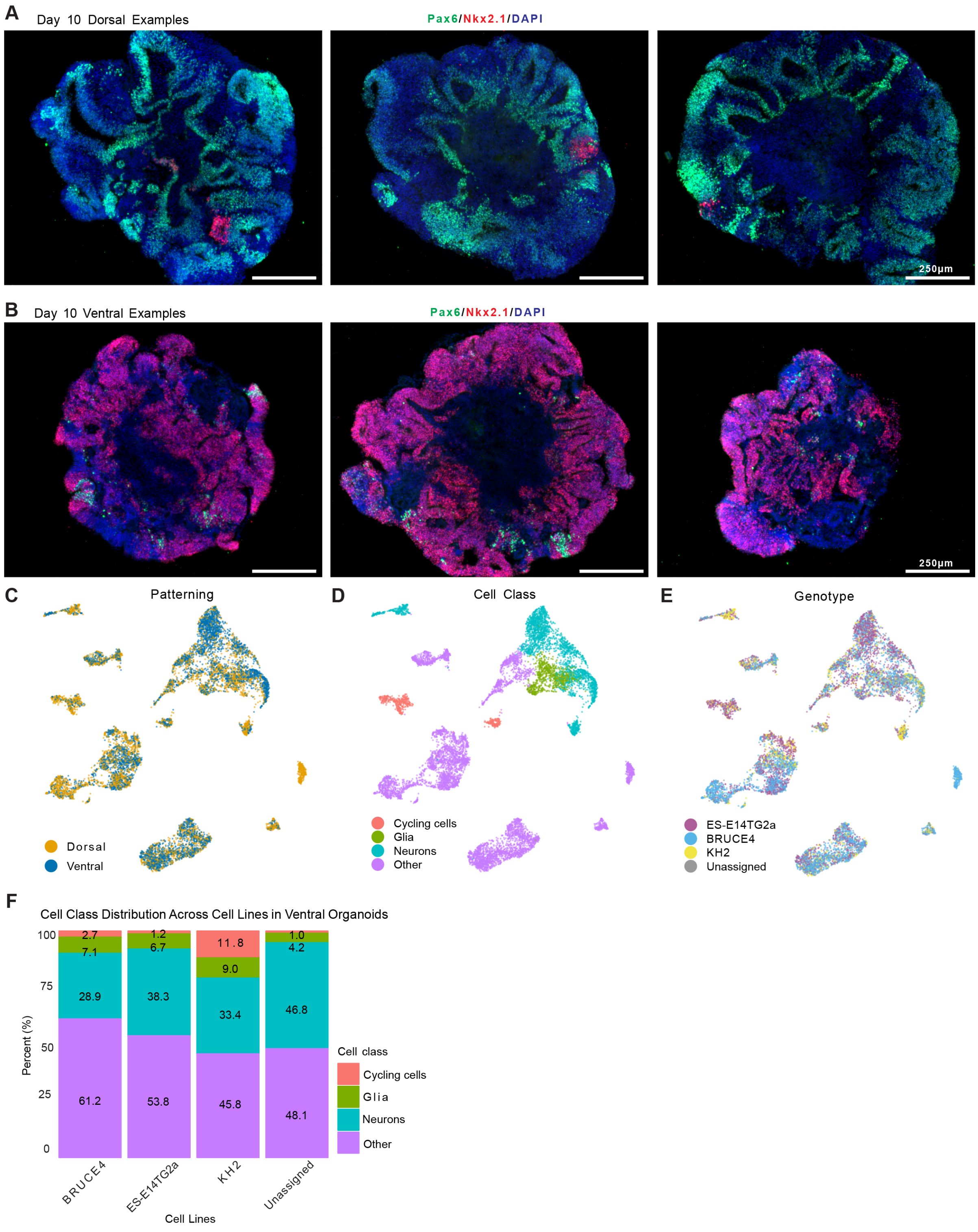


**Figure S8: Characterization of dorsal and ventral forebrain organoid development, related to Figure 4.**
**(A) IHC images of additional day 10 DF organoids stained for Pax6 (green) and Nkx2.1 (red). These organoids were used for quantifications in Figure 4D.**
**(B) IHC images of additional day 10 VF organoids stained for Pax6 (green) and Nkx2.1 (red). These organoids were used for quantifications in Figure 4D.**
**(C) UMAP visualization of scRNA-seq data colored by sample type (DF and VF).**
**(D) UMAP plot showing cell class (Cycling cells, Glia, Neurons, and Other).**
**(E) UMAP visualization colored by genotype (ES-E14TG2a, BRUCE4, KH2, and Unassigned).**
**(F) Stacked bar plot showing cell class distribution across genotypes in VF samples. Panels (A–B) include DAPI nuclear counterstain (blue) and scale bars as indicated (250 μm).**

Core-Periphery, Hubness & STTC through developement - DF Organoids
(Same chip: 23124)

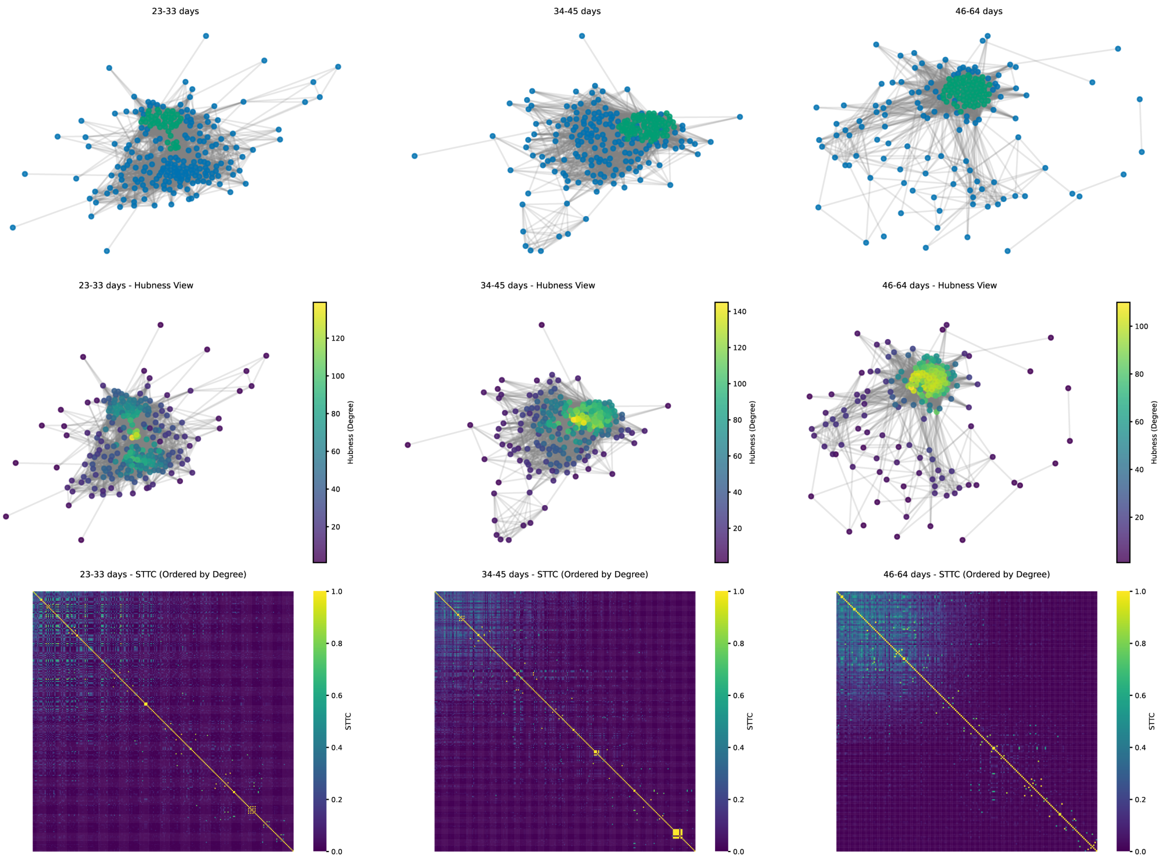


Core-Periphery, Hubness & STTC through developement - DF Organoids
(Same chip: 22710)

Core-Periphery, Hubness & STTC Matrix by Age (Dorsal)

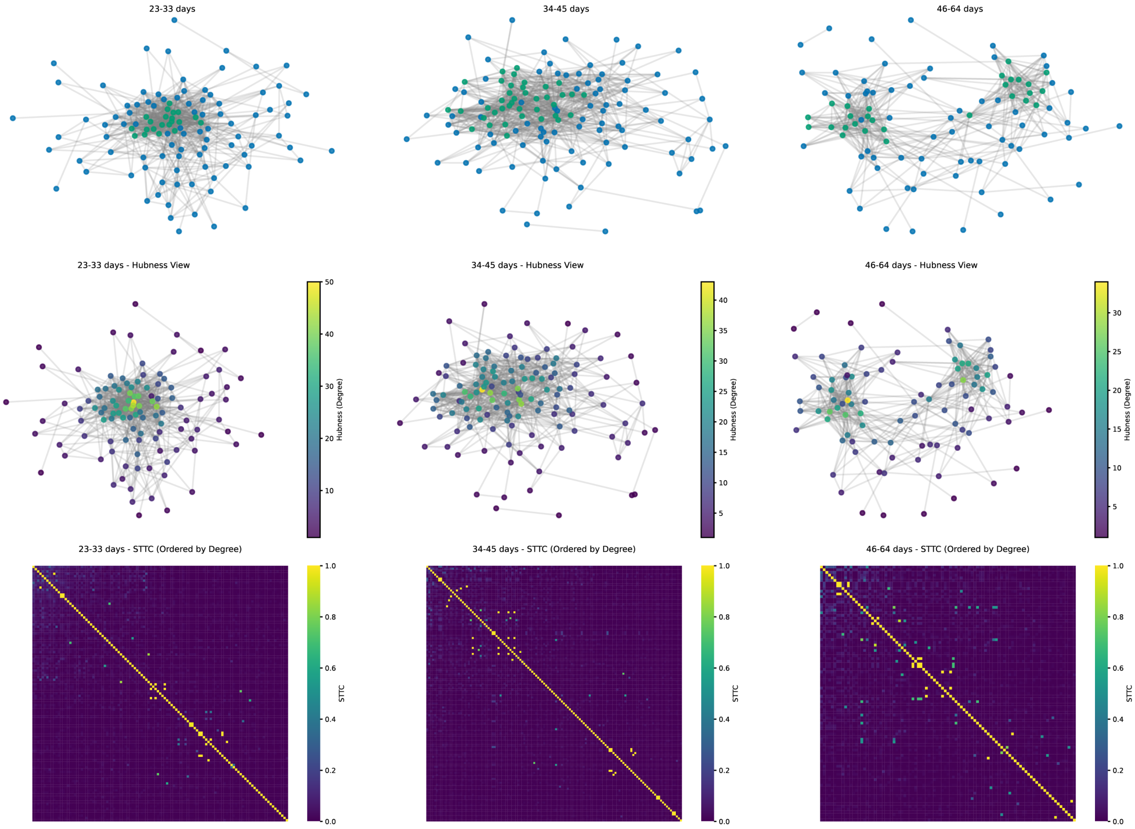

**Figure S9: Developmental changes in network connectivity of DF organoids tracked longitudinally, related to Figure 8.**
**(A) Longitudinal analysis of organoid chip 23120 shown at three developmental timepoints (23-33 days, 34-45 days, and 46-64 days). Upper panels display hubness visualizations where node colors represent hubness score. Lower panels show corresponding spike time tiling coefficient (STTC) matrices ordered by connection degree.**
**(B) Longitudinal analysis of organoid chip 23120 shown at three developmental timepoints (23-33 days, 34-45 days, and 46-64 days). Upper panels display hubness visualizations where node colors represent hubness score. Lower panels show corresponding spike time tiling coefficient (STTC) matrices ordered by connection degree.**

Core-Periphery, Hubness & STTC through developement - VF Organoids
(Same chip: 25136)

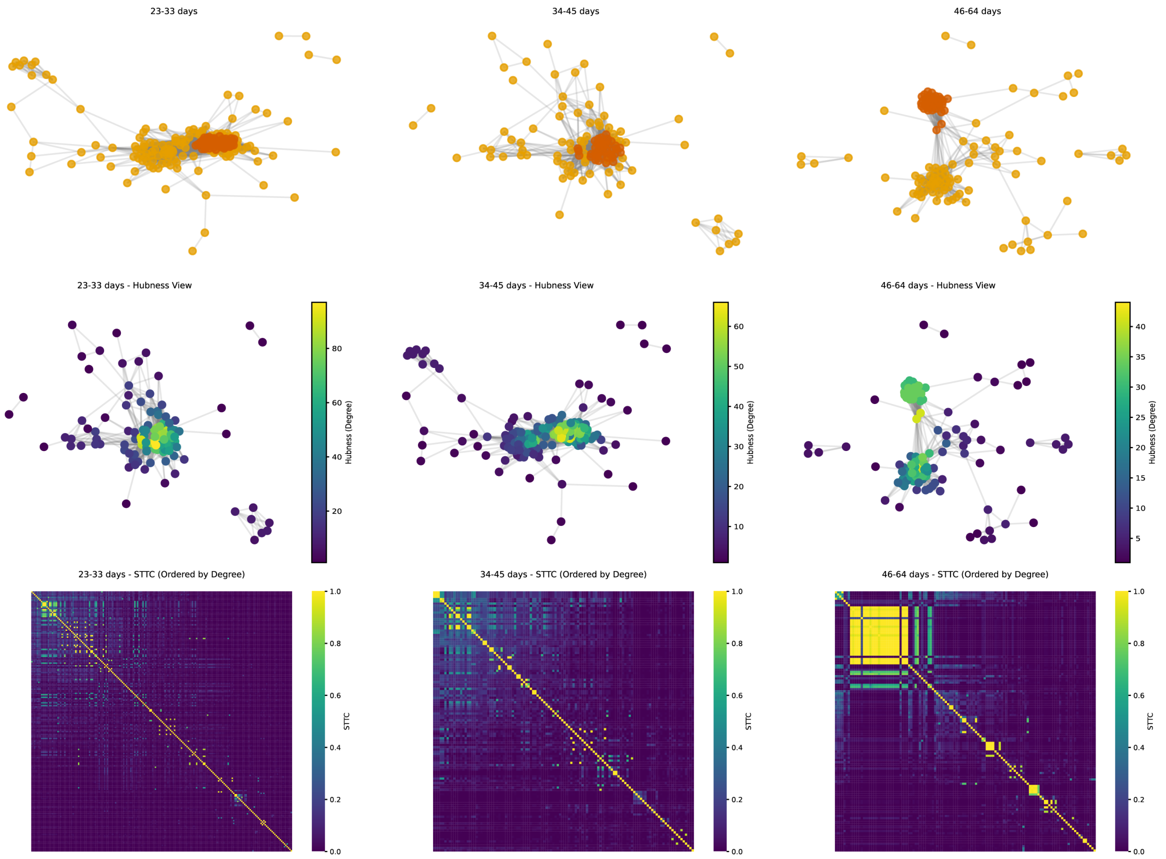


Core-Periphery, Hubness & STTC through developement - VF Organoids
(Same chip: 22064b)

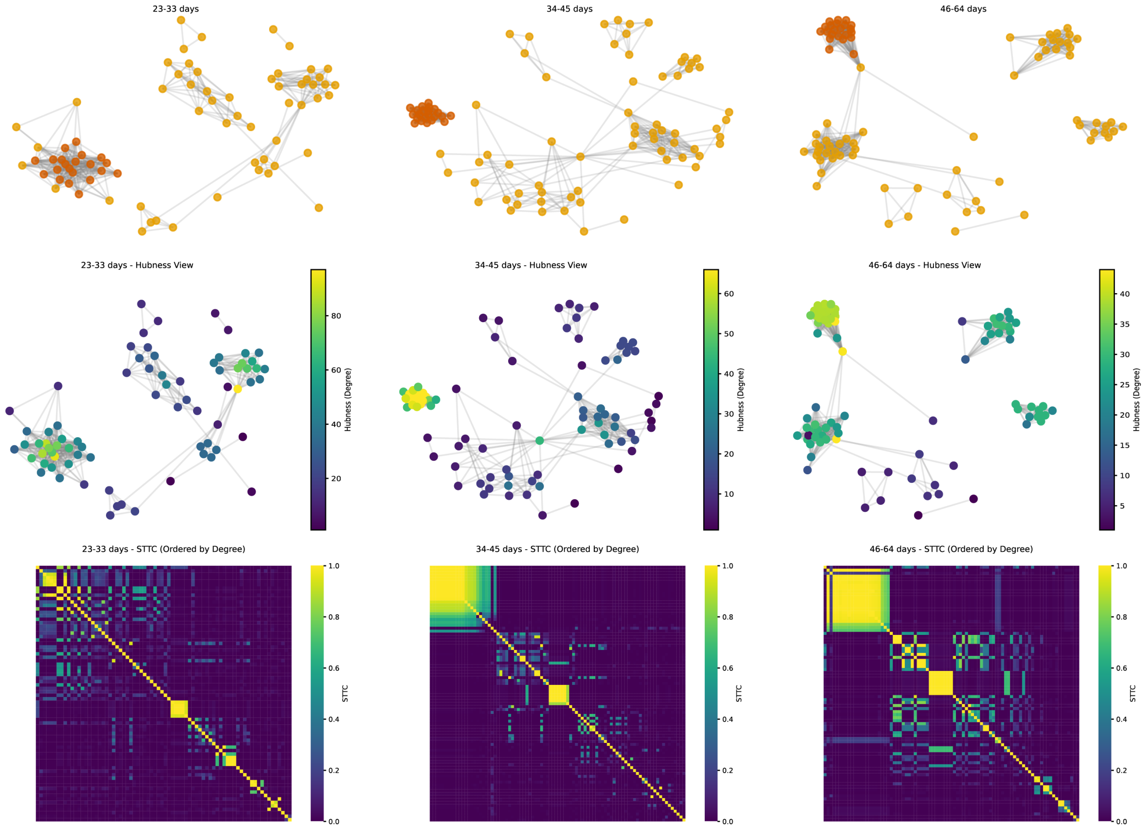

**Figure S10: Developmental changes in network connectivity of VF organoids tracked longitudinally, related to Figure 7.**
**(A) Longitudinal analysis of organoid chip 25136 shown at three developmental time-points (23-33 days, 34-45 days, and 46-64 days). Upper panels display hubness visualizations where node colors represent hubness score. Lower panels show corresponding spike time tiling coefficient (STTC) matrices ordered by connection degree.**
**(B) Longitudinal analysis of organoid chip 22064b shown at three developmental time-points (23-33 days, 34-45 days, and 46-64 days). Upper panels display hubness visualizations where node colors represent hubness score. Lower panels show corresponding spike time tiling coefficient (STTC) matrices ordered by connection degree.**

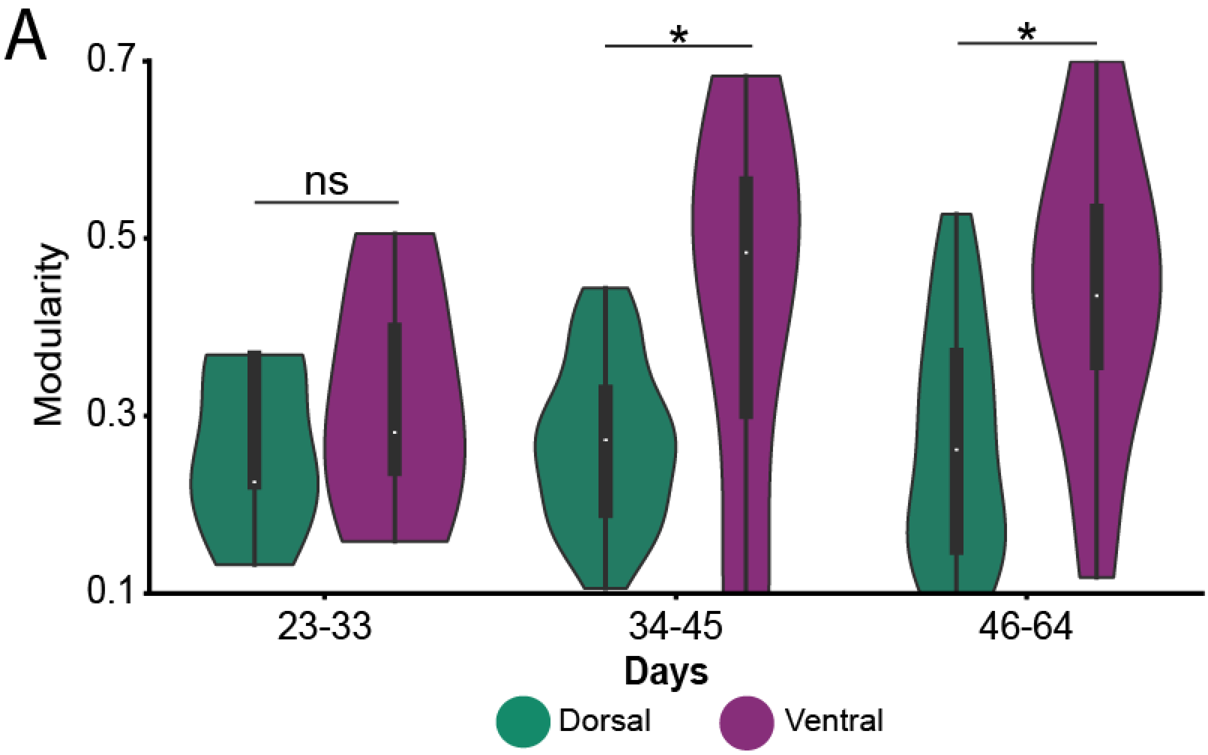


**Figure S11: Developmental differences in modularity metric between DF and VF organoids, related to Figure 7.**
**(A) Violin plots showing network modularity values for dorsal (green) and ventral (purple) organoids across three developmental time windows (23-33 days, 34-45 days, and 46-64 days). No significant difference in modularity is observed during early development (23-33 days, ns). However, ventral organoids display significantly higher modularity compared to dorsal organoids during both mid (34-45 days) and late (46-64 days) developmental stages (* indicates p<0.05 Mann–Whitney U test).**

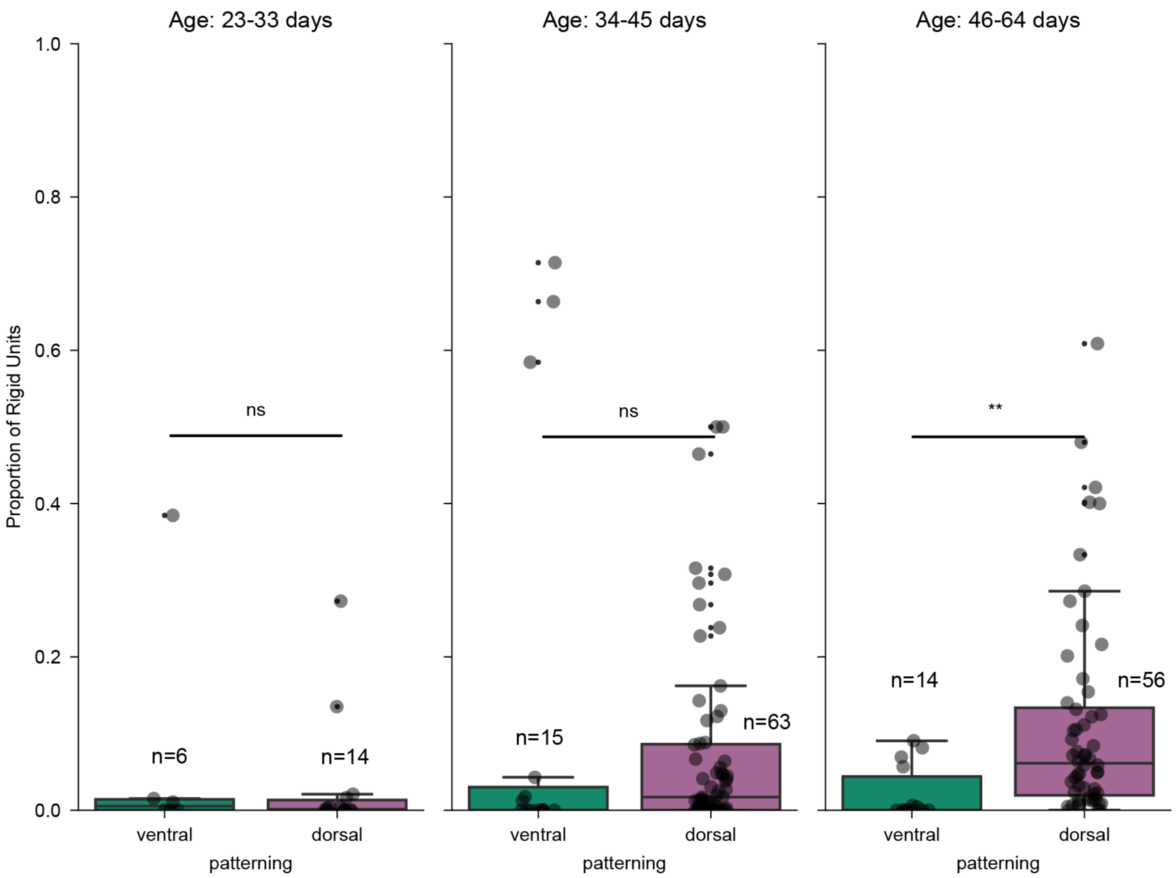


**Figure S12: Proportion of rigid units between development for DF and VF, related to Figure 9.**
**(A) Proportion of rigid units over development for VF (Left) and DF (Right).**
**ns = not significant, **p < 0.001, Mann-Whitney U test.**

#### Supplemental Tables

**Table S1: Statistical Comparison of Metrics Across Age Groups within Dorsal Organoids, related to Figure 2.**
**Firing rate (FR), Spike time tiling coefficient (STTC)**
**Mixed-effects model *p < 0.0167, **p < 0.0033, ***p < 0.00033 (Bonferroni corrected)**

| Metric | Pairwise Age Group Comparisons | | | | |
|---|---|---|---|---|---|
| | Age Groups | Diff | Std.Err. | z-value | p-value |
| $\log_{10}$ FR | 23-33 vs 34-45 | -0.202 | 0.053 | -3.825 | 0.000** |
| | 23-33 vs 46-64 | -0.271 | 0.054 | -5.063 | 0.000** |
| | 34-45 vs 46-64 | -0.069 | 0.035 | -1.954 | 0.051 |
| $\log_{10}$ STTC | 23-33 vs 34-45 | -0.087 | 0.040 | -2.202 | 0.028* |
| | 23-33 vs 46-64 | -0.190 | 0.040 | -4.728 | 0.000** |
| | 34-45 vs 46-64 | -0.103 | 0.027 | -3.841 | 0.000** |

**Table S2: Statistical Comparison of Metrics Across Cell Lines within Dorsal Organoids, related to Figure 2.**
**Firing rate (FR), Spike time tiling coefficient (STTC)**
**Mixed-effects model *p < 0.0167, **p < 0.0033, ***p < 0.00033 (Bonferroni corrected)**

| Metric | Age Group | Pairwise Cell Line Comparisons | | | | |
|---|---|---|---|---|---|---|
| | | Cell Lines | Diff | Std.Err. | z-value | p-value |
| $\log_{10}$ FR | 23-33 | C57BL6 vs E14 | 0.061 | 0.184 | 0.331 | 0.741 |
| | | C57BL6 vs KH2 | -0.119 | 0.165 | -0.722 | 0.470 |
| | | E14 vs KH2 | -0.180 | 0.129 | -1.399 | 0.162 |
| | 34-45 | C57BL6 vs E14 | 0.144 | 0.055 | 2.614 | 0.009** |
| | | C57BL6 vs KH2 | -0.049 | 0.050 | -0.973 | 0.331 |
| | | E14 vs KH2 | -0.193 | 0.056 | -3.464 | 0.001** |
| | 46-64 | C57BL6 vs E14 | 0.123 | 0.055 | 2.249 | 0.025* |
| | | C57BL6 vs KH2 | 0.007 | 0.063 | 0.109 | 0.913 |
| | | E14 vs KH2 | -0.116 | 0.061 | -1.907 | 0.056 |
| $\log_{10}$ STTC | 23-33 | C57BL6 vs E14 | 0.015 | 0.034 | 0.433 | 0.665 |
| | | C57BL6 vs KH2 | 0.046 | 0.031 | 1.509 | 0.131 |
| | | E14 vs KH2 | 0.031 | 0.024 | 1.315 | 0.188 |
| | 34-45 | C57BL6 vs E14 | 0.019 | 0.040 | 0.467 | 0.641 |
| | | C57BL6 vs KH2 | 0.115 | 0.036 | 3.154 | 0.002** |
| | | E14 vs KH2 | 0.096 | 0.041 | 2.370 | 0.018* |
| | 46-64 | C57BL6 vs E14 | -0.040 | 0.051 | -0.783 | 0.434 |
| | | C57BL6 vs KH2 | 0.106 | 0.058 | 1.819 | 0.069 |
| | | E14 vs KH2 | 0.146 | 0.057 | 2.586 | 0.010** |

**Table S3: Drug Effects on Firing Rate (FR), related to Figure 3.**
**Mixed-effects model, * Significant at p < 0.05. SEM = Standard Error of Mean**

| Drug | Baseline Mean (SEM) | Drug Mean (SEM) | p-value | Significant | Coefficient (SE) |
|---|---|---|---|---|---|
| APV | $21.94 \pm 1.04$ | $18.78 \pm 1.02$ | $0.0097$ | Yes* | $-0.080 \pm 0.108$ |
| DMSO | $22.03 \pm 1.19$ | $19.63 \pm 1.23$ | $0.2134$ | No | $-0.064 \pm 0.052$ |
| GABAZINE | $20.81 \pm 1.24$ | $23.06 \pm 1.46$ | $0.7893$ | No | $+0.017 \pm 0.063$ |
| NBQX | $16.64 \pm 1.19$ | $13.37 \pm 1.12$ | $0.0512$ | No | $-0.171 \pm 0.088$ |

* Significant at p < 0.05

**Table S4: Drug Effects on Spike Time Tiling Coefficient (STTC), related to Figure 3.**
**Mixed-effects model, * Significant at p < 0.05. SEM = Standard Error of Mean**

| Drug | Baseline Mean (SEM) | Drug Mean (SEM) | p-value | Significant | Coefficient (SE) |
|---|---|---|---|---|---|
| APV | $0.126 \pm 0.010$ | $0.132 \pm 0.011$ | $0.4584$ | No | $-0.080 \pm 0.108$ |
| DMSO | $0.168 \pm 0.014$ | $0.116 \pm 0.010$ | $0.5245$ | No | $-0.066 \pm 0.104$ |
| GABAZINE | $0.107 \pm 0.011$ | $0.188 \pm 0.014$ | $< 0.0001$ | Yes* | $+0.352 \pm 0.084$ |
| NBQX | $0.063 \pm 0.010$ | $0.022 \pm 0.004$ | $0.0033$ | Yes* | $-0.324 \pm 0.110$ |

**Table S5: Statistical Comparison of FR and STTC Across Age Groups within VF Organoids, related to Figure 5.**
**Firing rate (FR), Spike time tiling coefficient (STTC)**
**Mixed-effects model *p < 0.0167, **p < 0.0033, ***p < 0.00033 (Bonferroni corrected)**

| Metric | Pairwise Age Group Comparisons | | | | |
|---|---|---|---|---|---|
| | Age Groups | Diff | Std.Err. | z-value | p-value |
| $\log_{10}$ FR | 23-33 vs 34-45 | -0.292 | 0.090 | -3.259 | 0.001** |
| | 23-33 vs 46-64 | -0.271 | 0.088 | -3.089 | 0.002** |
| | 34-45 vs 46-64 | 0.021 | 0.067 | 0.309 | 0.758 |
| $\log_{10}$ STTC | 23-33 vs 34-45 | -0.100 | 0.048 | -2.106 | 0.035* |
| | 23-33 vs 46-64 | -0.056 | 0.047 | -1.193 | 0.233 |
| | 34-45 vs 46-64 | 0.045 | 0.036 | 1.251 | 0.211 |

**Table S6: Statistical Comparison of FR and STTC Between DF and VF Organoids Across Age Groups, related to Figure 5.**
**Firing rate (FR), Spike time tiling coefficient (STTC)**
**Mixed-effects model, * Significant at p < 0.05.**

| Age Group | Metric | Coefficient | | Significance | | |
|---|---|---|---|---|---|---|
| | | Estimate | Std.Err. | z-value | p-value | Significant |
| 23-33 days | FR | -0.077 | 0.093 | -0.828 | 0.408 | No |
| | STTC | 0.032 | 0.022 | 1.460 | 0.144 | No |
| 34-45 days | FR | 0.011 | 0.051 | 0.222 | 0.824 | No |
| | STTC | 0.046 | 0.036 | 1.261 | 0.207 | No |
| 46-64 days | FR | -0.078 | 0.053 | -1.459 | 0.145 | No |
| | STTC | -0.102 | 0.043 | -2.344 | 0.019* | Yes |

**Table S7: Age Group Comparisons for Small World Metrics in DF organoids, related to Figure 6.**
**Mixed-effects model *p < 0.0167, **p < 0.0033, ***p < 0.00033 (Bonferroni corrected)**

| **Metric** | **Measure** | **Age Group Comparisons** | | |
|---|---|---|---|---|
| | | 23-33 vs 34-45 days | 34-45 vs 46-64 days | 23-33 vs 46-64 days |
| **Small World (S)** | p-value | **< 0.001*****| **4.14e-06***** | **< 0.001***** |
| | Effect Size | 0.178 | 0.0208 | 0.199 |
| **Clustering (C_norm)** | p-value | **< 0.001***** | **1.68e-20***** | **< 0.001***** |
| | Effect Size | 0.319 | 0.0701 | 0.389 |
| **Path Length (L_norm)** | p-value | **< 0.001***** | **1.22e-09***** | **< 0.001***** |
| | Effect Size | 0.046 | 0.0101 | 0.056 |

**Table S8: Age Group Comparisons for Small World Metrics in VF organoids, related to Figure 6.**
**Mixed-effects model, *p < 0.0167, **p < 0.0033, ***p < 0.00033 (Bonferroni corrected)**

| Metric | Measure | Age Group Comparisons | | |
|---|---|---|---|---|
| | | 23-33 vs 34-45 days | 34-45 vs 46-64 days | 23-33 vs 46-64 days |
| **Small World (S)** | p-value | **0.003**** | **4.73e-05***** | **< 0.001***** |
| | Effect Size | 0.162 | 0.0727 | 0.235 |
| **Clustering (C_norm)** | p-value | **< 0.001***** | **< 0.001***** | **< 0.001***** |
| | Effect Size | 0.202 | 0.3615 | 0.563 |
| **Path Length (L_norm)** | p-value | **< 0.001***** | **< 0.001***** | **< 0.001***** |
| | Effect Size | 0.042 | 0.0377 | 0.080 |

**Table S9: Patterning Comparisons (DF vs VF) by Age Group for Small World metrics, related to Figure 6.**
**Mixed-effects model *p < 0.05, **p < 0.01, ***p < 0.001.**

| Metric | Measure | Age Groups | | |
|---|---|---|---|---|
| | | 23-33 days | 34-45 days | 46-64 days |
| **Small World (S)** | p-value | **< 0.001***** | **< 0.001***** | **< 0.001***** |
| | Dorsal Mean | 2.455 | 2.633 | 2.654 |
| | Ventral Mean | 3.137 | 3.300 | 3.372 |
| **Clustering (C_norm)** | p-value | **< 0.001***** | **< 0.001***** | **< 0.001***** |
| | Dorsal Mean | 2.708 | 3.027 | 3.097 |
| | Ventral Mean | 3.723 | 3.925 | 4.286 |
| **Path Length (L_norm)** | p-value | **< 0.001***** | **< 0.001***** | **< 0.001***** |
| | Dorsal Mean | 1.103 | 1.149 | 1.159 |
| | Ventral Mean | 1.206 | 1.248 | 1.285 |

**Table S10: Statistical Comparison of modularity Across DF and VF age ranges, related to Figure 7.**
**Mixed-effects model *p < 0.0167, **p < 0.0033, ***p < 0.00033 (Bonferroni corrected).**

| Comparison | p-value | Significant | Median | |
|---|---|---|---|---|
| | | | Group 1 | Group 2 |
| Dorsal vs Ventral (23-33 days) | 0.2977 | No | 0.2257 | 0.2814 |
| Dorsal vs Ventral (34-45 days) | 0.0066 | Yes** | 0.2730 | 0.4836 |
| Dorsal vs Ventral (46-64 days) | 0.0019 | Yes** | 0.2620 | 0.4349 |
| Dorsal: 23-33 days vs 34-45 days | 0.6612 | No | 0.2257 | 0.2730 |
| Dorsal: 34-45 days vs 46-64 days | 0.7433 | No | 0.2730 | 0.2620 |
| Dorsal: 23-33 days vs 46-64 days | 0.9405 | No | 0.2257 | 0.2620 |
| Ventral: 23-33 days vs 34-45 days | 0.2976 | No | 0.2814 | 0.4836 |
| Ventral: 34-45 days vs 46-64 days | 0.9420 | No | 0.4836 | 0.4349 |
| Ventral: 23-33 days vs 46-64 days | 0.2075 | No | 0.2814 | 0.4349 |

**Table S11: Comparison of Rigid Unit Proportion Between DF and VF Organoids by Age Group, related to Figure 9**
**Mann-Whitney U test, * Significant at p < 0.05**

| Age Group (days) | DF Mean (± SD) | VF Mean (± SD) | U-statistic | p-value | Significant |
|---|---|---|---|---|---|
| 23-33 | $0.033 \pm 0.078$ | $0.068 \pm 0.155$ | 39.5 | 0.8601 | No |
| 34-45 | $0.076 \pm 0.124$ | $0.136 \pm 0.270$ | 582.5 | 0.1572 | No |
| 46-64 | $0.112 \pm 0.137$ | $0.022 \pm 0.035$ | 640.0 | 0.0003 | Yes* |
| Overall | $0.087 \pm 0.131$ | $0.079 \pm 0.185$ | 3164.0 | 0.0010 | Yes* |

**Table S12: Comparison of Bursting Dynamics Between DF and VF Organoids, related to Figure 9.**
**Kolmogorov-Smirnov test, * Significant at p < 0.05. SEM = Standard Error of Mean**

| Measure | DF Mean (SEM) | VF Mean (SEM) | p-value | Sig | Effect Size |
|---|---|---|---|---|---|
| Burst Correlation | $0.239 \pm 0.017$ | $0.191 \pm 0.014$ | $0.001248$ | Yes* | $0.249$ |
| Timing Variability (ms) | $95.2 \pm 0.9$ | $94.0 \pm 1.4$ | $0.018706$ | Yes* | $0.021$ |
